# Chromosome-scale *Solanum pennellii* and *Solanum cheesmaniae* genome assemblies reveal structural variants, repeat content and recombination barriers of the tomato clade

**DOI:** 10.64898/2025.12.19.695418

**Authors:** Willem M. J. van Rengs, Roven Rommel Fuentes, Zahra Zangishei, Elias Primetis, Yazhong Wang, Joiselle Fernandes, Tamara Susanto, Qichao Lian, Sieglinde Effgen, Bruno Huettel, Saleh Alseekh, Björn Usadel, Charles J. Underwood

**Affiliations:** Department of Chromosome Biology, Max Planck Institute for Plant Breeding Research, Carl-von-Linné-Weg 10, 50829, Cologne, Germany; Faculty of Mathematics and Natural Sciences, Institute of Biological Data Science, CEPLAS, Heinrich Heine University Düsseldorf, Düsseldorf, Germany; Department of Plant and Animal Biology, Radboud Institute for Biological and Environmental Sciences, Radboud University, Nijmegen, the Netherlands; Max Planck Genome-center, Max Planck Institute for Plant Breeding Research, Carl-von-Linné-Weg 10, 50829, Cologne, Germany; Max-Planck-Institute of Molecular Plant Physiology, Am Mühlenberg 1, 14476, Potsdam-Golm, Germany; Institute of Bio- and Geosciences IBG-4, Bioinformatics, BioSC, CEPLAS, Forschungszentrum Jülich, Jülich, Germany; Keygene N.V., Wageningen, The Netherlands

**Keywords:** *Solanum lycopersicum*, *Solanum pennellii*, *Solanum cheesmaniae*, Structural variation, Transposable elements, Meiotic recombination, Linkage drag, Plant Breeding

## Abstract

Crop wild relatives are important resources for improving cultivated crops, yet the precise introgression of wild genetic material into cultivated crops is often difficult and cannot be fully defined without high-quality genome assemblies. Here we used PacBio HiFi, ONT ultra-long and Hi-C sequencing approaches to generate chromosome-scale *de novo* genome assemblies of two wild species related to the domesticated tomato (*Solanum lycopersicum*) — the broadly stress-resistant *Solanum pennellii* (accession LA0716) and the salt-resistant *Solanum cheesmaniae* (accession LA1039). K-mer and BUSCO analysis of both assemblies demonstrated above 99% completeness and the improved *S. pennellii* genome adds 146 Mbp to the 12 chromosomes compared with the original reference. We aligned the new assemblies with seven gold-standard assemblies from the *Lycopersicon* clade, using *Solanum tuberosum* as an outgroup, and identified shared and species-specific structural variants. The repeat content of all nine assemblies was characterized, providing evidence for independent explosions of *Tekay* (gypsy superfamily) retrotransposons in *S. pennellii* and *S. peruvianum*. Whole genome sequencing of 709 recombinant plants derived from male and female backcrosses of three different hybrids (*S.pennellii*, *S. cheesemaniae* and *Solanum lycopersicum* cv. Micro-Tom crossed with *Solanum lycopersicum* cv. Moneyberg-TMV) revealed higher crossover rate in female meiosis. Recombination landscape analysis identified conserved female-enhanced recombination regions, and coldspots that were completely devoid of meiotic crossovers including megabase-scale inversions and insertion-deletion polymorphisms between *S. lycopersicum* and *S. pennellii*. In summary, we harnessed our high-quality *S. pennellii* and *S. cheesmaniae* genome assemblies to reveal how repeat content diverged in nature and during breeding, and uncovered how reproductive gender interacts with structural variants to dictate the recombination landscape in tomato hybrids.

## Introduction

Modern molecular plant breeding harnesses near-complete genome sequences to reveal gene markers that control traits of agronomic importance. The cultivated tomato is the world’s number one vegetable crop and recent tomato improvement was catalyzed by the release of the first genome assembly of the *Solanum lycopersicum* variety Heinz1706 in 2012^1,2^. Furthermore, a combination of short-read sequencing, BAC sequencing and genetic map information resulted in a genome assembly of the stress-resistant distant tomato relative *Solanum pennellii* (accession LA0716)^3^. This assembly has been an important resource to elaborate on the genetic basis of important agronomic traits through the well-used *S. lycopersicum* cv. M82 x *S. pennellii* LA0716 recombinant inbred line population^4^. Although high-throughput short-read sequencing accelerated the discovery of natural genetic variants, it has also introduced an unavoidable bias: characterized variants are disproportionately skewed toward single-nucleotide polymorphisms (SNPs) and small indels^5,6^. Furthermore, short reads cannot accurately resolve complex genomic regions, repetitive sequences, and large structural variations, leading to fragmented genome assemblies that lack essential genomic context. Compared to Heinz1706 that has been re-assembled several times using newer technologies including long-read sequencing and chromatin conformation map^2,7–9^, the *S. pennellii* LA0716 genome remained as it was, notwithstanding efforts to use long read sequencing for the *S. pennellii* accession LA5240 which is used for a high resolution genetic mapping^10,11^. Despite the release of superpangenomes for the *Solanum* genus and the *Lycopersicon* clade^12,13^ the absence of an improved *S. pennellii* LA0716 genome and an assembly of the stress-resistant *Solanum cheesmaniae* (accession LA1039)^14^ hinders introgression efforts from these important sources of abiotic stress tolerance.

High-quality genomes enable the exploration of large and complex genetic variants, including structural variants (SVs) and transposable elements (TEs). Typically, SVs impose greater potential impact to gene structure, dosage and location than simple genetic variants, and SVs contribute to species evolution and diversity^12,13,15–20^. Recent studies on SVs have indicated they can be causal genetic variants for agronomically-relevant traits that are not accounted by simple variants like single nucleotide polymorphisms (SNPs) and small insertions or deletions^8,19,21^. For example, in the domesticated tomato an 83 kbp duplication at the *supressor of branching 3* locus was co-selected with a *jointless2* allele that improves fruit harvestibility^21^. Similar to SVs, TEs are among the major drivers of genome evolution and diversity in plants^22^, often comprising the majority of genomic content in crop species such as maize and wheat^23,24^. Retrotransposons (class I TEs) are often activated under biotic and abiotic stress conditions, and developmental stages, contributing to genome plasticity and potentially facilitating plant adaptation and evolution^25–29^. In the *Solanaceae* family, the majority of Long Terminal Repeat (LTR) retrotransposons are from the *Ty3* (Gypsy) superfamily^12^, and in the *Solanum* genus the *Tekay* (*Ty3*) and *Tork* (*Ty1*) lineages are most abundant^30,31^. The *Tekay* lineage is also a prevalently active LTR retrotransposon lineage in several other plant families^32–34^. It was previously estimated that repeats make up ∼82% of the *S. pennellii* genome, 45% of which consist of retrotransposons^3^.

Apart from genetic polymorphisms, investigating meiotic recombination or crossovers (COs) in the genome is essential for the precise introgression of alleles from wild relatives to domesticated tomato. COs are not distributed uniformly across plant genomes but occur in so-called hotspots often distributed in distal regions of the chromosomes^35–37^. This non-uniform crossover distribution limits the shuffling of alleles in regions of the genome that are devoid of COs, resulting in linkage between advantageous and negative traits (so-called linkage drag) and thereby inefficiency in breeding. CO hotspots are associated with gene-rich areas, whereas coldspots are observed in retrotransposon-rich, heterochromatic or inverted regions^38–41^. Studies in different species have linked recombination suppression with structural variations^42–45^, but it was only recently that the causal proof was provided by reverting an inversion through genome editing^46^. A relatively underexplored topic in meiotic CO patterning is the differential rate of meiotic recombination between male and female gametes, known as heterochiasmy. Unlike in *Arabidopsis thaliana*, where meiotic COs are more frequent in male meiosis than female meiosis^47–50^, there is evidence from genetic marker studies in a *S. lycopersicum* x *S. pennellii* hybrid that female recombination rate is higher than male^51^. Exploring tomato heterochiasmy in finer resolution and in multiple genetic backgrounds has yet to be carried out. Whole genome sequencing of bi-directional backcross populations could now enable not just the detection and comparison of crossovers in different hybrid crosses, but the identification of possible female-enriched crossover hotspots.

In this study, we generated a high-quality long-read based reference genome of *S. pennellii* LA0716 and the first assembly of *S. cheesmaniae* LA1039. We identified genomic variants including megabase-scale SVs and transposable elements expansions between each genome and five cultivated tomato accessions (Heinz1706 SL5.0, Moneyberg-TMV, Micro-Tom, M82, Sweet100), *Solanum galapagense*, *Solanum habrochaites*, and *Solanum tuberosum*^8,52–56^ (Supplementary Table 1). Moreover, we explored the crossover distribution and heterochiasmy in tomato using re-sequencing data from six large backcross populations derived from three hybrids with variable divergence at the DNA sequence level. This new resource provides a compendium of variation between cultivated and wild tomatoes, and could enable the genome-guided introgression of wild traits to the cultivated tomato during the development of stress-and disease-resistant varieties.

## Results

### Chromosome-scale *Solanum pennellii* and *Solanum cheesmaniae* genome assemblies

We set out to assemble high-quality chromosome scale genome assemblies of *Solanum cheesmaniae* accession LA1039 and *Solanum pennellii* accession LA0716. To this end we generated PacBio HiFi data using two different input sizes per genotype leading to datasets of 51.4 Gbp (*S. cheesmaniae* LA1039) and 49 Gbp (*S. pennellii* LA0716) (Supplementary Table 2). K-mer (k=21) analysis of HiFi data revealed only a single clear peak for both accessions (Supplementary Fig. 1), suggesting all accessions were highly homozygous. Due to the known higher repeat content of *S. pennellii*^3^, we additionally generated 381.2 Gbp of ONT data (Supplementary Table 3) and filtered for high-quality reads that were longer than 90 kbp, resulting in 16.08 Gbp of high-quality long reads. We also generated chromosome confirmation capture data (Omni-C (Dovetail)) to provide datasets for reference-free scaffolding^57,58^, including 41 Gbp for *S. cheesmaniae* LA1039 and 56 Gbp for *S. pennellii* LA0716 (Supplementary Table 4).

Due to the high homozygosity of both species, we assembled them as single genome haplotypes using Hifiasm^59^. This resulted in assemblies totaling 862 Mbp (*S. cheesmaniae*) – including the direct assembly of the complete *S. cheesmaniae* chromosome 6 - and 1109 Mbp (*S. pennellii*) with respective N50 values 27.9 Mbp and 26.2 Mbp (Supplementary Table 5). Next, we used the Omni-C data for automated scaffolding of the raw assembled contigs. Contiguity of our *de novo S. cheesmaniae* and *S. pennellii* genome assemblies was further improved through multiple rounds of manual scaffolding and scaffolding correction, until chromosome-scale genomes were obtained (Fig 1a-b and Table 1). The final assemblies were supported by chromatin confirmation capture as inter-chromosomal interaction was only observed between highly similar telomeric sequences, and none of the sequences were broken by re-running automated scaffolding (Fig. 1a-b). We observed highly co-linear alignments of the twelve assembled *S. cheesmaniae* chromosomes to *S. lycopersicum* cv. Moneyberg-TMV (hereafter MbTMV) chromosomes 1-12, whereas the chromosome set of the divergent relative *S. pennellii* showed more structural variations, as expected (Fig. 1c-d). Unplaced *S. cheesmaniae* and *S. pennellii* sequences did not show any further Hi-C interaction with any of the assembled chromosomes and were therefore not placed into the chromosomes (Fig. 1a-b).

**Fig. 1.**
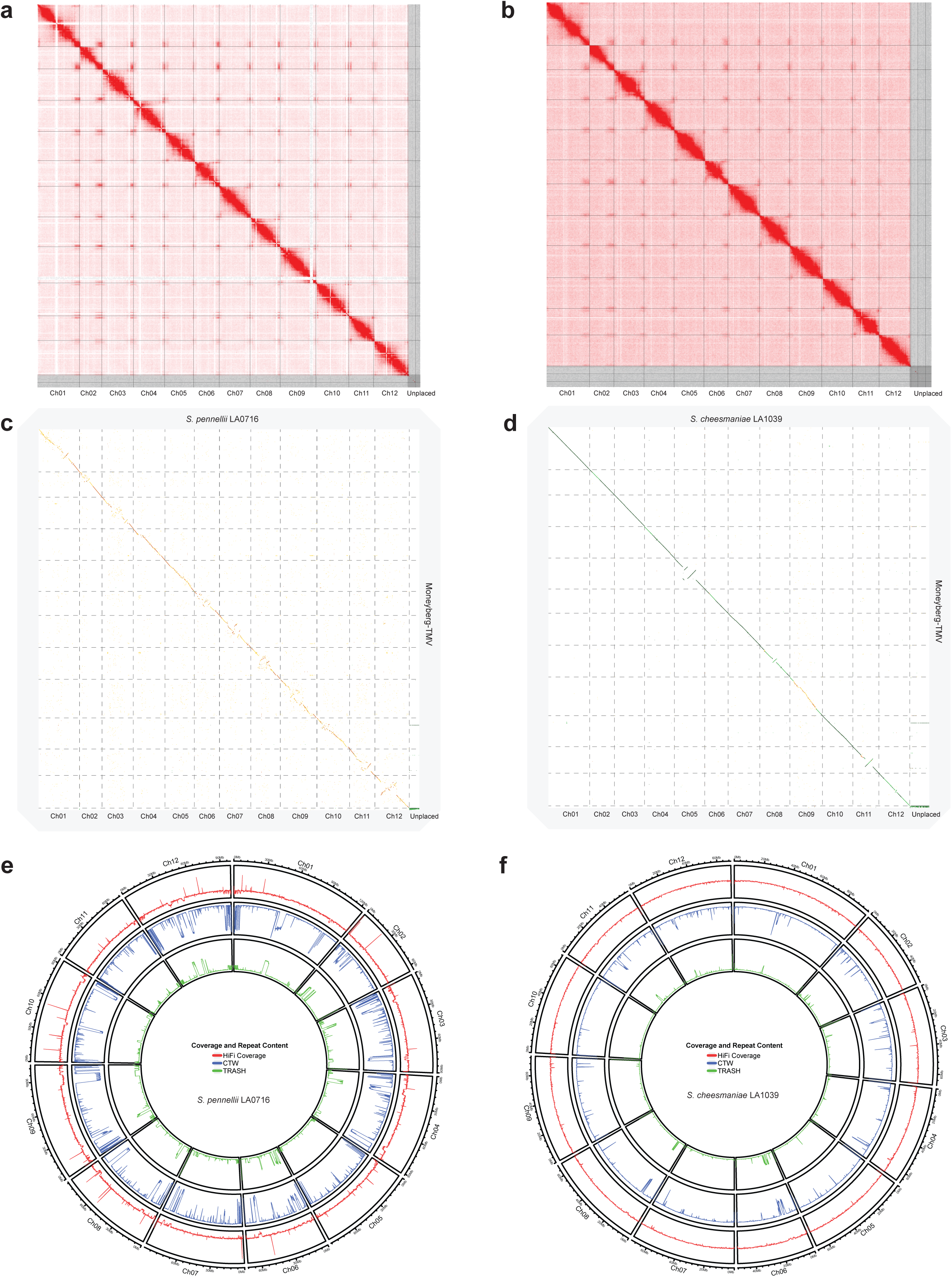
Chromosome-scale *Solanum pennellii* and *Solanum cheesmaniae* genome assemblies. **a** *S. pennellii* LA0716 Hi-C interaction plot. **b** *S. cheesmaniae* LA1039 Hi-C interaction plot. Hi-C interaction plotted by Juicebox, demonstrating strong interactions over the 12 assembled chromosomes of the *S. pennellii* (**a**) and *S. cheesmaniae* (**b**) genome assemblies. Grey lines depict sequence borders. **c** Whole genome comparison dotplot of *S. pennellii* (top axis) aligned to *S. lycopersicum* cv. Moneyberg-TMV (right axis). **d** Whole genome dotplot of *S. cheesmaniae* (top axis) aligned to *S. lycopersicum* cv. Moneyberg-TMV (right axis). Horizontal grey dashed lines represent Moneyberg-TMV chromosome borders. Vertical grey lines represent *S. pennellii* (**c**) and *S. cheesmaniae* (**d**) chromosome borders. Dotplots visualizations are created by D-genies (Cabanettes and Klop, 2018). Genome coverage and repeat content for *S. pennellii* (**e**) and *S. cheesmaniae* (**f**). From outer to inner layer (in 100 Kbp windows): logarithmically normalised and scaled HiFi genome coverage (red line), the log-probability of the CTW algorithm (Kontoyiannis et al., 2022) (blue line) and the abundance of monomeric repeats using TRASH (Wlodzimierz et al., 2023) (green line) for *S.pennelli* (**e**) and *S.cheesmaniae* (**f**).

**Table 1:** Genome assembly statistics of *S. pennellii* and *S. cheesmaniae* compared to published assemblies. Table 1 shows the summary statistics of the 12 chromosome scale sequences making up each genome assembly. BUSCO analysis of the completeness of gene content is using the Solanales benchmark set. QV and completeness are estimated by K-mer based analysis using HiFi data only.

| Accession | Moneyberg-TMV | Micro-Tom | LA1039 | LA0716 | LA2093 | LA0716 |
| --- | --- | --- | --- | --- | --- | --- |
| Species | *S. lycopersicum* | *S. lycopersicum* | *S. cheesmaniae* | *S. pennellii* | *S. pimpinellifolium* | *S. pennellii* |
| Reference | van Rengs et al., 2022 | Wang et al., 2024 | This study | This study | Wang et al., 2020 | Bolger et al., 2014 |
| Number of sequences | 12 | 12 | 12 | 12 | 12 | 12 |
| Number of sequences (>50kb) | 12 | 12 | 12 | 12 | 12 | 12 |
| Cumulative size (Mbp) | 824.45 | 812.44 | 814.47 | 1072.33 | 800.50 | 926.43 |
| N50 (Mbp) | 68.5 | 67.8 | 68.2 | 90.2 | 67.2 | 78 |
| N90 (Mbp) | 54.7 | 56.8 | 55.8 | 72.3 | 55.6 | 60.7 |
| L50 | 6 | 6 | 6 | 6 | 6 | 6 |
| L90 | 11 | 11 | 11 | 11 | 11 | 11 |
| Longest sequence (Mbp) | 96.5 | 96.3 | 96.7 | 123.5 | 95.5 | 109.3 |
| Number of N's | 0 | 4000 | 3300 | 4400 | 487215 | 71074709 |
| Number of internal N-regions | 0 | 40 | 33 | 44 | 373 | 41684 |
| Raw LAI | 9.67 | 8.42 | 8.64 | 12.43 | 8.49 | 3.24 |
| LAI | 15.3 | 14.05 | 14.27 | 18.06 | 14.12 | 8.87 |
| Complete BUSCO (C) | 5852 | 5851 | 5852 | 5857 | 5851 | 5827 |
| Complete Single copy (S) | 5747 | 5743 | 5741 | 5738 | 5744 | 5706 |
| Complete Duplicated (D) | 105 | 108 | 111 | 119 | 107 | 121 |
| Fragmented (F) | 12 | 12 | 12 | 14 | 12 | 15 |
| Missing (M) | 86 | 87 | 86 | 79 | 87 | 108 |
| Total searched (Solanales) | 5950 | 5950 | 5950 | 5950 | 5950 | 5950 |
| QV | 54.12 | 72.40 | 75.69 | 69.97 | N.A. | N.A. |
| Completeness | 99.10 | 99.22 | 99.17 | 99.17 | N.A. | N.A. |

To assess the quality of our chromosome-scale genome assemblies, we compared statistics of the assembled *S. cheesmaniae* and *S. pennellii* pseudochromosomes to the 12 pseudochromosomes of reference quality tomato assemblies including Moneyberg-TMV^53^, the original *S. pennellii* assembly^3^ and an *S. pimpinellifolium* assembly^60^ (Table 1). We found our scaffolded assemblies performed favorably in terms of contiguity for *S. cheesmaniae* (N50=68.2 Mbp, N90=55.8 Mbp) and *S. pennellii* (N50=90.2 Mbp, N90=72.3 Mbp) and repeat content assembly (both with LAI above 14)^61^, had few internal gap regions and performed similar in terms of gene completeness (BUSCO)^62^ metrics (Table 1). K-mer based assembly evaluation using Merqury^63^ showed completeness above 99% for the *S. cheesmaniae* and *S. pennellii* assemblies (QVs above 69) (Table 1), indicating the assemblies were of high-quality.

The genome assemblies were further validated by means of read coverage analysis and characterization of nucleotide composition and repeat elements including Simple Sequence Repeats (SSRs) and LTRs (Fig. 1e-f, Supplementary Fig. 2 and 3). After re-mapping the long reads, we found mostly even coverage along the chromosomes with a few outlier regions. For both genomes, multiple windows with reduced coverage often overlaid drastic changes in nucleotide composition and reduced SSR and LTR densities (Fig. 1e-f, Supplementary Fig. 2 and 3). We previously found such a behavior on the MbTMV chromosome 8, where these changes correspond to irregular genomic features including telomeres, 45S rDNA, and sub-telomeric Tomato Genome Repeat I (TGR1) sequences^53^, suggesting a similar phenomenon may occur in the *S. cheesmaniae* and *S. pennellii* assemblies. Notably, these genomic features are extensively present in our *S. pennellii* assembly (Supplementary Fig. 3) and could be more completely assembled due to utilization of >90kb ONT reads.

**Fig. 2.**
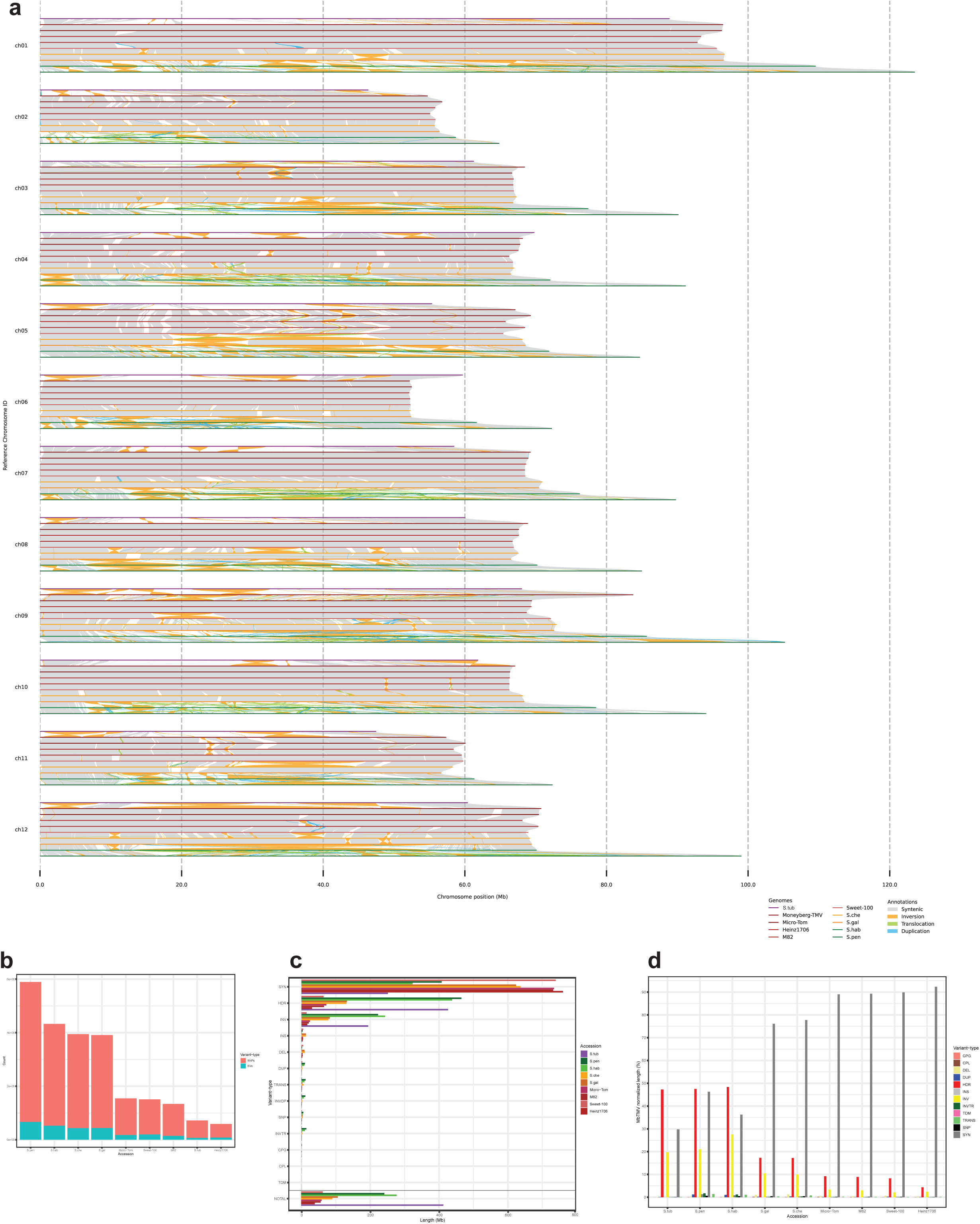
Whole genome alignment of nine high-quality Lycopersicon clade genomes reveals shared and species-specific structural variants. **a** Pan-genome visualization of 10 solanum genomes, including 9 species of the tomato clade (*Solanum sect. Lycopersicon*) and 1 potato (*S. tuberosum*). Visualization was generated with SyRI and plotsr. *Solanum sect. Lycopersicon* genomes include 5 *S. lycopersicum* species (Heinz1706, Moneyberg-TMV, Micro-Tom, M82, Sweet-100), *S. cheesmaniae* LA1039 (S.che), *S. galapagense* LA0317 (S.gal), *S. habrochaites* LA0407 (S.hab), *S. pennellii* LA0716 (S.pen). *S. tuberosum* DM8 (S.tub) is included as outgroup. **b** Total number of SNPs and Structural variants (SVs) identified per genome used in figure a and compared to the Moneyberg-TMV reference genome. SNP numbers exceed SV numbers in all lines. All except the following SVs are included: DUPAL, INVAL, INVDPAL and TRANSAL. **c** Summary of variant lengths per genome, based on the SyRi annotation blocks in comparison to the Moneyberg-TMV reference genome. Besides syntenic regions, Highly diverse regions (HDR) and inversions (INV) makeup the largest total length of SVs. A degree of overlap in variant length can be expected due to genome size differences and variant definitions, e.g. regions containing duplications (DUP & INVDP) and HDRs. Not aligned regions (NOTAL) are inferred from regions not being part of any annotation blocks. **d** Lenght-wise genomic composition of each line compared to the Moneyberg-TMV reference genome. A decline of synthenic regions (SYN) is clearly visible following the speciation within all 10 solanum genomes, where the *S.lycopersicum* species show highest syntheny.

**Fig. 3.**
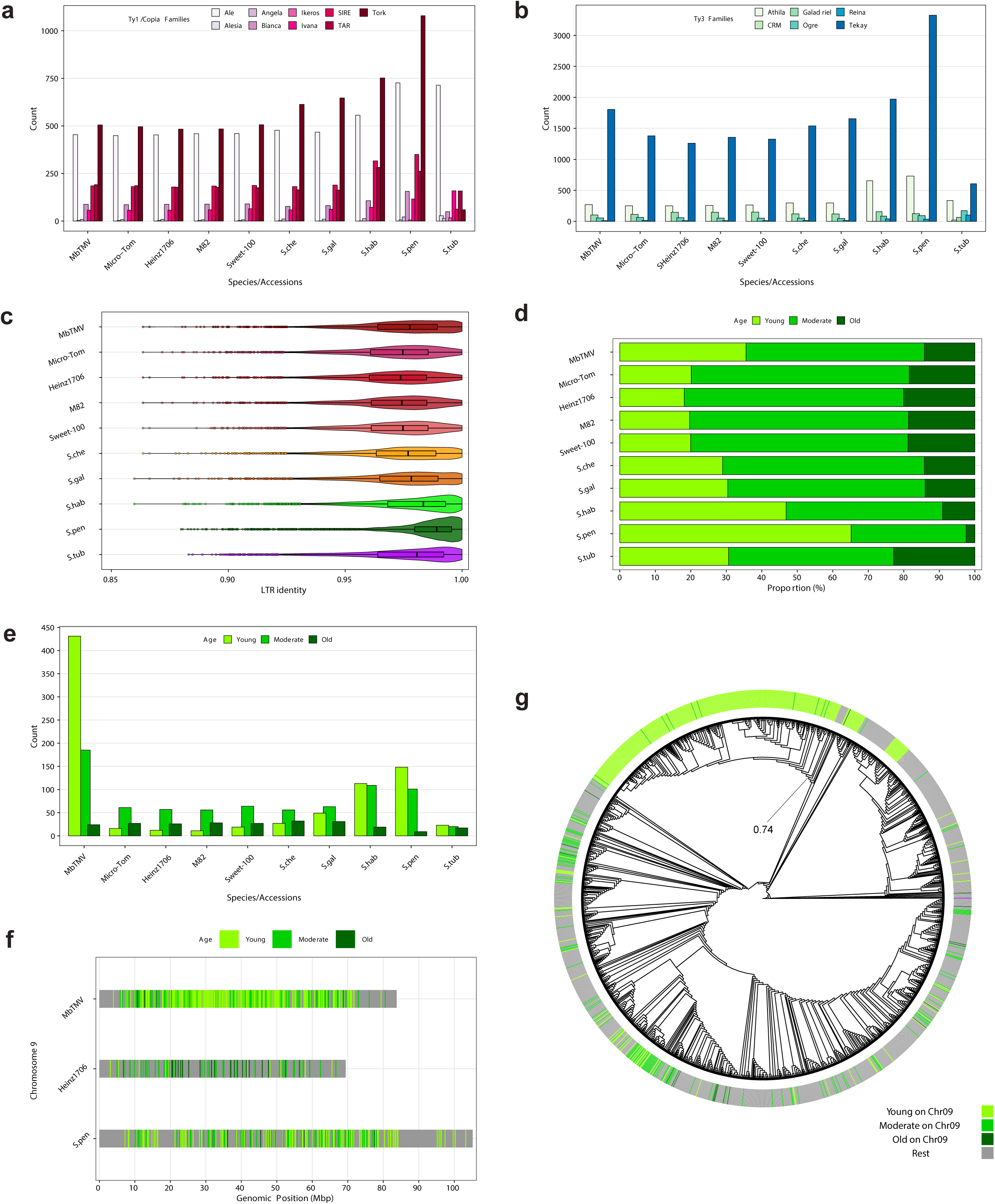
Transposable element annotation of the *Lycopersicon* clade reveals independent explosions of Tekay retrotransposons. **a** and **b** Quantification of the intact elements in the main *Ty1/Copia* and *Ty3* lineages. **c** Age of all the LTR retrotransposons (*Ty1/Copia* and *Ty3*) in all the species using the LTR identity metric. **d** The proportion of the *Tekay* elements distinguished in the different age categories (young <0.5 mya, 0.5 >= moderate <= 1.5 mya and old >1.5 mya) in all the species. **e** The abundance of the different age categories of *Tekay* elements on chromosome 9 of each species. **f** The distribution of the *Tekay* elements on chromosome 9 of *S. lycopersicum* cv. Moneyberg-TMV (MbTMV), *S. lycopersicum* cv Heinz1706 (Heinz1706) and *S. pennellii* LA0716 (S.pen). Different colours represent the age category of each element. **g** Phylogenetic tree of the high-quality integrases of the *Tekay* elements. The heatmap on the outer layer shows the age of the element. The tree is rooted using a *Ty1/Copia Tork* element from S.tub (purple block on the right of the heatmap). The bootstrap value for the main cluster of young elements is shown.

The short-read based assembly of *S. pennellii* LA0716 has been an important platform for research into genetically encoded abiotic and biotic stress resistance in this wild relative of the cultivated tomato^3^. We performed an extensive comparison of the original and new *S. pennellii* genome assemblies and found a total increase in assembly size of 145.9 Mbp between the 12 assembled chromosomes (Supplementary Fig. 4 and Supplementary Table 6). The new *S. pennellii* assembly had higher gene completeness based on BUSCO analysis^62^, the repeat content was more completely assembled based on LAI analysis^61^ and more than 41,000 gaps were filled (Table 1). Whole genome alignment and pairwise comparison demonstrated highly syntenic regions especially in the distal ends of each chromosome. However, we also found several large differences including inversions (Chromosomes 1, 2, 3, 6, 7, 8) between the two assemblies (Supplementary Fig. 4 and 5) and expect that the absence of recombination could have hindered genetic map-based scaffolding in the original assembly. Re-mapping of raw sequencing reads to both assemblies indicated that the new assembly has a more even read coverage across the chromosomes compared to the short-read based assembly (Supplementary Fig. 5).

**Fig. 4.**
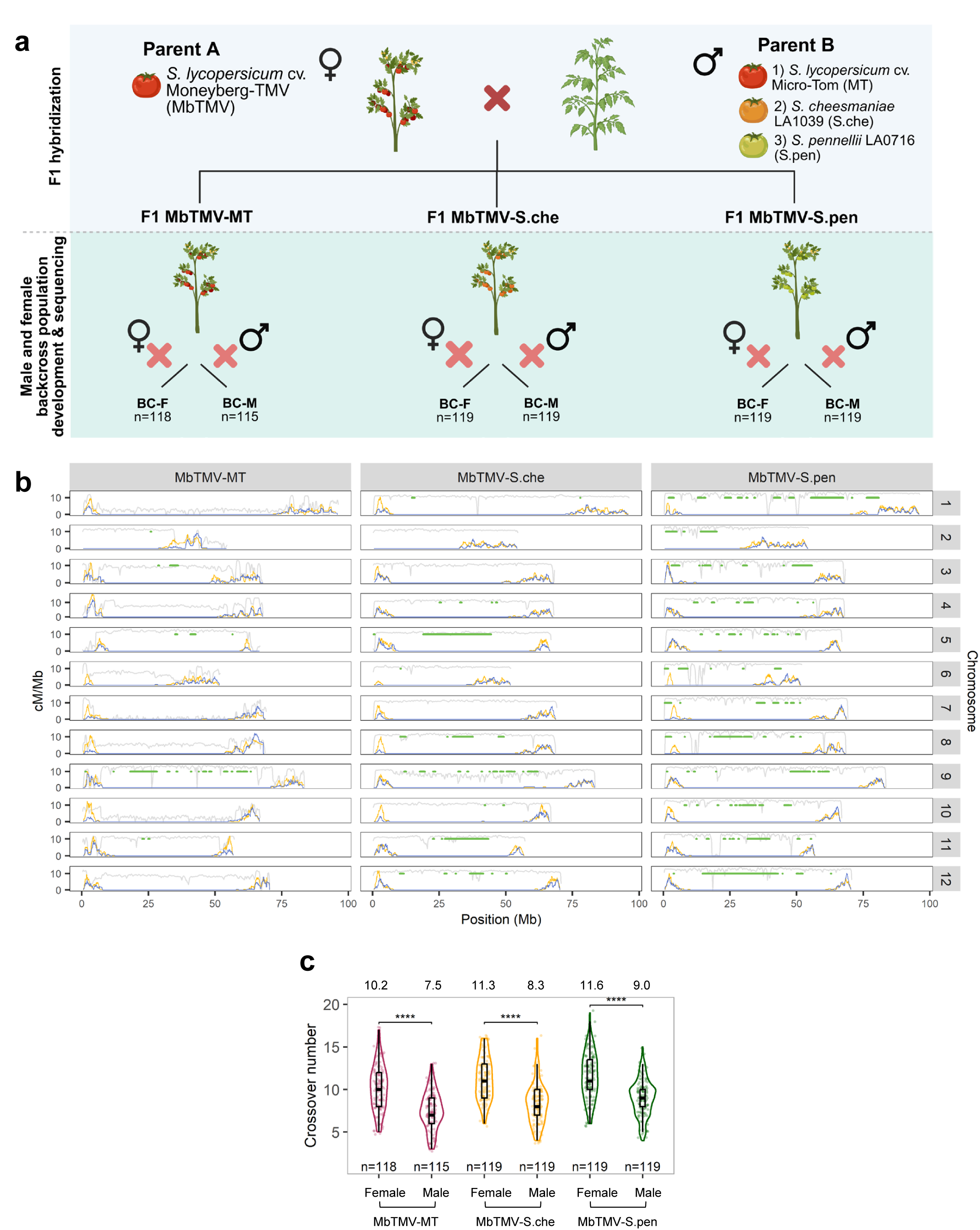
Higher crossover rate in female meiosis underlies female-enhanced recombinaton regions in tomato intraspecific and interspecific hybrid backcross populations. **a** Generation of female and male backcross populations for genome sequencing. Created in BioRender. van Rengs, W. (https://BioRender.com/fz42vck). **b** Recombination landscape in female (yellow) and male (blue) backcross populations. Gray lines indicate the distribution of SNP (log scaled), while green horizontal lines show the locations of large inversions that are larger than 100 kbp. Some distal regions in MbTMV-MT such as chromosome 2 and 5 have low SNP density, limiting crossover detection. **c** Female backcross populations have significantly higher number of crossovers than male populations (Wilcoxon Rank Sum Test; ****: <10-3). The mean crossover number per population is specified on top of the graph.

**Fig. 5.**
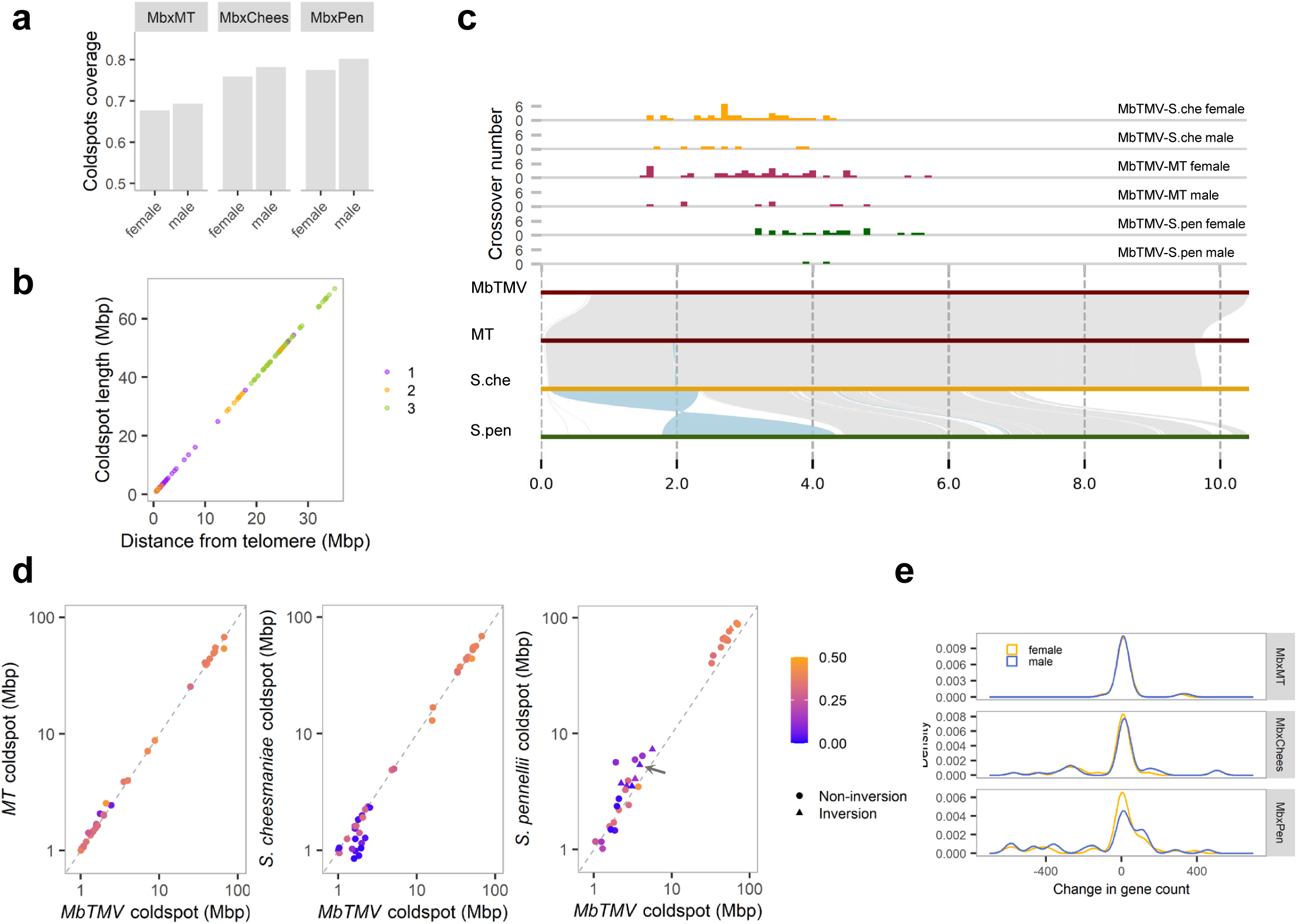
Novel retrotransposon expansions and genome inversions associated with recombination coldspots in a divergent tomato hybrid. **a** Proportion of coldspots in different hybrid crosses expressed as a percentage of the MbTMV genome. **b** Relative position of cross-specific and conserved coldspots across the three hybrid genotypes. The color indicates the number of hybrid genotypes where the coldspot is observed. **c** A 2.2 Mbp inversion suppresses recombination between MbTMV and S. pennellii chromosome 8 short arm, whereas other hybrid populations exhibit recombination in the same region. **d** Size of male coldspot regions relative to each parental genome. Deviation from the diagonal line indicates expansion or contraction of genomic segments, while the color indicates the total proportion of Gyspy and Copia elements relative to the MbTMV genome. We define inversion-associated coldspot as having more than 50% inversion coverage. Orange dots in the upper right of each plot represent large coldspots in the pericentromeres. The arrow points to the chromosome 8 coldspot illustrated in **b**. **e** Change in coldspot gene counts between the parental genomes (Δ gene count = MbTMV – Parent 2). Cross-specific expansions and translocations in the chromosome arms contributed to the changes in the distributions in MbTMV-S.pen.

Next, we examined the distribution of repeats across the genomes of *S. cheesmaniae* and *S. pennellii* using complementary CTW (Context Tree Weighting)^64^ and TRASH (Tandem Repeat Annotation and Structural Hierarchy)^65^ analysis (Fig. 1e-f). In general, both tools identify more extensive repetitive regions in *S. pennelli* than *S. cheesmaniae* or MbTMV (Fig. 1e-f and Supplementary Fig. 6). We find that in *S. pennelli* repetitive regions are much more broadly distributed along the chromosomes than in *S. cheesmaniae* and MbTMV where punctate repetitive arrays are more common (Fig. 1e-f and Supplementary Fig. 6). The highly repetitive 5S rDNA (Chromosome 1) and tandem arrays of 45S rDNA derived satellite (Chromosomes 6, 8 and 11) were identified in all three assemblies, while other repetitive regions of unknown origin were found by both tools on *S. pennellii* chromosomes 2, 3, 4, 9 and 12 (Fig. 1e-f and Supplementary Fig. 6). In most chromosomes the repetitive content of *S. cheesmaniae* and MbTMV was very similar, indicative of the closer evolutionary relationship of the two genomes compared to the more distant *S. pennelli* (Fig. 1e-f and Supplementary Fig. 6). Interestingly, some repetitive regions were not identified by both tools (Fig. 1e-f), reflecting the different methodological (mathematical and biological) approaches. In summary, our findings suggest that the *S. pennelli* genome has a more complex repetitive content than the genomes of *S. cheesmaniae* and the domesticated tomato.

### Whole genome alignment of nine high-quality *Lycopersicon* clade genomes reveals shared and species-specific structural variants

To analyze the structural variants of the new *S. cheesmaniae* and *S. pennellii* genome assemblies in the context of the complete *Lycopersicon* clade, we included eight additional high-quality genome assemblies that were constructed with at least PacBio HiFi and Hi-C data. These assemblies included five cultivated tomato accessions (Heinz1706 SL5.0, MbTMV, Micro-Tom, M82, Sweet-100), two additional wild tomato species (*S. galapagense* LA0317 and *S. habrochaites* LA0407), and *S. tuberosum* (DM8.1) as outgroup (Supplementary Table 1). We identified both small and large genomic variants, between our reference model accession *S. lycopersicum* cv. Moneyberg-TMV and the other genomes through whole genome pairwise comparison (Fig. 2, Supplementary Fig. 7 and 8).

The five *S. lycopersicum* genomes show a high degree of genome-wide synteny, making up more than 700 Mbp of their complete genome sizes (Fig. 2a,c). Comparing *S. lycopersicum* MbTMV with the four other cultivated tomato accessions identified between 506,477-1,366,260 SNPs, and 74,706-179,406 SVs (Fig. 2a-c). Notably, the distal, gene-rich, ends of all twelve chromosomes are fully syntenic in the five *S. lycopersicum* genomes apart from rare inversions on the first arms of chromosomes 9 (Micro-Tom, Heinz1706 and M82) and 12 (Micro-Tom and Sweet100) (Fig. 2a). Moreover, the complete chromosomes 6, 7 and 8, including the vast pericentromeric regions, are structurally almost identical in all five cultivated tomatoes (Fig. 2a). In the other nine chromosomes, different pericentromeric haplotypes were identified in specific accessions (Fig 2a). For example, on chromosome 1 both Heinz1706 and M82 share a shorter pericentromeric haplotype compared to the other three accessions, while on chromosome 3 Micro-Tom contains two accession-specific large inversions in the pericentromeric region (Fig 2a). Alternative pericentromeric haplotypes were also found on chromosomes 2 (Micro-Tom), 4 (M82 and Sweet100), 5 (Micro-Tom and M82), 9 (MbTMV and Sweet100), 10 (M82), 11 (MbTMV and Heinz1706) and 12 (Heinz1706 and Sweet100) compared to the other cultivated tomato species. Overall, the structural diversity in the cultivated tomato genomes is largely restricted to the pericentromeric heterochromatic regions.

The structural diversity between the five *S. lycopersicum* genomes can be considered modest in the context of the genomes of the four wild relatives (*S. galapagense*, *S. cheesmaniae*, *S. habrochaites* and *S. pennellii*) and outgroup (*S. tuberosum*). The independently assembled Galapagos tomato (*S. galapagense* and *S. cheesmaniae*) genomes show a high degree of synteny genome-wide sharing large inversions on chromosomes 5 (18-21 Mbp), 8 (7.4 Mbp) and 11 (16-17 Mbp) compared to the five *S. lycopersicum* accessions (Fig. 2a). Although over 600 Mbp of the Galapagos tomato genomes are syntenic to *S. lycopersicum,* a two-fold increase in the total number of variants was detected with respect to SNPs (3,473,617-3,510,695) and SVs (435,906-439,402) between *S. lycopersicum* and the two Galapagos tomato (Fig. 2b-c). Despite the similarities between the *S. cheesmaniae* and *S. galapagense* genomes, large inversions (greater than 2 Mbp) on chromosomes 1, 3, 5 and 12 (Fig. 2a), and differences in the total variant lengths (Fig. 2c), demonstrate they are divergent from one another.

Next we explored simple and structural divergence with the more divergent “Hirsutum” group species *S. habrochaites* and *S. pennellii*, and the outgroup *S. tuberosum*. Between *S. lycopersicum* and *S. habrochaites*/*S. pennellii*/*S. tuberosum* we identified SNPs (3,805,503/5,230,709/649,002) and SVs (536,721/ 682,778/ 74,706), with the lower number of variants with *S. tuberosum* being explained due to only 29.8% of the sequence being syntenic (Fig 2b-d). Consistent with previous reports we found complete inversions on the chromosome arms between *S. lycopersicum* and *S. pennellii* (first arms of chromosomes six and seven), and between *S. lycopersicum* and *S. tuberosum* (one arm inversion on chromosomes two, five, six, eight, nine, ten, eleven and twelve)^66–68^. Our analysis led to the discovery of two large unreported inversions on the chromosome arms. We found a 4.7 Mbp inversion between *S. lycopersicum* and *S. habrochaites* (on the first arm of chromosome 4) and a 2.2 Mbp inversion of the majority of first arm of chromosome 8 between *S. lycopersicum* and *S. pennellii* (Fig. 2a). The former inversion may be a technical artifact due to the presence of an assembly gap 11 kbp from the proximal inversion breakpoint, whereas the latter inversion between *S. lycopersicum* and *S. pennellii* could be validated by detailed Hi-C analysis (Supplementary Fig. 16). In total, we found 202 inversions between the *S. lycopersicum* and *S. pennellii* genomes making up over 200 Mbp in length (Fig. 2c). In addition, the new *S. pennellii* assembly demonstrates the presence of a 7.1 Mbp inversion on chromosome 3 that was previously reported between *S. lycopersicum* and many other wild relatives yet was thought to be absent in *S. pennellii*^3,13^(Supplementary Fig. 4). In general, the structure of the pericentromeric regions differs greatly between *S. habrochaites* and *S. pennellii* and yet a 300 kbp inversion common to the two “Hirsutum” group species was identified on the first arm of chromosome 9 (Fig 2a) and is evidence of their close phylogenetic relationship. Finally, our qualitative assessment was validated by the normalization of total variant lengths compared to the MbTMV genome which demonstrated low synteny with *S. pennelllii* (46.3%) and *S. habrochaites* (36.2%) (Fig. 2d). In conclusion, high-quality genomes derived from long read sequencing and Hi-C data allowed us to uncover novel examples of genomic rearrangements in heterochromatic pericentromeric regions and euchromatic chromosome arms between related *Solanum* species.

### Transposable element annotation of the *Lycopersicon* clade reveals independent explosions of *Tekay* retrotransposons

To elucidate the TE content of the *Lycopersicon* clade, we conducted a detailed annotation retrotransposons (class I TEs) and DNA transposons (class II TEs) of the aforementioned nine tomato genomes and single potato genome (Fig. 3, Supplementary Fig. 9-10, Supplementary Table 1, Table 2). Globally we found that the total TE content tends to be higher in wild tomato species compared to cultivated tomatoes, with nearly 65% of the *S. pennellii* genome composed of TEs, compared to just 54.74% of the *S. lycopersicum* MbTMV genome (Table 2). We noticed that the TE component is related to genome size, which is higher in non-domesticated accessions (Table 2). For example, the *S. pennellii* genome is 30.1% larger than the *S. lycopersicum* MbTMV genome (Table 2). We found that *Ty3* LTR retrotransposons are the most abundant TE superfamily across the ten species, representing 17.4% of the *S. galapagense* genome and up to 24.43% of the potato genome (Table 2). The genomic percentage of the *Ty1/Copia* Class I superfamily does not follow the same trend, with the highest percentage observed in *S. cheesmaniae* (6.26%) and the lowest in potato (5%) (Table 2). Other LTR retrotransposon classes account for 4.88-6.79% of the nine tomato genomes, whereas this grouping makes up 11.15% of the potato genome (Table 2). Class II TEs are less prominent in the studied genomes, where the *Mutator* family is most prevalent with its highest genomic percentage detected in *S. pennellii* (14.88%), while the lowest is found in potato (4.46%) (Table 2).

**Table 2:** Genomic percentage of the TE superfamilies in nine tomato accessions and the potato reference genome DM8.1. Table 2 shows the genomic percentage of the main TE superfamilies, including LTR retrotransposons and DNA transposons, in nine tomato accessions and the potato reference genome DM8.1. In addition, the total TE percentage and genome size in Mbp for each accession are presented. The TE annotation was performed using EDTA (v.2.0.1) (Ou et al., 2019).

| Species/TE superfamily | *Ty1/Copia* | *Ty3* | Unknown LTR | *CACTA* | *Mutator* | PIF/<br>Harbinger | *TC1/*<br>*Mariner* | *hAT* | *Helitron* | Total TE | Genome Size (Mbp) |
| --- | --- | --- | --- | --- | --- | --- | --- | --- | --- | --- | --- |
| ***S.lyc*-MbTMV** | 5.77 | 18.63 | 5.99 | 4.53 | 8.66 | 1.10 | 1.58 | 1.13 | 7.53 | 54.73 | 824.45 |
| ***S.lyc*-MicroTom** | 5.88 | 18.51 | 5.15 | 5.01 | 9.86 | 1.48 | 1.19 | 0.94 | 8.50 | 56.53 | 812.44 |
| ***S.lyc*-Heinz1706** | 6.16 | 18.98 | 4.88 | 4.30 | 8.85 | 1.74 | 1.48 | 1.01 | 8.19 | 55.59 | 800.12 |
| ***S.lyc*-M82** | 5.87 | 18.89 | 5.26 | 5.25 | 9.47 | 1.56 | 1.18 | 0.85 | 8.42 | 56.74 | 801.96 |
| ***S.lyc*-Sweet100** | 6.12 | 18.41 | 6.26 | 4.69 | 9.26 | 1.30 | 1.39 | 1.21 | 7.80 | 56.45 | 805.19 |
| ***S.che*** | 6.26 | 17.78 | 6.35 | 6.14 | 7.83 | 1.76 | 1.22 | 0.92 | 9.06 | 57.32 | 814.47 |
| ***S.gal*** | 5.42 | 17.40 | 6.52 | 6.53 | 8.34 | 1.49 | 1.34 | 0.98 | 9.76 | 57.77 | 811.44 |
| ***S.hab*** | 6.10 | 19.86 | 5.69 | 6.93 | 9.74 | 2.15 | 1.12 | 1.17 | 6.90 | 59.64 | 893.57 |
| ***S.pen*** | 5.48 | 22.13 | 6.79 | 4.11 | 14.88 | 1.12 | 1.07 | 1.37 | 8.04 | 64.99 | 1072.33 |
| ***S.tub*** | 5.00 | 24.43 | 11.15 | 1.93 | 4.46 | 0.39 | 2.07 | 1.92 | 5.53 | 56.88 | 737.87 |

For most of the *Ty1/Copia* and *Ty3* lineages, the abundance of intact elements is stable across the domesticated tomato accessions, while it is higher in wild tomato species and highest in the most distant relative *S. pennellii*^3^ (Fig. 3a). The *Ty1/Copia Tork* lineage is exemplary with 500 elements in MbTMV (domesticated tomatoes have between 480-505 *Tork* copies), while there are more than 1,100 in *S. pennelli* (Fig. 3a). The abundant *Ty3 Tekay* lineage is represented by 1261-1380 elements in the four cultivated tomato accessions Micro-Tom, Heinz, M82 and Sweet100, while in *S. pennellii*, there are around 3,200 *Tekay* elements, which makes it the most successful lineage in the ten species (Fig. 3a). We found that MbTMV has approximately 500 more *Tekay* elements compared to the other domesticated species (Fig. 3b). *Ty1*/*Copia* and *Ty3* elements are less abundant in potato than the tomato accessions, with the only exception being the prolific *Ale* lineage (Fig. 3a,b). Overall, the variation in specific *Ty1/Copia* and *Ty3* lineage abundances indicated that the aging of individual elements would allow us to understand how TE dynamics may have shaped the evolution of the *Lycopersicon* clade.

The age of LTR retrotransposons provides insights into transpositional activity in plant genomes^69^. Younger insertions suggest recent bursts of transposition whereas older elements are more degraded^70,71^. Estimating the timing of LTR insertions helps identify genome expansion and could indicate stress-induced mobilisation over evolutionary periods^72,73^. We determined the age of the intact LTR retrotransposons by using the LTR identity metric, which shows how much the LTRs have altered since the insertion time. We found that LTR retrotransposons are younger and more recently integrated in wild tomato genomes compared to domesticated accessions (Fig. 3c). Moreover, MbTMV has more young elements compared to the other four domesticated accessions, while *S.cheesmaniae* and *S.galapagense* have an intermediate age profile compared to *S. habrochaites* and *S. pennellii* that have the most recent integrations (Fig. 3c).

The high abundance of *Tekay* elements in MbTMV, together with the accumulation of many young LTR retrotransposons in this species, led us to investigate whether there is a connection between those characteristics. To this end, we examined the LTR identity of all the LTR retrotransposon lineages in MbTMV separately. We found the *Ogre* lineage is the youngest, while the *Tekay* lineage is the second youngest and contains the most elements (Fig. 3b and Supplementary Fig. 9b and 10a). Generally, *Tekay* is the most successful LTR retrotransposon lineage in the *Lycopersicon* clade^30,31^; hence, we intended to categorise *Tekay* elements based on their integration time into the genome (see Materials and Methods). Utilizing this age classification, we were able to quantify the proportion of different-aged elements in the ten species (Fig. 3d). Correspondingly, the majority of *Tekay* elements in *S. habrochaites* and *S. pennelli* are young, while for the rest of the species, they are of moderate age (Fig. 3d). Interestingly, the proportion of young elements in MbTMV is higher compared to the rest of the domesticated accessions (Fig. 3d). Finally, the fraction of old elements is always the lowest, illustrating the difficulty in identifying TEs after mutation accumulation.

Due to the presence of a 64.1 Mbp introgression from *S. peruvianum* on chromosome 9 of MbTMV that contains the *Tm2*^2^ TMV resistance gene^53^, we suspected this could be the source of young *Tekay* elements within MbTMV. We located the young, moderate and old *Tekay* elements on chromosome 9 from the ten accessions, and we found a specific enrichment of more than two-thirds of young *Tekay* elements on MbTMV chromosome 9 that was not identified in the other accessions (Supplementary Fig. 10b) (Fig. 3e). We validated that the majority of the young *Tekay* elements in MbTMV are found in the pericentromeric region of chromosome 9 which is the location of the *S. peruvianum* introgression^53^ (Fig. 3f). The abundance and location of young *Tekay* elements is different between the chromosome 9 copies of MbTMV, Heinz1706 and *S. pennellii* (Fig. 3e and 3f). Finally, a complete phylogenetic reconstruction of all *Tekay* elements in the MbTMV genome was carried out using the integrase gene of each element (Fig. 3g). We found that the majority of young elements on chromosome 9 are clustered together with a maximum likelihood value of 0.74, indicating those sequences were not randomly clustered together (Fig. 3g). Summarising, the introgression of the TMV resistance gene in MbTMV and all modern commercial tomato varieties has brought with it an abundance of *Tekay* TEs that likely exploded in the wild *S.peruvianum* genome in an independent event from *S. pennellii*.

### Higher crossover rate in female meiosis underlies female-enhanced recombination regions in tomato intraspecific and interspecific hybrid backcross populations

To investigate the effect of sequence divergence and reproductive gender on crossover number and location, we pollinated *S. lycopersicum* cv. MbTMV with three other parents (*S. lycopersicum* cv. Micro-Tom, *S. cheesmaniae* LA1039 and *S. pennellii* LA0716), generating three sequence-divergent F1 hybrids with increased amounts of genomic variation (Fig. 4a, Supplementary Table 7). Next, we generated three male and three female backcross populations derived from the intra- and interspecific F1 hybrids (See Materials and Methods). From the six backcross populations a total of 709 individuals were subjected to low-coverage sequencing resulting in 2.5-5 Gbp data per individual sample (Supplementary Fig. 11-13).

A total of 6,853 crossovers were detected from all the samples, divided into female and male populations that represent recombination in female meiotic cells and male meiotic cells, respectively. We anaylzed recombination landscapes for the six backcross populations (Fig. 4b) and found that recombination is skewed to the distal euchromatic arms in all cases, is consistent with previous observations in tomato^74–76^. We found that all three female backcross populations have significantly more crossovers than their counterpart male backcross populations (Fig. 4c), which is consistent with previous findings on heterochiasmy in a *S. lycopersicum* x *S. pennellii* hybrid^51^. The respective female populations have 36% (MbTMV x Micro-Tom; hereafter MbTMV-MT), 35% (*S. lycopersicum* x *S. cheesmaniae*; hereafter MbTMV-S.che) and 28% (*S. lycopersicum* x *S. pennellii;* hereafter MbTMV-S.pen) higher CO number than the counterpart male populations (Fig. 4b). The consistently higher CO rate in the three female populations suggests that heterochiasmy is conserved in intraspecific hybrid tomato and minimally divergent interspecific hybrid tomato. Although we observed higher CO numbers in interspecific than intraspecific populations, some COs in MbTMV-MT backcross offspring may not be detected due to the absence of SNPs in some genomic regions that are identical by descent. Compared to the previous study^51^, we now have high-resolution mapping of crossover sites that enable examination of recombination sites in finer detail, revealing female-enhanced recombination regions (FRRs) such as those in the short arms of chromosomes 1, 6, 7, 8 and 10 (Fig. 4b and Supplementary Fig. 14). Comparing the overall recombination landscapes, heterochiasmy in tomato appears to be confined to specific genomic regions and is not a genome-wide effect. We selected chromosome 7 and examined the allele ratios in the offspring populations and found no evidence of segregation distortion. Overall, our analysis identified megabase-scale FRRs in three different tomato hybrids that appear to be over-represented on short chromosome arms.

Apart from the effect of reproductive gender on crossover number, we found that the genetic differences between the parental genomes also influence the distribution of COs along the chromosomes. Between 67% (MbTMV-MT) and 80% (MbTMV-S.pen) of the tomato genome is suppressed of recombination, limiting the genetic shuffling and resulting in linkage drag (Fig. 5a). Among these CO-suppressed regions, which we call coldspots, about 64% (529 Mb) are conserved across all three hybrid backgrounds. Conserved coldspots, spanning tens of megabases, are mostly in the pericentromeric regions while smaller cross-specific coldspots are located at the distal chromosomal ends (Fig. 5b). There is a significant depletion of COs in regions with structural variants (p-value < 0.0001, Fisher’s exact test), concordant with previous studies in *A. thaliana* and tomato^42,43,76^. The elevated proportion of coldspots in both MbTMV-S.pen backcross populations is partly due to the presence of large inversions between the *S. lycopersicum* and *S. pennellii* genomes (Supplementary Fig 15). The short arms of chromosome 6 and 7 contain inversions between the *S. lycopersicum* and *S. pennellii* genomes that suppressed recombination, consistent with previous observations^68,76^. As reported above, we discovered a 2.2 Mbp inversion in chromosome 8 between the eight *Lycopersicon* genomes and *S. pennellii* and found that in the MbTMV-S.pen population where this inversion is heterozygous no COs occur. In contrast, the same region in the other two hybrids, where no inversion is present, meiotic COs can occur (Fig. 2a, Fig. 5c and Supplementary Fig. 16). This inversion overlaps 268 *S. lycopersicum* and 278 *S. pennellii* genes, implying that both agronomically desirable and detrimental alleles are inherited together. The assembly of *S. pennellii* genome not only enables the discovery of many other unknown SVs relative to *S. lycopersicum* but also reveals massive linkage drag that may result from introgressing *S. pennellii* segment into the domesticated tomato.

### Novel retrotransposon expansions and genome inversions associated with recombination coldspots in a divergent tomato hybrid

Similar to SVs, retrotransposons such as *Gypsy* and *Copia* have been associated with CO suppression as well^77–82^. To further determine the influence of SVs and TEs in patterns of COs across all three populations, we identified the genomic positions of the coldspots relative to the assemblies of each parent. After mapping coordinates, we found that some coldspots have different lengths based on one of the parental genomes, implying that the genome may contain deletions or insertions or may have undergone repeat expansion or contraction (Fig. 5d and Supplementary Fig. 17). MbTMV-MT coldspots are located in regions where MbTMV and MicroTom are syntenic but have high density of *Gypsy* and *Copia*. On the contrary, several MbTMV-S.che and MbTMV-S.pen coldspots overlap regions with chromatin expansions or inversions between the parents. Compared with Mb-MT and MbTMV-S.che, the retrotransposon-associated coldspots in MbTMV-S.pen contain significantly more retrotransposons (Supplementary Fig. 18), consistent with our results above about the expansion of *Tekay* elements in *S. pennellii* compared to other tomato and wild species. Apart from repeat elements, some coldspots contain novel insertions or translocations that may affect synapsis during pachetene, which in turn prohibits recombination just like in *A.thaliana*^42,43,83^. Moreover, some of these insertions overlapping Mb-Pen coldspots contain hemizygous genes (Fig. 5e), which is important for breeding because these genes may confer novel functions but at the same time are tightly linked. Utilizing our high-quality genomes, we identified large structural genomic variants and differential TE composition between parental genomes that directly suppressed recombination. Our results add further resolution and insights to previous findings on heterochiasmy, tightly linked genes^51^ and suppressed recombination between *S. lycopersicum* and other wild species^68,84^.

## Discussion

Here, we have assembled high-quality *de novo* chromosome scale genomes of *S. cheesmaniae* LA1039 and *S. pennellii* LA0716. We compared our two new assemblies with high-quality assemblies of eight related accessions and species, and extensively analysed the TE content of all genomes. Finally, three different hybrids (*S.pennellii*, *S. cheesemaniae* and *Solanum lycopersicum* cv. Micro-Tom crossed with *Solanum lycopersicum* cv. Moneyberg-TMV) were used to develop large recombinant offspring populations to reveal the genomic basis of heterochiasmy in hybrid tomato.

### An improved *S. pennellii* assembly and a novel *S. cheesmaniae* assembly

Our improved assembly of *S. pennellii* LA0716 integrated an additional 145.9 Mbp to the 12 chromosomes including in regions that are rich in repetitive elements (Fig. 1). Sequencing data coverage analysis showed that the improved assembly had fewer deviating regions, unmapped and supplementary reads compared to the original assembly^3^. The improved *S. pennellii* assembly contains inversions at the start of chromosomes 6 and 7 that were not present in the original assembly but are confirmed by BAC-FISH results^68^. The improved assembly also includes a 7.1 Mbp inversion on chromosome 3 and a 2.2 Mbp inversion at the start of chromosome 8 that are validated by Hi-C and our recombination mapping results (Fig 1a, Fig 4b, Supp). The *S. cheesmaniae* LA1039 assembly gives new genomic insights into an understudied species that contains useful salinity stress tolerance that could be utilized for the improvement of the cultivated tomato^85^. As a relatively close relative of the cultivated tomato the *S. cheesmaniae* assembly shows that in the euchromatic arms are largely syntenic to *S. lycopersicum* but we did discover a 35.7 Mbp inversion in the pericentromeric region of chromosome 5 (Fig 2a). Overall, the improved *S. pennellii* LA0716 and novel *S. cheesmaniae* LA1039 genomes improve structural information and gene identification in those accessions, and can facilitate breeding in interspecific crosses.

### Long reads as a lens for the structural evaluation of the *Lycopersicon* clade

Over the last ten years, developments in long-read sequencing technologies have revolutionized the field of genomics^19,86–89^. PacBio HiFi produces highly accurate reads, limited in size (generally 20kb), whereas ONT reads are somewhat less accurate but generally longer (up to 200kb). Structural variants can be detected directly from long-read sequencing^90^, however, genome assembly is needed to properly detect and characterize SVs in the megabase scale^19,53^. Recently, the construction of a tomato super-pan genome assembled from legacy PacBio data has led to insights in SVs within the whole tomato clade and discovery of a wild tomato gene with the potential to increase yields^13^. However, our results show PacBio HiFi and ONT ultra-long sequencing can lead to improved genome assemblies, and it is likely ONT ultra-long sequencing will prevail in challenging repetitive genomes in future. High quality *de novo* genomes have been assembled from HiFi^8,91^ or ONT^10,19^, however a combination of both approaches can overcome extended repetitive regions and generate *de novo* assemblies of the highest quality^88,89,92^.

We have used a combination of long-read sequencing data with chromosome confirmation capture data yielding high-quality chromosome scale reference genomes which enabled the accurate detection of large structural variants spanning several megabases. Thus we identified several novel megabase-scale inversions, including a roughly 20 Mbp inversion on chromosome 5 that is specific to the Galapagos tomato species *S. galapagense* and *S. cheesmaniae* (Fig. 2). Our finding that the complete chromosomes 6, 7 and 8, inclusive of the pericentromeric regions, are structurally almost identical in all five cultivated tomatoes is consistent with the thesis that these chromosomes contain a key set of agriculturally relevant genes. Short-read sequencing has shown chromosomes 6, 7 and 8 have low nucleotide diversity in fresh consumption and modern processing tomatoes^93^, while low numbers of SVs were detected in cultivated tomatoes in the pericentromeric regions of the same chromosomes^19^. Notably chromosome 7 contains several sweep loci including *fw7.2*, while chromosome 6 additionally contains a fruit weight locus embedded within the pericentromere^94^. Our results have shown pairwise comparison of *de novo* reference quality chromosome scale genomes leads to full characterization of genomic variants between species along the chromosomes. Our high-quality genomes further the recently constructed tomato super-pan genome and facilitate the characterization of genomic variation between cultivated tomato and wild related species^12,13^.

### Diversity of TE and repeat profiles in the *Lycopersicon* clade

Transposons and repeats play a crucial role in plant evolution, diversity, and adaptation. We demonstrated that the repeat component is increased in wild relatives of the domesticated tomato (Fig. 1e-f and Supplementary Fig. 6), and this increase also includes TEs, regardless of class (Fig. 3a-b and Supplementary Fig. 9). Additionally, most of the LTR retrotransposons in these species have recently been integrated into the genome (Fig. 3c and Supplementary Fig. 10a), and the *Tekay* lineage is the most abundant and one of the youngest across the *Lycopersicon* clade (Supplementary Fig. 10a). The *S. lycopersicum* cv. MbTMV shows an increase in *Tekay* elements compared to the other domesticated accessions specifically on chromosome 9 (Fig. 3d and Supplementary Fig. 10b). MbTMV is known for resistance to tobacco mosaic virus (TMV), attributed to the *S. peruvianum* introgression on chromosome 9^53^. Consequently, along with the resistance trait, a significant number of *Tekay* elements were introduced (Fig. 3f), and phylogenetic analysis indicated that most of the young elements on chromosome 9 share a common origin (Fig. 3g).

In summary, the overall amount of TEs and repeat content is reduced in cultivated accessions. This reduction may be linked to the domestication of tomato, which significantly decreased the tomato mobilome, possibly due to a genetic bottleneck caused by increased inbreeding^95^. Similarly, wild rice species exhibit extensive variation in TE content that correlates with genome size differences and LTR retrotransposon expansions, compared to the domesticated rice, which has more constrained TE landscapes reflecting both TE amplification in wild lineages and TE reduction during the domestication and selection process^96^. Targeted comparisons between cultivated and wild rice suggest that artificial selection during domestication may have purged TE insertions that negatively influence gene function or expression stability^97^. In essence high mobilome activity may compromise the long-term viability of cultivated accessions. Conversely, introgression of genetic material from wild germplasm may serve to invertedly introduce young TEs into the breeding germplasm, as in the MbTMV case, and this possibility warrants further investigation.

### Recombination analysis in sequence-divergent tomato hybrids

Understanding where meiotic crossovers (COs) occur is crucial for understanding the evolution of sexually reproducing species and for optimizing crop breeding. Hybridization of three sequentially divergent species (*S. lycopersicum* cv. Micro-Tom, *S. cheesmaniae* LA1039 and *S. pennellii* LA0716) to *S. lycopersicum* cv. Moneyberg-TMV and generation of six derived male and female backcrosses populations (Fig. 4a) allowed for investigation of genomic polymorphisms on recombination distribution and heterochiasmy. We have found that large recombination coldspots are mostly located in heterozygous inversions or retrotransposon-rich pericentromeric regions. We detect meiotic recombination events that are inherited in viable progeny and it is plausible that crossovers within inversions could produce unviable gametes due to non-reciprocal exchanges. The divergent MbTMV-S.pen hybrid populations have less sites for recombination, consequently limiting the shuffling of alleles in the short arms of chromosome 6, 7 and 8 that are crucial for breeding (Fig. 4b). Knowing the patterns of recombination in a specific hybrid cross enables careful inspection of potential regions to introgress from wild species into breeding lines, revealing possible consequences of tight linkage such as those ones due to large inversions.

The heterochiasmy consistently observed in all three tomato F1 hybrids suggests that elevated meiotic CO rate in female meiosis is likely universal throughout the tomato clade. Opposite to tomato, *A.thaliana* has higher male recombination rate than female, in both inbreds and hybrids^47–50^, whereas heterochiasmy is not observed in the maize B73 x Mo17 hybrid^98,99^. We noticed that female-enhanced recombination regions (FRRs) were more abundant on the short chromosome arms in the tomato genome. In male meiosis, the longer, more gene-rich, arms accumulate crossovers more easily, whereas in female meiosis, where total crossover number is higher, the shorter arms also begin to accumulate crossovers. This observation is potentially consistent with the recently proposed coarsening model for meiotic crossover interference where well-spaced crossover-promoting HEI10 foci determine the crossover landscape^100–102^. In the future, heterochiasmy in tomato hybrids could be studied by the mutagenesis of genes involved in crossover in control in the hybrids studied herein. Moreover, it is clear from our results that tomato introgression breeding should clearly focus on using hybrid (or partially hybrid) genotypes as mother, and the recurrent inbred parent as father, in order to maximize crossover number and exploit female-enhanced recombination regions to expedite breeding.

## Materials and Methods

### Plant materials and growth

Seeds of S. lycopersicum cv. Moneyberg-TMV, *S. lycopersicum* cv. Micro-Tom and *S. cheesmaniae* LA1039 were pre-soaked in water for 2 hours, followed by 15 min incubation in Na_3_PO_4_, 30 min incubation in 1:1 (volume:volume) with water-diluted household bleach (DanKlorix 2.8g NaOCl per 100g liquid) and at least 3 subsequent washes with sterilised water. Seeds of *S. pennellii* LA0716 were soaked in 700 ppm GA3 for 20 hours, followed by 30 min incubation in 1:1 (volume:volume) with water-diluted household bleach (DanKlorix 2.8g NaOCl per 100g liquid) and at least 3 subsequent washes with sterilised water. All seeds were sown on 0.8% agar and incubated in growth chambers with 16h daylight, and 25°C daytime and 21°C nighttime temperatures. For seed origin information, please see the acknowledgements.

Seedlings of *S. lycopersicum* cv. Moneyberg-TMV, cv. Micro-Tom and *S. pennellii* LA0716 were transplanted to MPIPZ greenhouses and grown during late spring of 2020 under natural light supplemented with artificial light to ensure 16h light per day. Seedlings of *S. cheesmaniae* LA1039 were transplanted to Bronson growth chambers and grown in short-day conditions (12 hours of daylight) with 28°C daytime, 22°C night-time and 60-80% relative humidity, to mimic native growth conditions.

### Hybridization

Immature unopened green to yellowish Moneyberg-TMV flowers were carefully emasculated using forceps and pollinated with freshly extracted pollen of the other parent (e.g. Micro-Tom, *S. cheesmaniae* LA1039 and *S. pennellii* LA0716) 1-3 days after emasculation. Pollinated Moneyberg-TMV inflorescences were labelled with crossing dates and left to develop fruits. Mature fruits were collected, after which seeds were extracted and cleaned in a 1:1 (volume:volume) pulp to 2% HCl solution for 1 hour, followed by rinsing with excess water and drying at room temperature.

To overcome possible immature fruits, Moneyberg-TMV x *S. pennellii* LA0716 were collected 28-33 days after pollination, sterilised in 6% NaOCl for 10 minutes, followed by 3 consecutive washes in sterile water. Embryos were carefully removed from the endosperm using a binocular and transferred to 0.5 MS including vitamins (Sigma) agar plates containing 10g sucrose/L, followed by incubation in growth chambers with 16h daylight, and 25°C daytime and 21°C nighttime temperatures.

To generate backcross populations, the three F1 hybrid genotypes were cultivated in the greenhouse and used for reciprocal crosses with *S. lycopersicum* cv. Moneyberg-TMV, apart from the *S. lycopersicum* Moneyberg-TMV x *S. pennellii* LA0716 female population where *S. pennellii* LA0716 was used as pollen donor (*S. lycopersicum* Moneyberg-TMV x *S. pennellii* LA0716 does not accept pollen from *S. lycopersicum* Moneyberg-TMV due to Unilateral interspecific incompatibility)^103^.

### High Molecular Weight (HMW) DNA extraction and sequencing

HMW DNA of LA1039 and LA0716 was isolated from 1.5 g of young leaf material with a NucleoBond HMW DNA kit (Macherey Nagel, Düren, Germany). DNA quality was assessed with a FEMTOpulse device (Agilent, Santa Clara, CA, USA), and quantity was measured using a Quantus Fluorometer (Promega, Madison, WI, USA). HiFi libraries were prepared according to the manual “Procedure & Checklist – Preparing HiFi SMRTbell® Libraries using SMRTbell Express Template Prep Kit 2.0” with an initial DNA fragmentation by Megaruptor 3 (Diagenode) and final library size binning into defined fractions by SageELF (Sage Science). Size distribution was again controlled by FEMTOpulse (Agilent). Size-selected libraries with expected insert sizes of 20 and 27 Kbp for LA1039 and 19 and 25 Kbp for LA0716 were independently sequenced on a single SMRT cell of a Pacific Biosciences Sequel II on Sequel device at the MPGC Cologne with Binding Kit 2.0 and Sequel II Sequencing Kit 2.0 for 30 h (Pacific Biosciences).

K-mer analysis of HiFi data was performed using the k-mer Analysis Toolkit (KAT) hist v2.4.1 with options -m 21 (Mapleson *et al.*, 2017) and plotted using R v.4.2.3 (https://cran.r-project.org/) and packages from the Tidyverse collection v2.0.0^104^ and utilising the maximum number of distinct k-mers within the homozygous peak to scale the y-axis.

High-molecular weight DNA that was used for PacBio sequencing was size selected using the Circulomics Short-Read Eliminator XL Kit (Circulomics Cat# SKU SS-100-111-01). The size-selected DNA was used as an input for 10 library preparations with the Oxford Nanopore SQK_LSK112 ligation sequencing kit. Sequencing was performed on an Oxford Nanopore PromethION on 10 Flowcells FLO-PRO112 (R10.4) and sequenced for up to 72h with reloading and washing where needed. ONT-basecalling was performed with “super accuracy” model (dna_r10.4_e8.1_sup) using guppy_6.0.1/2 or guppy_6.1.1 (https://nanoporetech.com/software/other/guppy/).

Chromatin conformation capture libraries were prepared using 0.5 grams of young leaf material as the input. All treatments were according to the recommendations of the kit vendor (Omni-C, Dovetail) for plants. As a final step, the Illumina-compatible libraries were prepared (Dovetail) and test sequenced (paired-end 2 x 150 bp) on an Illumina NextSeq 2000 device at the MPGC in Cologne, followed by sequencing on a NovaSeq 6000^105^ at Novogene for increased coverage.

### DNA extraction and sequencing of BC populations

Young true leaf material was collected and lyophilised overnight in an Alpha 1-4 freeze dryer (Martin Christ GmbH). DNA extraction was performed on the KingFisher robot (Thermofisher) using the NucleoMag Plant Kit (Macherey and Nagel, P/O 744400.4) and eluted in 200 μl buffer. Transposase-based libraries (Tn5, Nextera) were prepared according to^42^ and test sequenced on an Illumina NextSeq 2000 in the MPGC Cologne. BC samples were outsourced to Novogene and sequenced on the NovaSeq 6000, aiming to generate approximately 3-4 Gb of paired-end data per sample. Sequencing data were provided as deduplicated and adapter-removed reads in zipped FASTQ format.

### Sequencing data statistics were obtained using seqkit stats v2.0.0^106^ with options -a and FastQC with default settings (Andrews S. (2010). FastQC: a quality control tool for high-throughput sequence data. Available online at: https://www.bioinformatics.babraham.ac.uk/projects/fastqc/)

### Genome assembly

#### Assembly

Hifiasm v0.16.1-r375^59^ was used with option -l0 to assemble the HiFi reads of LA1039. LA0716 ONT reads were filtered for length (> 90 Kbp) and quality (q > 90) using Filtlong-0.2.1 (--min_length 90000 --min_mean_q 90). Hifiasm v0.17.6-r460^59^ was used to assemble HiFi and filtered ONT reads with options -ul.

#### Scaffolding

The LA1039 and LA0716 assemblies were scaffolded following the esrice/hic-pipeline (https://github.com/esrice/hic-pipeline). First, Omni-C (Dovetail) paired-end reads were mapped separately using Burrows-Wheeler Aligner v0.7^107^ and the filter_chimeras.py script. Next, the mapped reads were combined using the combine_ends.py script, with three iteration steps and a minimum mapping quality value of 20 (default), followed by adding read mate scores, sorting and removing duplicate reads using SAMtools v1.9^108^. The resulting BAM file was converted to a BED and sorted (-k 4) using BEDtools v2.30^109^. Per assembly, we ran one single round of Salsa v2.2^110^ with optional settings: -e DNASE -m yes -p yes. A modified version of the convert (convert.sh) script was used to convert the Salsa2 output to a Hi-C file, which was utilised within a local installation of Juicebox v.1.11.08 (https://github.com/aidenlab/Juicebox) to generate a Hi-C contact plot.

#### D-genies alignment to MbTMV

The LA1039 and LA0716 genome assemblies and scaffolded genome assemblies were aligned to MbTMV^53^ using Minimap2 2.24-r1122^111^ with option -x asm5 to generate pairwise alignment format files. Used reference and query genomes were indexed using a Python-based custom D-genies index script, followed by generating dotplots using D-GENIES version 1.4.0^112^.

#### Manual fine-tuning

Assemblies scaffolded with the automated Salsa2 pipeline were further fine-tuned, whereby smaller scaffolds were manually placed within larger scaffolds, based on Hi-C interaction and alignment to MbTMV. Fused chromosome-scale scaffolds, observed for LA0716 (chromosomes 1 and 2), likely a result of interaction between chromosome-wide repetitive elements (e.g. LTR-RTs) or specific repeat regions (e.g. TGR1) present in tomato^53,113^, were manually broken at the ends of each chromosome based on Hi-C interaction and alignment to MbTMV. Multiple rounds of fine-tuning were performed, whereby changes in each round were kept as AGP files, followed by conversion into FASTA files using RagTag agp2fasta^52^ with gap sizes of 100 N’s connecting scaffolds. After each round, the generated FASTA files were checked by alignment to MbTMV and Hi-C interaction as described previously. Broken contigs that were scaffolded after another were checked for continuity by alignment of HiFi reads using pbmm2 v1.7.0 (https://github.com/PacificBiosciences/pbmm2) and ONT reads in the case of LA0716 using Minimap2 version 2.24-r1122^111^ as described below and repaired if supported by read continuity.

### Validation of genome assemblies

#### Statistics

For the validation of the 12 assembled chromosomes, we checked assembly statistics, BUSCO completeness, LTR assembly index, and both base quality and k-mer completeness. Quast v5.0.2^114^ and GAAS v1.1.0 (https://github.com/NBISweden/GAAS) were used to calculate statistics on fasta files. BUSCO was calculated using BUSCO v5.2.1^115^ depending on hmmsearch v3.1, and metaeuk v4.a0f584d was used with lineage datasets solanales (https://busco-data.ezlab.org/v5/data/lineages/solanales_odb10.2020-08-05.tar.gz) to obtain evolutionarily informed expectations of gene content. To assess the LAI of each genome assembly, LTR retriever v2.9.0^61^ (Ou *et al.*, 2018) was run with recommended default settings invoking GenomeTools v1.6.2 (https://github.com/genometools/genometools) with options - minlenltr 100 -maxlenltr 7000 -mintsd 4 -maxtsd 6 -motif TGCA -motifmis 1 -similar 85 -vic 10 -seed 20 -seqids yes and LTR_FINDER_parallel v1.2 (https://github.com/oushujun/LTR_FINDER_parallel) with options -threads 10 -harvest_out - size 1000000 -time 300. Base accuracy (QV) and k-mer completeness were assessed using Merqury v1.3^63^ with default settings, using the read databases and genomes. The read database for the Merqury run was generated using the meryl algorithm v1.3, with k-mer size equal to 20 and default parameters, which is also part of the Merqury genome evaluation tool^63^.

#### Read alignments & coverage analysis

PacBio HiFi data were mapped to each corresponding genome using pbmm2 (version 1.7.0) with the preset ‘CCS’ (https://github.com/PacificBiosciences/pbmm2). ONT data were mapped to each corresponding genome using Minimap2 version 2.24-r1122^111^ with the preset ‘map-ont’ and ‘eqx’, followed by converting, sorting and indexing of the output file using Samtools v1.9^108^. Primary read coverage (default) was extracted in 100-kbp windows using Mosdepth v0.3.2^116^ and plotted in the middle of each interval using R v.4.2.3 (https://cran.r-project.org/) and packages from the Tidyverse collection v2.0.0^104^.

#### Re-mapping Omni-C

Following the esrice/hic-pipeline (https://github.com/esrice/hic-pipeline), we separately mapped Omni-C (Dovetail) paired-end reads to each genome using Burrows-Wheeler Aligner v0.7^107^ and the filter_chimeras.py script. Mapped reads were combined using the combine_ends.py script, with three iteration steps and a minimum mapping quality value of 20 (default), followed by adding read mate scores, sorting and removing duplicate reads using SAMtools v1.9^108^. The resulting BAM file was converted to a BED file and sorted (-k 4) using Bedtools v2.30^109^. We ran one single round of Salsa v2.2^110^ with optional settings: -e DNASE -m yes -p yes. A modified version of the convert.sh script (https://github.com/marbl/SALSA) was used to convert the Salsa2 output to a Hi-C file, which was used within a local installation of Juicebox v.1.11.08 (https://github.com/aidenlab/Juicebox) to generate a Hi-C contact plot.

#### Alignment to MbTMV

The finalised LA1039 and LA0716 genomes were aligned to MbTMV^53^ using Minimap2 2.24-r1122^111^ with option -x asm5 to generate pairwise alignments. Used reference and query genomes were indexed using a Python-based custom D-genies index script, followed by generating dotplots using D-GENIES version 1.4.0^112^.

### Annotation of sequence features

#### Nucleotide composition

To obtain nucleotide composition in 100kbp windows of each genome, we first converted the FASTA index file to a genome.txt file by extracting only the first two columns, including the sequence identifier and length. Next, we used BEDtools makewindows command v.2.29.0^109^ to generate 100kbp windows in BED format. Finally, we extracted the nucleotide composition using seqtk comp command v.1.3 (https://github.com/lh3/seqtk) with each respective FASTA file and 100kbp window BED file as inputs. Nucleotide composition was converted to a ratio relative to each other nucleotide and plotted on the midpoint of each interval using R v.4.0.3 (https://cran.r-project.org/) and packages from the Tidyverse collection v2.0.0^104^.

#### Simple sequence repeats

Simple sequence repeats (SSR’s) were identified using MISA (https://github.com/schellt/msat-loci-identification) running perl (v.5.24.1) invoking the following default settings: definition (unit_size, min_repeats): 1-10 2-6 3-5 4-5 5-5 6-5, interruptions (max_difference_between_2_SSRs): 100, GFF: true. Generated GFF files were modified using a series of Linux and R commands, where identified repeats were summarised in 100Kbp windows per class (e.g. monomer, dinucleotide repeat, trinucleotide repeat, etc.) and plotted on the midpoint of each interval using R v.4.0.3 (https://cran.r-project.org/) and packages from the Tidyverse collection v2.0.0^104^.

#### Long terminal repeat retrotransposons (LTR-RT)

The reported LTRs by the LTR-retriever and LAI pipeline “pass.list.gff3” were extracted using a custom pipeline invoking grep and awk commands. In short, we selected for reported LTRs on chromosomes 1 to 12 and extracted reported chromosome information, repeat region, starting position end position which were used for length calculation before plotting the distribution over the genomic position using R v.4.0.3 (https://cran.r-project.org/) and packages from the Tidyverse collection v2.0.0^104^.

#### Gene annotation

An integrated pipeline composing both ab-initio and evidence-based approaches was employed for MbTMV and LA0716 structural annotation.

The ab-initio predictions were performed for MbTMV, LA1039 and LA0716 genomes using Helixer v0.3.3 with the model land_plant_v0.3_a_0080.h5, integrating deep learning and Hidden Markov Models to enable accurate ab-initio gene annotations^117^, with subsequence-length set to 504000. For the post-processing into primary gene models with HelixerPost, the parameters window_size 100, edge_thresh 0.1, peak_thresh 0.80, and min_coding_length 60 were applied. For LA0716, HelixerPost was utilised using peak_thresh 0.60.

For evidence-based gene predictions for MbTMV and LA0716 genomes, a multi-step approach which involved both short- and long-read datasets was employed. In brief, publicly available short-read RNA-seq data (Supplementary Table 8) were mapped to the genomes using Hisat2 v2.2.1^118^ with the parameter –dta. PacBio Isoseq reads (Supplementary Table 9) were mapped to the respective genome using minimap2 v2.28^111^ with parameters -ax splice:hq and -uf. Subsequently, all of the mapped reads were merged using SAMtools v1.20^108^ and sorted with -m 5G and 3G for MbTMV and LA0716, respectively. Finally, transcript assemblies were generated using StringTie v2.2.3^119^ with --mix -L -j 3 -f 0.1 as well as -c 10 and -c 50 for MbTMV and LA0716, respectively.

Subsequently, for the filtering of valid splice junctions from pre-aligned RNA-seq data for MbTMV and LA0716 genomes, Portcullis 1.2.4^120^ was used.

Finally, evidence from both the ab-initio and homology-based gene models, together with validated splice junctions, was integrated using Mikado v2.3.3^121^. For MbTMV, weighting factors favouring ab-initio evidence (three for Helixer and one for StringTie) resulted in a reliable consensus gene model with high accuracy and completeness. For LA0716, weighting was balanced for ab-initio and homology-based evidence (one for Helixer and one for StringTie). Furthermore, the LA0716 gene model was subjected to the downstream fine-tuning steps through validation techniques to filter out potential false positives (Supplementary Fig. 19) as follows.

A Pfam-based validation where the high-confidence sequences were identified by searching against the Pfam database using MMseqs2 v16-747c6^122^. This step yielded a “Pfam-verified” protein set comprising 10,154 entries with protein domains. Secondly, validation was conducted against a curated custom database built from UniProtKB entries taxonomically restricted to *Viridiplantae* (taxonomy ID 33090, reviewed as of February 18, 2025). This database was merged with proteins validated by PSAURON v1.0.0^123^ from *Arabidopsis thaliana* (Araport11, https://www.arabidopsis.org/) and *Oryza sativa* (osa1r7, https://rice.uga.edu/). The resulting “CustomDB I-verified” set consisted of 33,023 high-confidence proteins. Thirdly, proteins were checked with PSAURON v1.0.0, generating a “PSAURON-verified” set containing 49,488 reliable proteins. Finally, all the verified proteins from the previous steps were re-evaluated against a second custom reference database. This “CustomDB II” included PSAURON v1.0.0-validated proteins from *Solanum lycopersicum* (ITAG4.0: https://solgenomics.net/) and *Solanum tuberosum* (PGSC6.1, https://solgenomics.net/), using MMseqs2 with a stringent coverage threshold -c 0.9. This final validation yielded a refined set of 37,099 verified proteins (comprising 31,998 representative sequences), culminating in a highly reliable consensus gene model characterised by both high accuracy and completeness.

The agat_sp_keep_longest_isoform.pl script from AGAT v1.4.0 (https://github.com/NBISweden/AGAT/releases/tag/v1.4.0) and Gffread v0.12.7^124^ were used, respectively, to retain the longest isoform per gene locus and to extract the corresponding protein or coding sequences (CDS) from the gene models.

The quality of the gene repertoires annotated from genomic sequences was further validated based on the BUSCO completeness calculated using compleasm v0.2.6^125^ with the lineage datasets solanales_odb10 and estimations for the proportion of accurate and erroneous gene models in the proteomes were performed using OMArk v 0.3.0^126^ with the OMA orthology database, in which exploits orthology relationships.

### Characterization of genomic variation

To identify homozygous SNPs between Moneyberg-TMV and Micro-Tom, LA1039 and LA0716 respectively, we aligned HiFi reads of Micro-Tom, LA1039 and LA0716 to MbTMV using pbmm2 (version 1.7.0) with the preset ‘CCS’ (https://github.com/PacificBiosciences/pbbioconda). Next, we used GATK HaplotypeCaller v4.2.5.0, HTSJDK v2.24.1 and Picard v2.25.4^127^ with default options to identify variants. To remove possible variants due to mapping artefacts and obtain only homozygous SNPs, we filtered the identified variants for INFO/DP => 5% percentile, INFO/DP <= 95% percentile, INFO/MQ => 5% percentile, GT = “1/1” and TYPE = “snp” using BCFtools v1.15^108^.

#### Synteny and Rearangement Identifyer and Plotsr

Chromosome scale *S. cheesmaniae* LA1039, *S. pennellii* LA0716 genomes and publicly available genomes (Supplementary Table 1) were aligned to MbTMV^53^ using Minimap2 2.24-r1122^111^ with options -ax asm5 --eqx, followed by running SyRI v1.6^128^ with default settings and the BAM file as input, to identify and call polymorphisms and structural variations and generate a VCF file per comparison. LA1039 and LA0716 synteny and rearrangement plots were created using plotsr v0.5.4^129^ with default options. LA0716 genomes were compared following similar methods as described above by whole genome alignment and SyRI/plotsr.

For pairwise genome comparison, step-wise alignments were generated following the aforementioned method whereby each query genome was used as reference for the subsequent alignment, followed by generating synteny and rearrangement plots using plotsr v0.5.4^129^ with options -H <1-9> and -W <1-9> and –genomes.txt.

### Repeat identification using two distinct algorithms

The repetitive regions of *S.lycopersicum*-MbTMV, *S. cheesmaniae* and *S. pennellii* genomes were detected using two distinct methods. First, the mathematical Context Tree Weighting (CTW) algorithm^64^ from the Bayesian Context Trees (BCT) R package^130^ was used to estimate per-base sequence probabilities based on variable-length nucleotide contexts. Regions with low CTW probabilities indicate high sequence predictability, allowing the identification of biologically relevant repetitive or low-complexity regions without prior alignment or annotation. The second approach was to detect the repeat regions of those two genomes utilising the Tandem Repeat Annotation and Structural Hierarchy (TRASH) pipeline^65^. TRASH also annotates tandem repeats within the genomic sequences and organises them into a hierarchical structure, enabling the identification of complex or nested repeat architectures^65^. The CTW probabilities and the abundance of the detected monomeric repeats from TRASH were calculated in 100 Kbp windows and plotted using the R package circlize v.0.4.16^131^.

### Transposable Element Annotation and Lineage Classification

The transposable elements of all species included in this study (Supplementary Table 1) were identified using the Extensive De novo TE Annotator pipeline v.2.0.1^132^ with the parameters — -anno 1 –cds. For the five domesticated accessions of tomato, the *S.lyc*-Heinz1706 cds file^7^ was used, to purge the gene coding sequences in the transposable element annotation, while for the *S. tuberosum*, the cds file was retrieved from the same database with the genome^55^. Additionally, for *S. galapagense* and *S. habrochaites* the cds files were obtained from a different publication from their genomes, which included the cds file for those two species^13^. For *S. cheesmaniae* the cds file was created using liftoff v1.6.1 to liftover ITAG4.0 gene annotation^7^ and non-reference gene annotation of the *Lycopersicon* clade^6^ to the *S. cheesmaniae* genome with default settings, while for *S. pennellii* the cds file from Bolger et al. was used^3^. The summary output of each species was used to create the genomic percentage table (Table 2). Intact and fragmented *Ty1/Copia* and *Ty3* LTR retrotransposons were further classified into lineages using TEsorter v.1.4.6^133^ with parameters -db rexdb-plant. For the downstream analysis, all LTR retrotransposons classified into different superfamilies using Extensive De novo TE Annotator and TEsorter (for example, an element needed to be classified as *Ty1/Copia* by both pipelines) were excluded.

### Age of LTR Retrotransposons and Phylogenetic Analysis of *Tekay* Elements

The age of the LTR retrotransposons was estimated based on the sequence divergence between the two LTRs of an intact element. This LTR identity metric is included in the EDTA outputs. By using the LTR identity, we were able to calculate the age of the LTR retrotransposons by utilising the following formula:

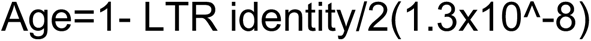

where 1.3x10^-8 is the mutation rate in tomato and 2 is the number of different sequences in which the mutation rate is applied and in this case are the two LTRs of the LTR retrotransposon.

For the phylogenetic analysis, we retained *Tekay* elements that contained all six genes (gag, protease, reverse transcriptase, RNaseH, integrase, and chromodomain) in the correct order according to TEsorter. Those elements were further filtered based on statistics (coverage equal to or above 70% and e-value equal to or smaller than 1e-100) of their integrase domains. The amino acid sequences of integrase domains that passed the above filtering were aligned using MAFFT v.7.526^134^ with parameters –-auto. We employed FastTree v.2.1.11^135^ with parameters -wag and -gamma to generate maximum-likelihood trees.

Downstream analysis was conducted and plotted using the R language and environment (R Core Team, 2025), utilising packages included in the Tidyverse collection v.2.0.0^104^, data.table v.1.16.4 (https://r-datatable.com/), ape v.5.8.1^136^, ggtree v.3.10.1^137^, circlize v.0.4.16^131^, and ComplexHeatmap v.2.18.0^138^.

### Recombination analysis

We used the initial list of SNPs detected between parental genomes using HiFi reads. Subsequently, Illumina reads from a MbTMV x S. pennellii F1 plant were used to detect SNPs relative to the MbTMV reference. The Illumina reads of the F1 plant were trimmed using trimgalore (https://github.com/FelixKrueger/TrimGalore) and aligned to Moneyberg-TMV reference genome using BWA-MEM^107^. SNPs were detected from the alignment output using GATK HaplotyperCaller^127^ and were subsequently filtered by performing GATK hard-filtering and retaining only the heterozygous variants. Also, a SNP is excluded if it is found within a region with excessive SNP density (5-kb window; 95^th^ percentile), within or near homopolymers (distance < 5 bp), or having excessive read coverage (95^th^ percentile). SNPs from the F1 plant with an unusual allele frequency ratio (allele_1_ – allele_2_ > 0.8) were used to further filter the *S.pennellii* SNPs, resulting in the final set of markers distinct between the two parental genomes. The same marker detection was performed on the other two hybrid crosses (MbTMV x Micro-Tom, and MbTMV x *S. cheesemaniae*).

BC Illumina data from all populations were aligned to MbTMV^53^ using BWA-MEM^107^ with default parameters followed by converting SAM to BAM and sorting of BAM files using SAMtools v1.9^108^. Duplicates were removed using Sambamba markdup v0.8.1^139^ with the option -r. For each BC sample, bcftools mpileup was used to detect SNPs. Afterwards, we ran the Meiotic CrossOver finder (MeiCOfi, Fuentes et al., in preparation) to detect crossovers with window size = 500kb, step size = 250kb, window count = 6, minimum SNPs = 30.

### Coldspot comparison

We first computed the number of crossovers using a sliding 50Kbp window with 10Kbp step size. All overlapping windows with zero crossovers were merged, and only those with at least 1-Mb total length are reported as coldspots. We excluded telomeric coldspots if the length is below 1.5Mb or if they are from MbxMT male or female populations, where the density of SNPs is low, and detection of telomeric crossovers is inaccurate. The resulting coldspot locations were compared to the SV annotations to determine the putative crossover suppressor, such as large inversions.

We ran hometools mapbp (https://github.com/mnshgl0110/hometools) on the previously generated Syri output to convert the coldspot coordinates from MbTMV to the other three parental genomes. If hometools cannot map the locations, we convert the location of the syntenic region closest to the coldspots boundaries. After generating the coldspot coordinate mapping, we compared the relative lengths of the coldspots and estimated the genomic expansion or contraction. We also used the TE and gene annotations of each parental genome to compute the difference in TE composition and gene count within the coldspot regions.

## Data availability

Raw HiFi, Omni-C and ONT sequencing data has been uploaded to the European Nucleotide Archive (ENA) under project codes: PRJEB62444 (*S. cheesmaniae* LA1039) and PRJEB62445 (*S. pennellii* LA0716). Illumina short-read sequencing data is available at the ENA for *S. lycopersicum* Moneyberg-TMV x *S. lycopersicum* Micro-Tom backcross populations (PRJEB101732), *S. lycopersicum* Moneyberg-TMV x *S. cheesmaniae* backcross populations (PRJEB101733) and *S. lycopersicum* Moneyberg-TMV x *S. pennellii* backcross populations (PRJEB101734). The *S. lycopersicum* Moneyberg-TMV Isoseq and *S. pennellii* LA0716 Isoseq data has been uploaded to the ENA under project codes PRJEB104324. Genomic variants between *S. lycopersicum* cv Moneyberg-TMV and *S. lycopersicum* cv. Micro-Tom or *S. cheesmaniae* LA1039 or *S. pennellii* LA0716 as identified by SyRI are freely available in VCF format upon request to the corresponding author.

## Acknowledgements

1. *S. cheesmaniae* LA1039 and *S. pennellii* LA0716 seeds were obtained from TGRC (UC Davis, California, USA). Moneyberg-TMV seeds were kindly provided for research purposes by Cilia Lelivelt and Maarten Verlaan (Rijk Zwaan, Netherlands). S. lycopersicum cv Micro-Tom seeds were originally sourced from Tomato Growers Supply company (Ft. Myers, Florida, USA). We thank Andre Marques for his contributions on the interpretation of Hi-C contact plots with regards to genome assemblies. The authors thank Manish Goel for his help with run-case scenarios in whole genome comparison. Finally, we thank Marianne Harperscheidt and Christine Saenger for their help in with maintenance of plant materials.

## Author contributions

CJU and WMJR designed the study. SE maintained all plant materials with contributions of WMJR and sampled plant materials of backcross populations. SE performed hybridization with contributions of WMJR. BH performed library preparation and sequencing of backcross populations. BH performed high molecular weight DNA isolation and library preparation for Omni-C and HiFi sequencing. ZZ and BU performed ONT library preparation and ONT sequencing. WMJR generated the LA1039 genome assembly. ZZ generated the mixed ONT and HiFi LA0716 assembly. WMJR scaffolded the LA1039 and LA0716 genome assemblies and performed validation analyses with contributions of CJU, RRF, EP, ZZ and BU. WMJR performed pairwise comparison of genome assemblies and sequencing data with contributions of CJU, RRF and EP. Iso-seq data for Moneyberg-TMV was generated by YW, WMJR and BH. Iso-seq data for LA0716 was generated by SA and analysed by ZZ. Gene annotation was performed by ZZ and BU. WMJR characterized genomic variation with contributions of RRF, EP and CJU. EP performed the repeat content and transposable element analysis. RRF performed crossover analysis with contributions from WMJR, JF, TS, CJU and QL. WMJR, RRF, EP and CJU wrote the manuscript and generated the figures with contributions of all authors.

