## Supplementary Figures 1-19 for "Chromosome-scale *Solanum pennellii* and *Solanum cheesmaniae* genome assemblies reveal structural variants, repeat content and recombination barriers of the tomato clade"

### Supplementary Fig. 1

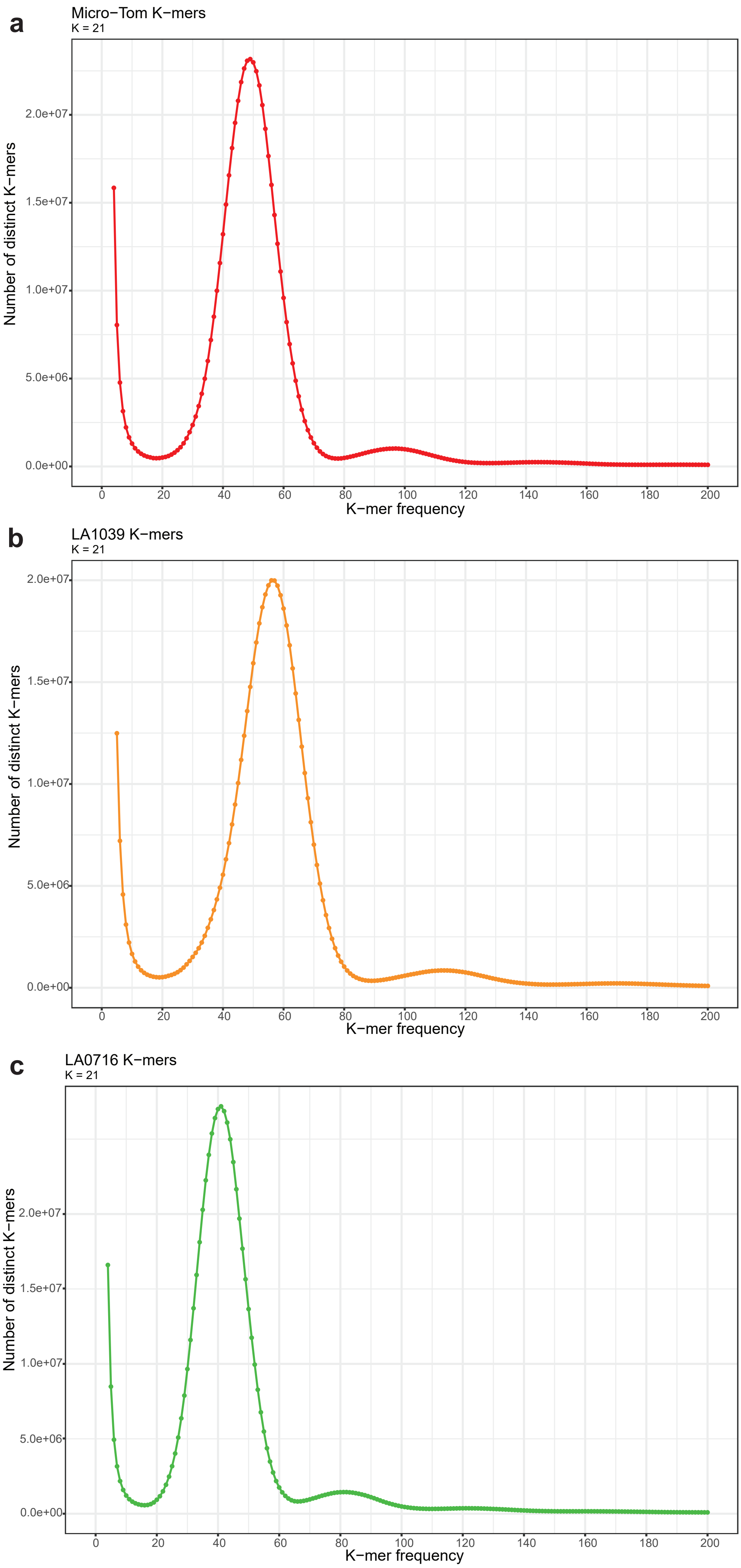


K-mer Analysis Toolkit (KAT) results of the K-mer distribution in Micro-Tom, LA1039 and LA0716 HiFi data.
**a** *S. lycopersicum* cv. Micro-Tom HiFi K-mer frequency distribution where K = 21. **b** *S. cheesmaniae* LA1039 HiFi K-mer frequency distribution where K = 21. **c** *S. pennellii* LA0716 HiFi K-mer frequency distribution where K = 21.

#### Supplementary Fig. 2

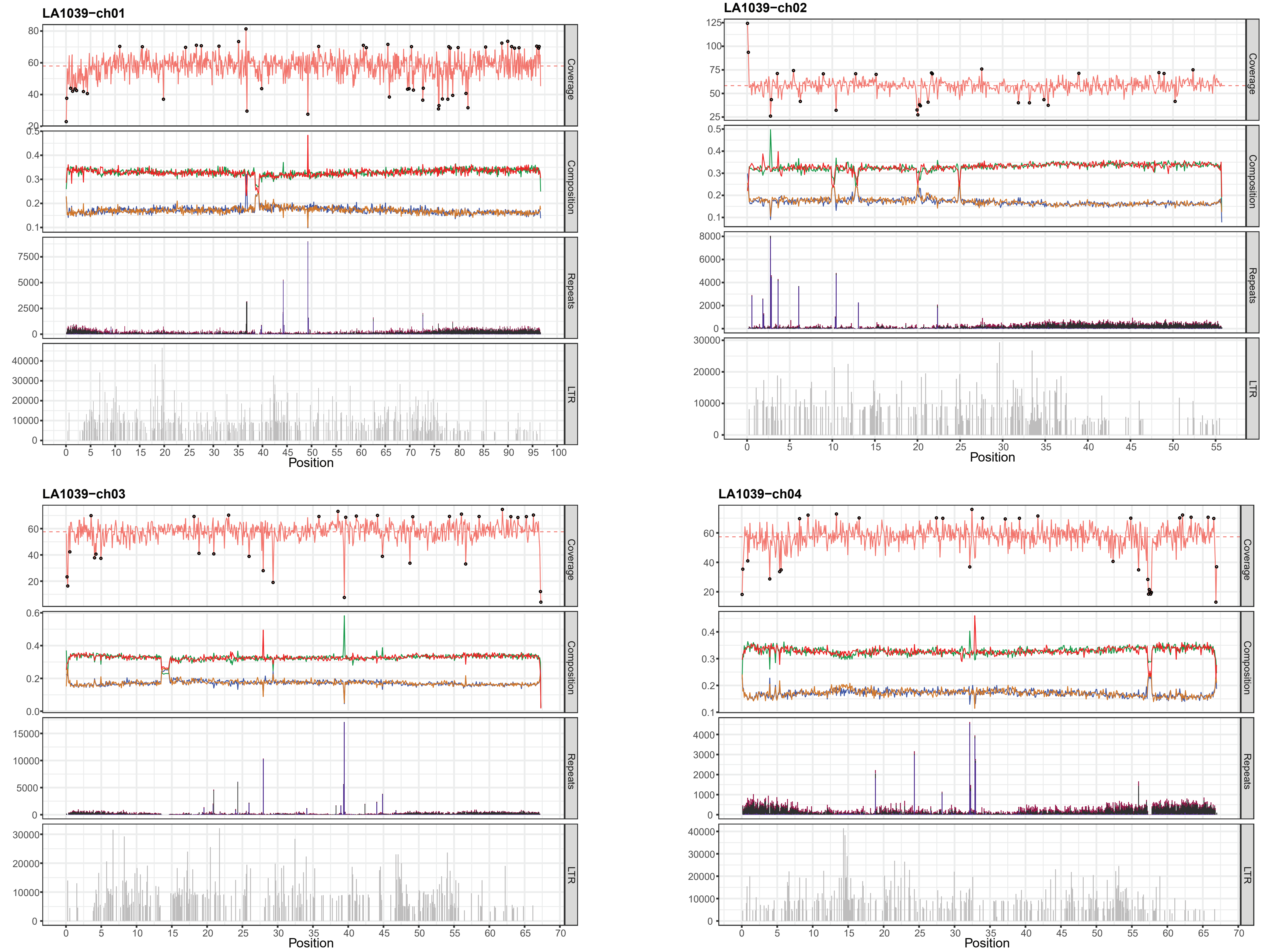

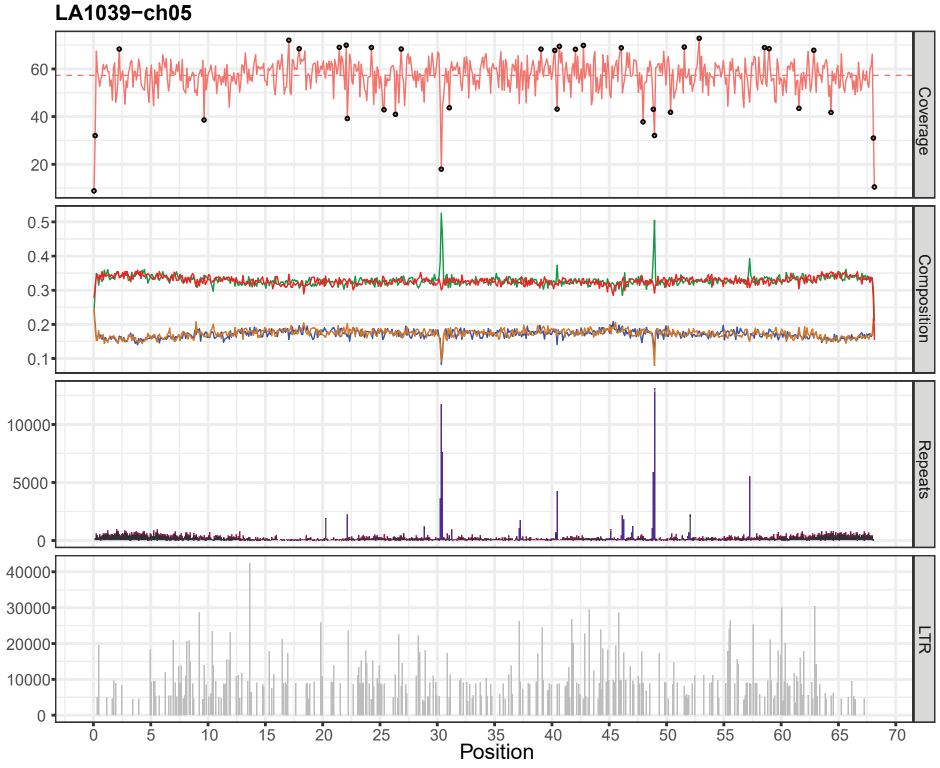
LA1039−ch05
Coverage
Composition
Repeats
LTR
Position

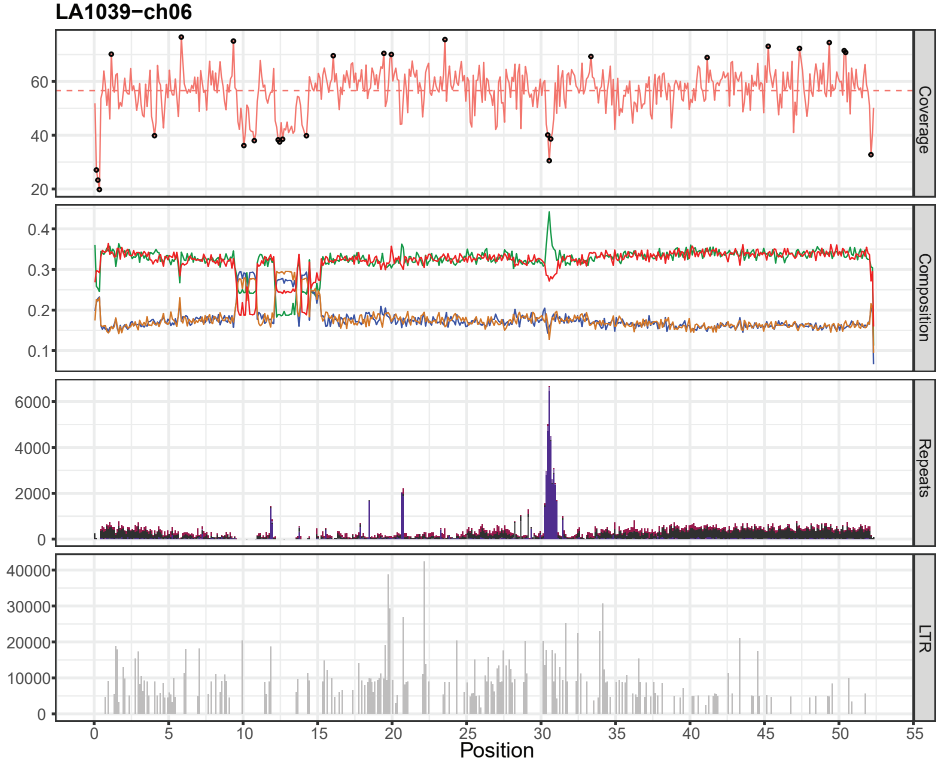
LA1039−ch06
Coverage
Composition
Repeats
LTR
Position

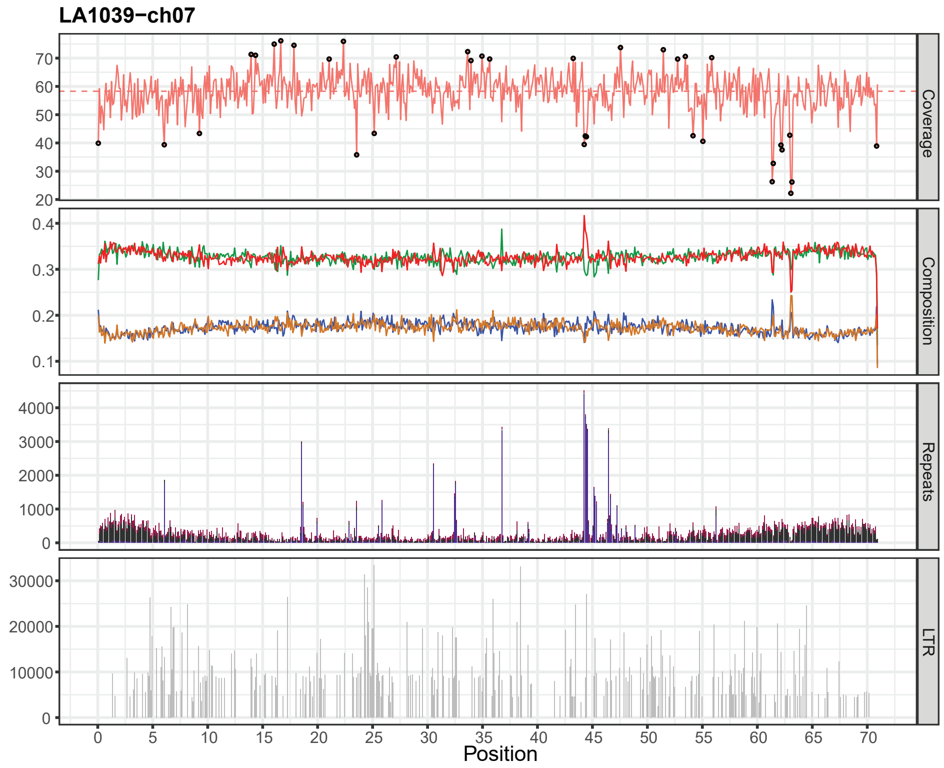
LA1039−ch07
Coverage
Composition
Repeats
LTR
Position

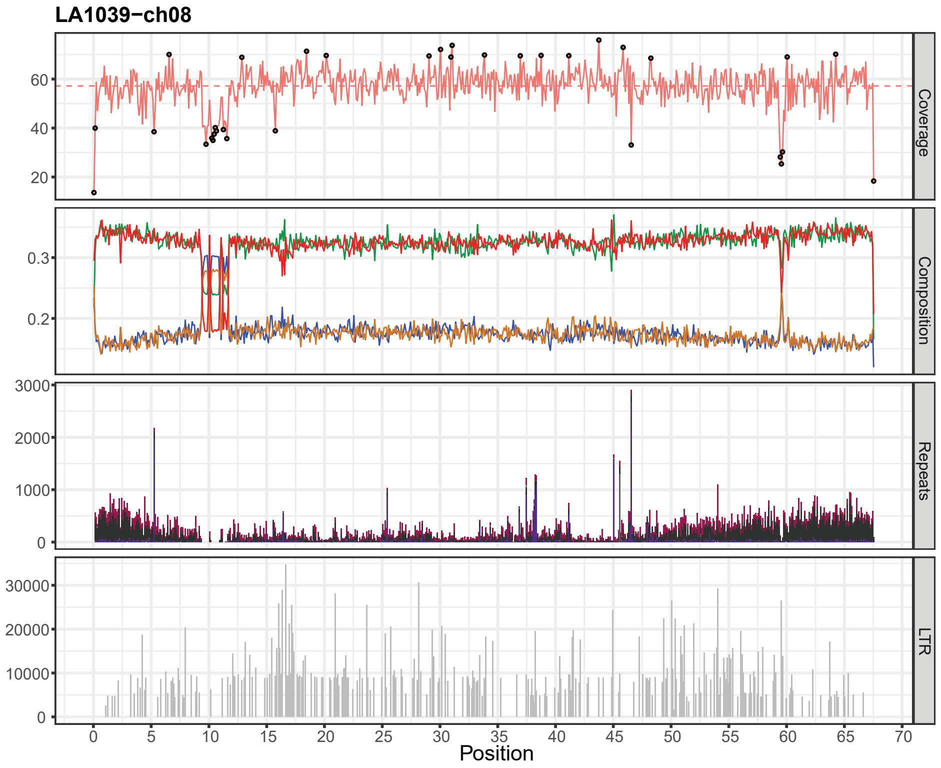
LA1039−ch08
Coverage
Composition
Repeats
LTR
Position

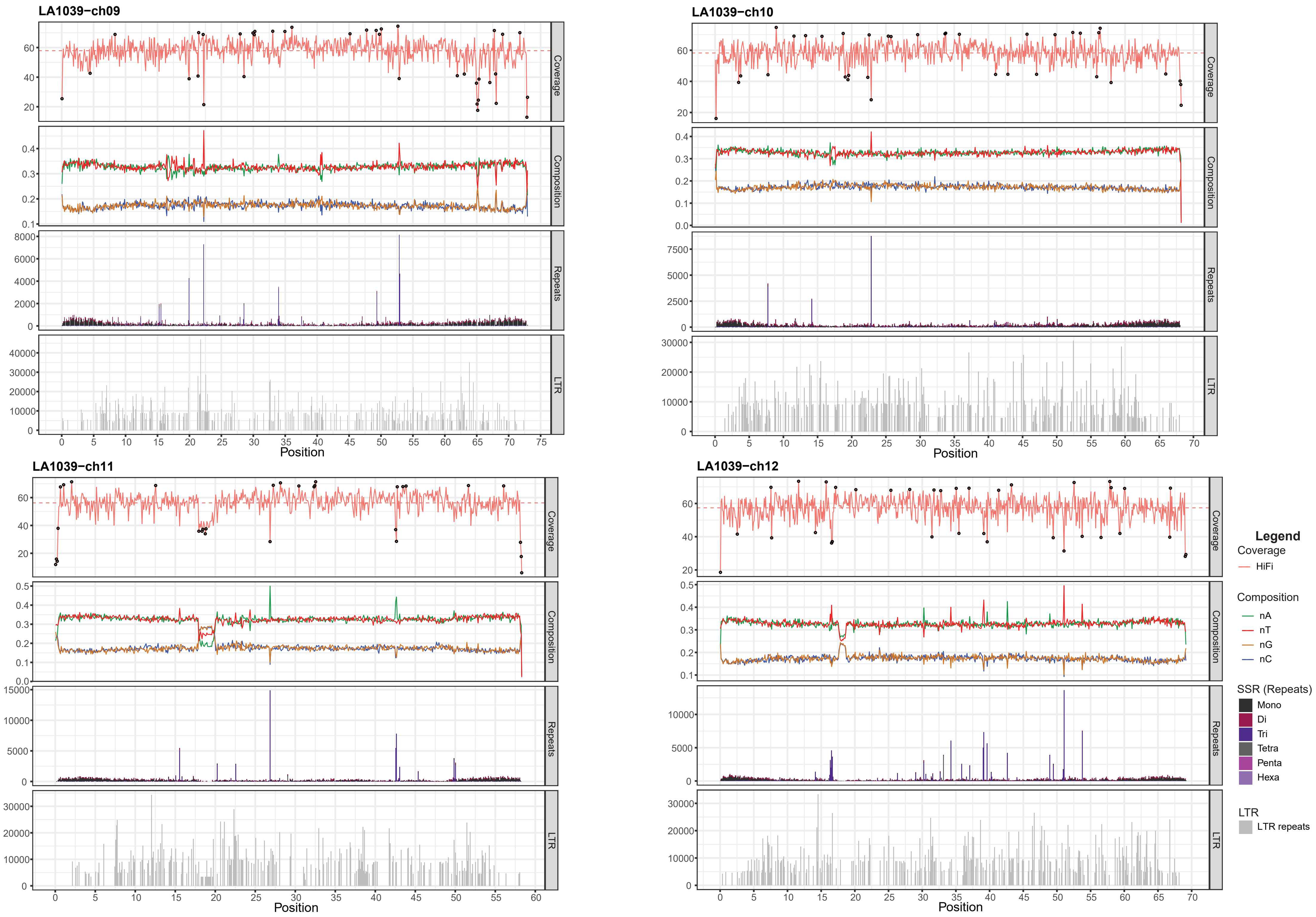


Characterization and validation of the 12 *S. cheesmaniae* LA1039 chromosomes include primary read coverage analysis (Coverage panel), nucleotide composition ratios (Composition panel), SSR Repeat analysis (Repeats panel) and LTR repeat analysis (LTR panel) in 100kb windows plotted over the genomic position on the genome. Coverage panel: dashed horizontal line represents mean coverage. Black circles represent coverage outliers (<2.5% and >97.5% percentiles). X-axis is chromosome position in megabasepair.

#### Supplementary Fig. 3

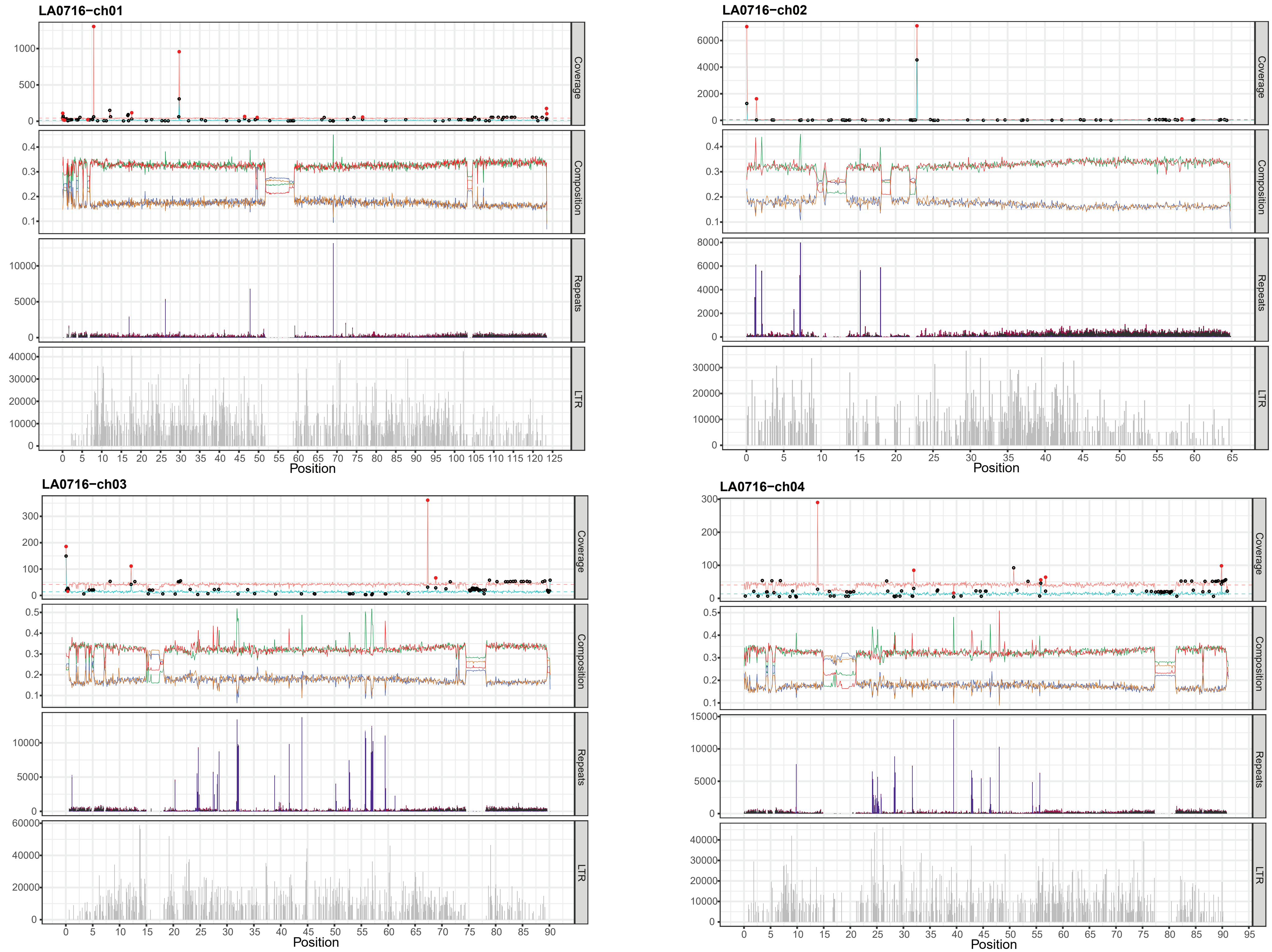

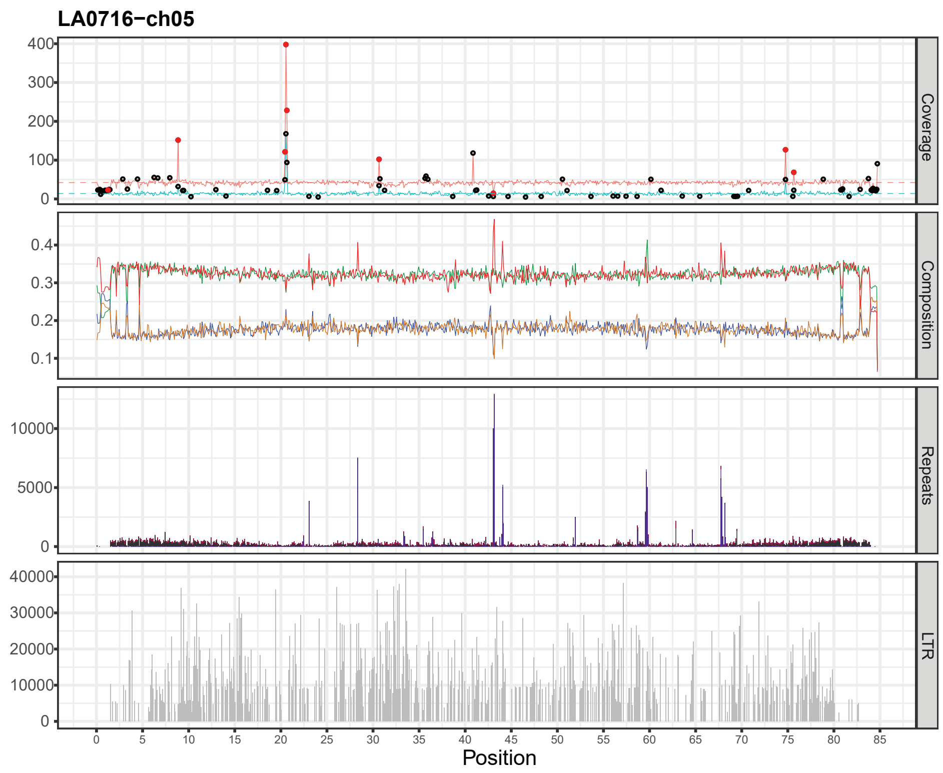
LA0716−ch05
Coverage
Composition
Repeats
LTR
Position

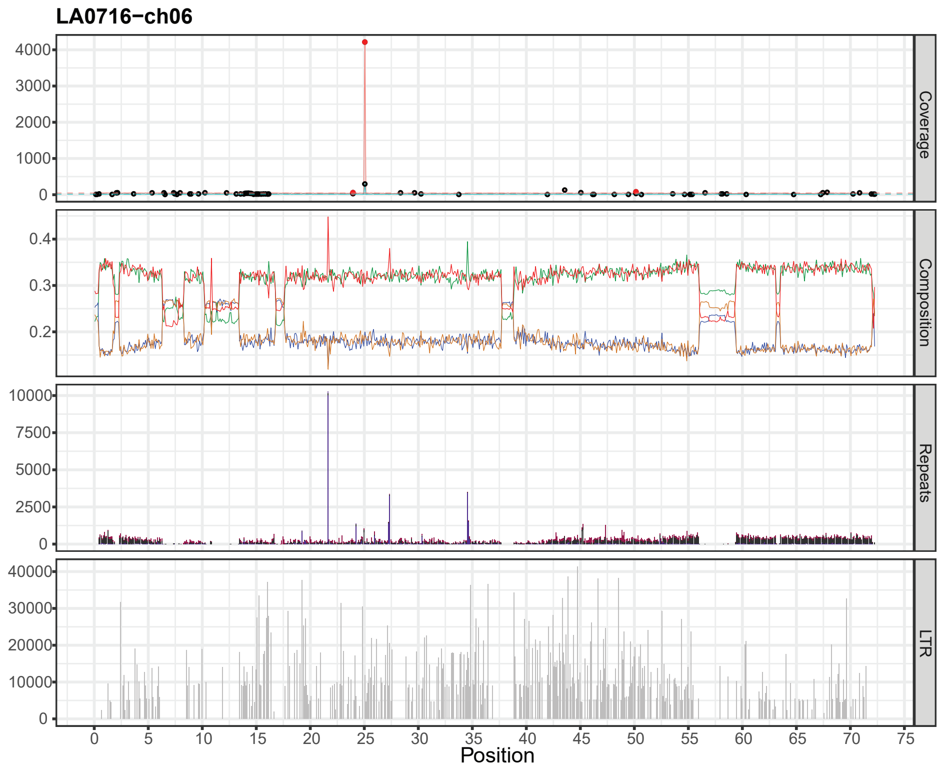
LA0716−ch06
Coverage
Composition
Repeats
LTR
Position

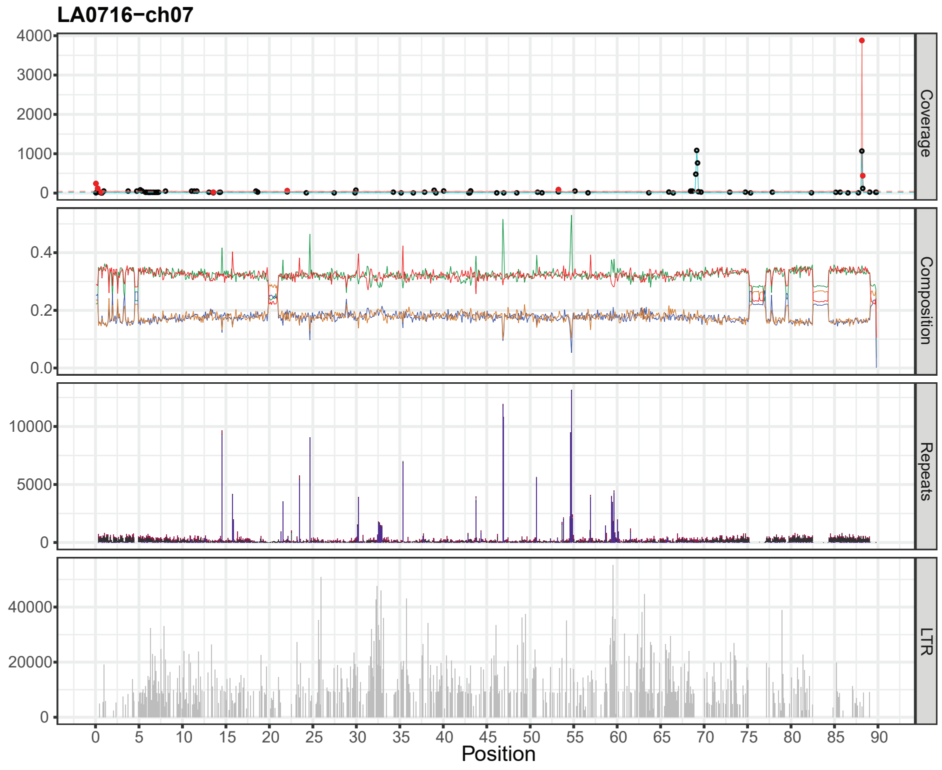
LA0716−ch07
Coverage
Composition
Repeats
LTR
Position

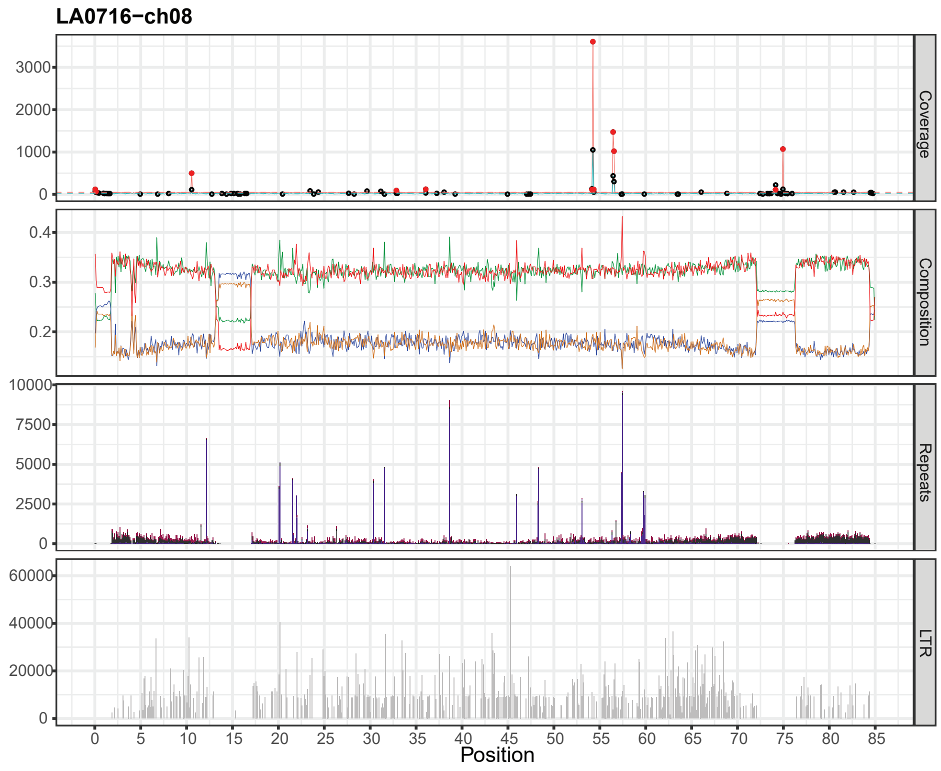
LA0716−ch08
Coverage
Composition
Repeats
LTR
Position

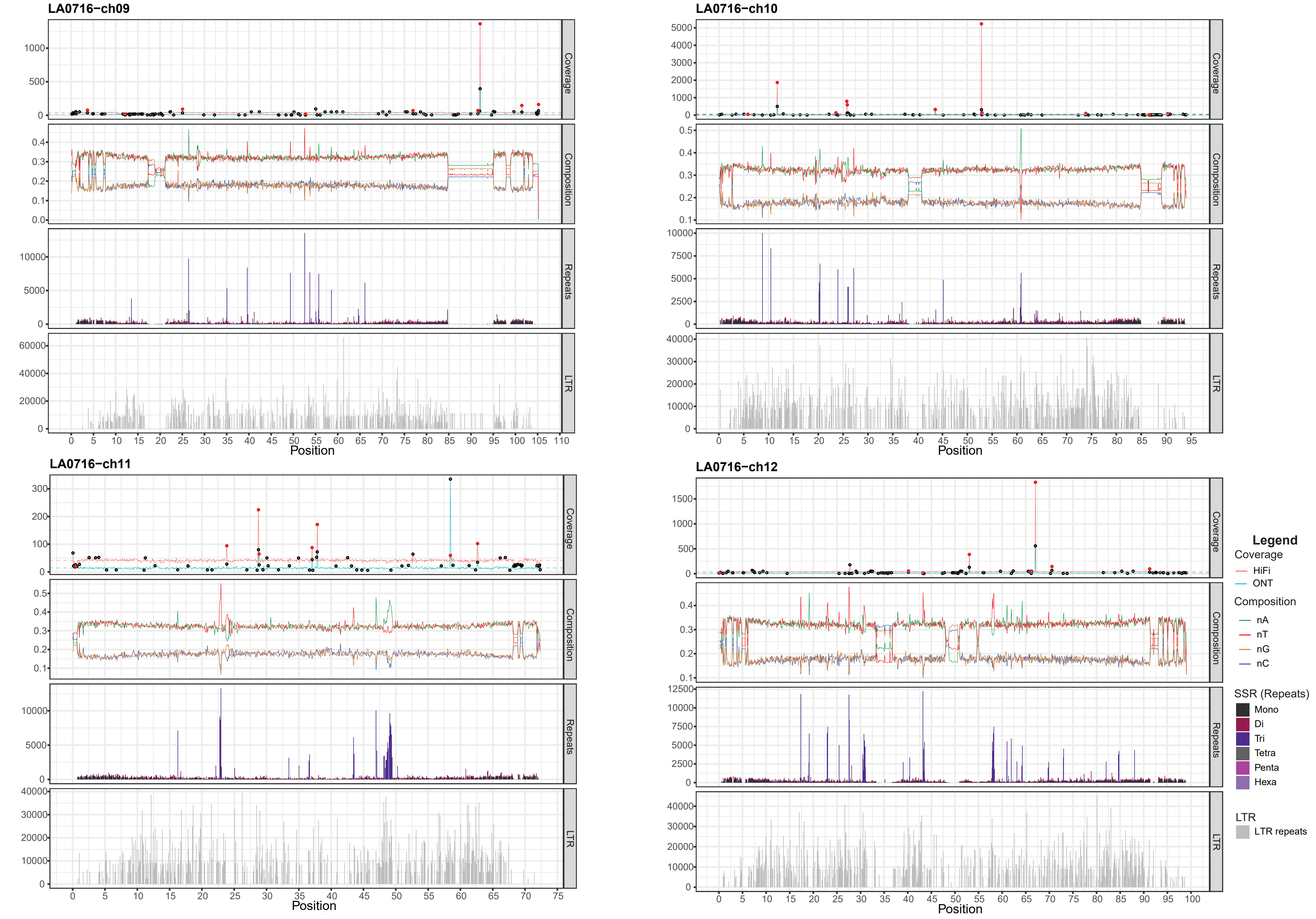


Characterization and validation of the 12 *S. pennellii* LA0716 chromosomes including primary read coverage analysis (Coverage panel), nucleotide composition ratios (Composition panel), SSR Repeat analysis (Repeats panel) and LTR repeat analysis (LTR panel) in 100kb windows plotted over the genomic position on the genome. Coverage panel: dashed horizontal line represents mean coverage. Black circles represent coverage outliers (<2.5% and >97.5% percentiles), red circles represent shared coverage outliers. X-axis is chromosome position in megabasepair.

### Supplementary Fig. 4

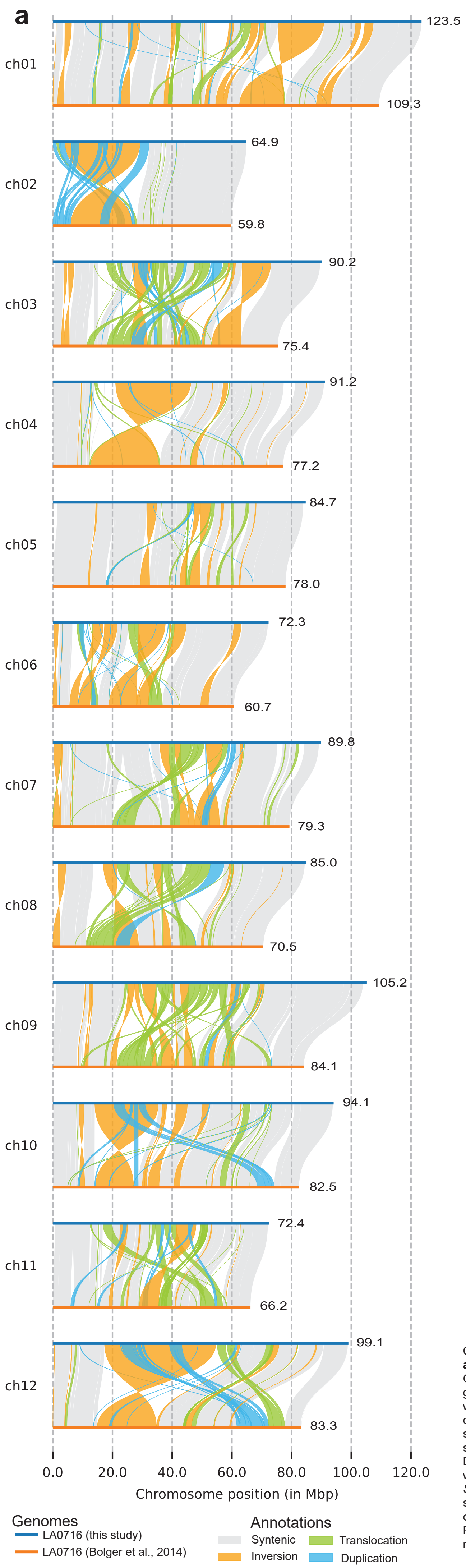


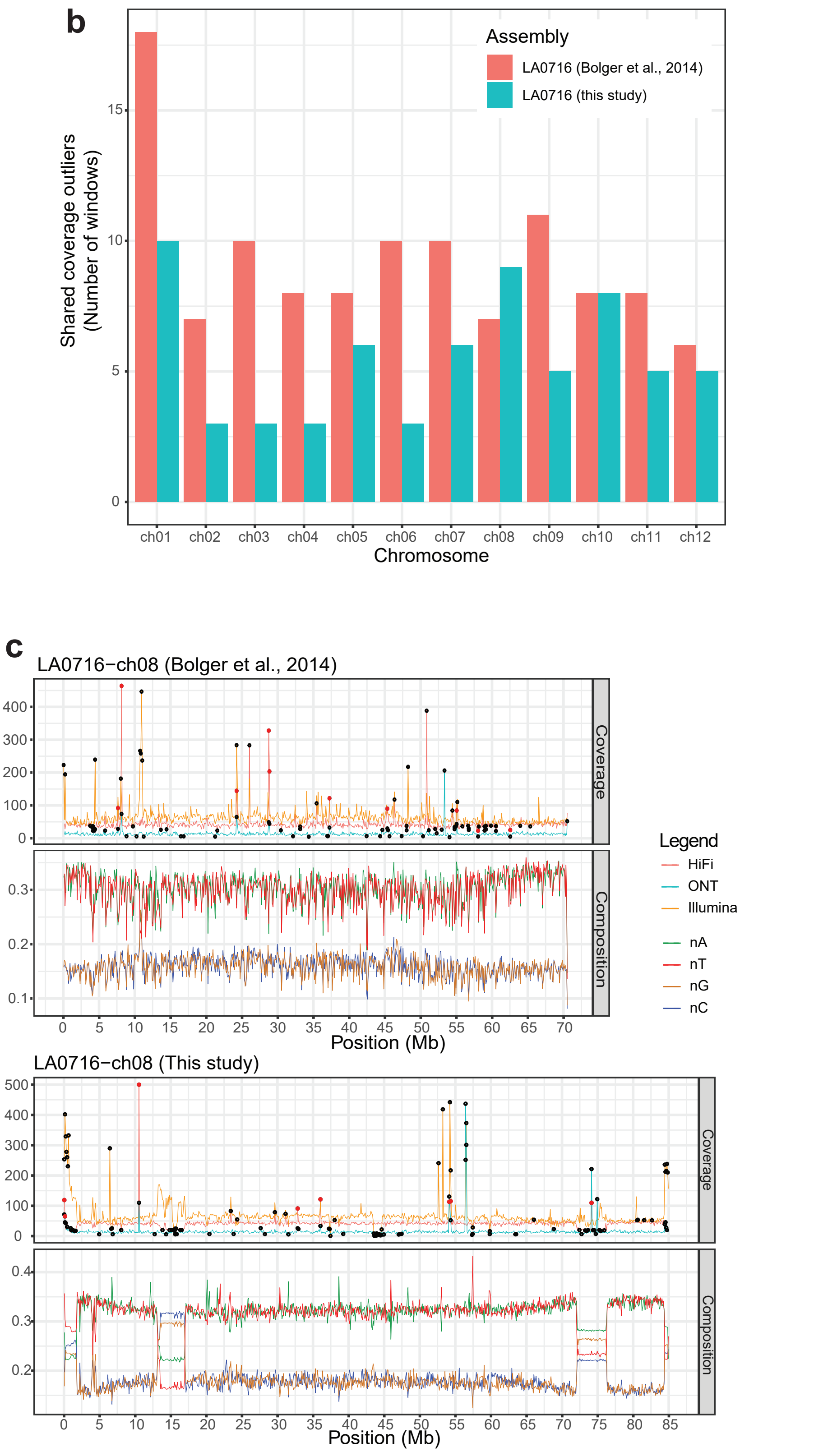


Comparison of original and new *S. pennellii* LA0716 genome assemblies
**a** Synteny and Rearangement plot (SyRI) between LA0716 (this study) and the 2014 LA0716 genome (Bolger et al., 2014). Genome tracks are LA0716 (this study) in blue and the 2014 LA0716 genome in orange. Annotations include syntenic regions in grey, inversions in yellow, translocations in green and duplications in blue. Non-aligned or non-syntenic regions are visualized by white gaps and can include insertions and deletions. Increased chromosome length is observed for all LA0716 (this study) chromosomes. Large genomic variation between the genomes is observed especially in pericentromeric regions and also at the start of chromosomes 6, 7 and 8. **b** Pairwise analysis of both *S. pennellii* assemblies for deviating read coverage in 100kb windows showing the number of shared windows between HiFi (this study), ONT (this study) and Illumina (Bolger et al., 2014) data. Deviating coverage is concidered coverage (<2.5% percentile and >97.5% percentile). Equal or increased numbers of windows with shared deviating coverage are observed 2014 LA0716 genome for all chromosomes except ch08. **c** Pairwise analysis of both *S. pennellii* assemblies for sequencing coverage and nucleotide composition. Sequencing coverage of HiFi (this study), ONT (this study) and Illumina (Bolger et al., 2014) data in 100kb windows. Read coverage is supportive of both assemblies. Characteristic changes in nucleotide composition patterns are lacking within the 2014 LA0716 genome, although supported by Illumina coverage. Read coverage was maximized at 500. Black circles represent coverage outliers (whereby coverage <2.5% or >97.5% percentile), red circles represent shared coverage outliers between 2 or more datasets annotated on the HiFi track.

### Supplementary Fig. 5

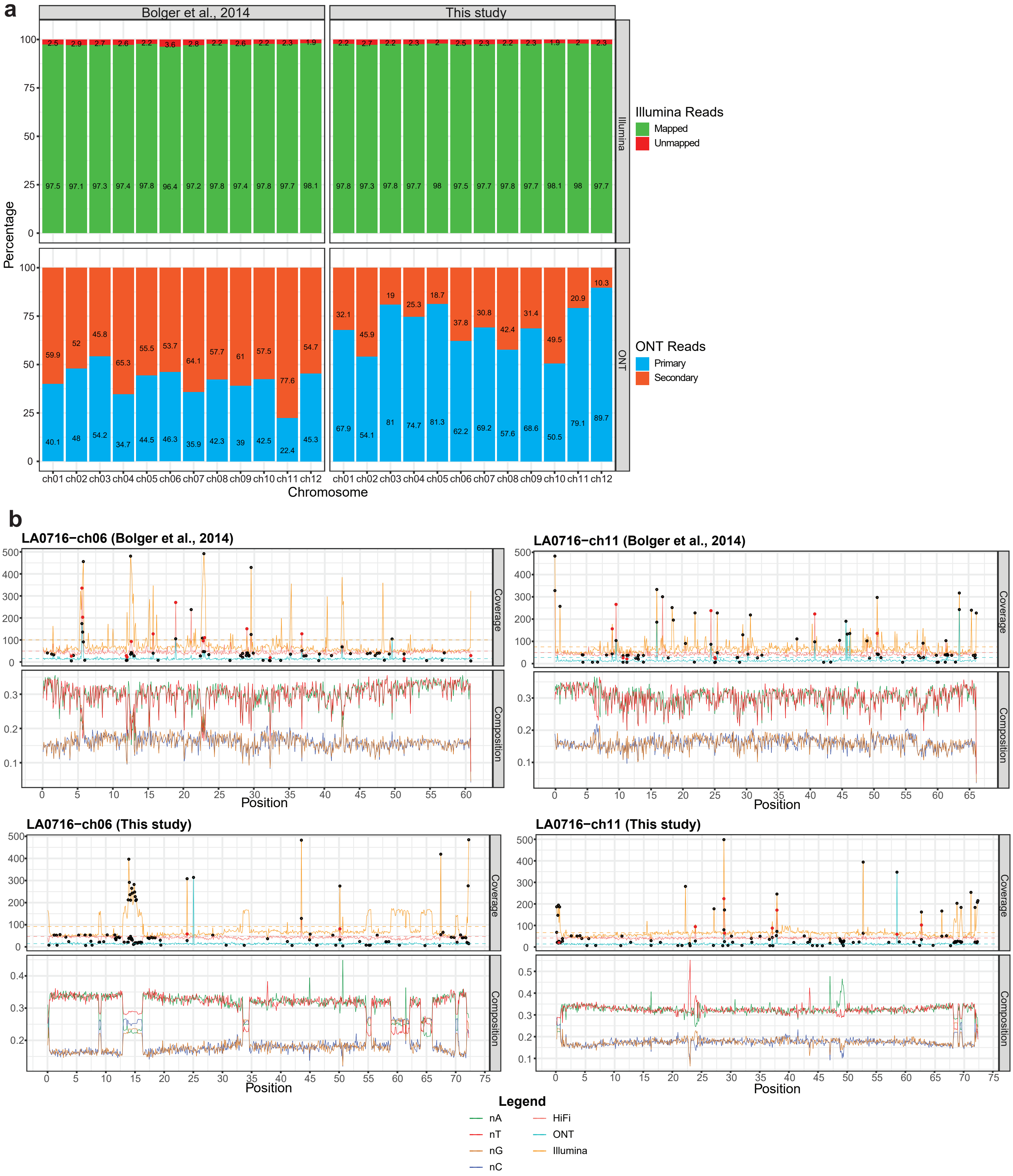


**a** Pairwise sequencing data (Illumina and ONT) analysis. For Illumina data (Bolger et al., 2014), percentage of mapped (green) and unmapped (red) reads to respective LA0716 genomes (this study or Bolger et al., 2014) is shown. For ONT data (This study), percentage of primary (blue) and secondary (orange) aligned reads to respecitve LA0716 genomes is shown. Increased numbers of unmapped and secondary read alignments are shown for the 2014 LA0716 genome (Bolger et al., 2014). **b** Pairwise sequencing coverage comparison of HiFi (this study), ONT (this study) and Illumina (Bolger et al., 2014) data aligned agains the two LA0716 genomes. Read coverage is supportive of both assemblies. Characteristic changes in nucleotide composition patterns are lacking within the 2014 LA0716 genome, although supported by Illumina coverage in our LA0716 genome. Black circles represent coverage outliers (whereby coverage <2.5% or >97.5% percentile), red circles represent shared coverage outliers between 2 or more datasets annotated on the HiFi track.

### Supplementary Fig. 6

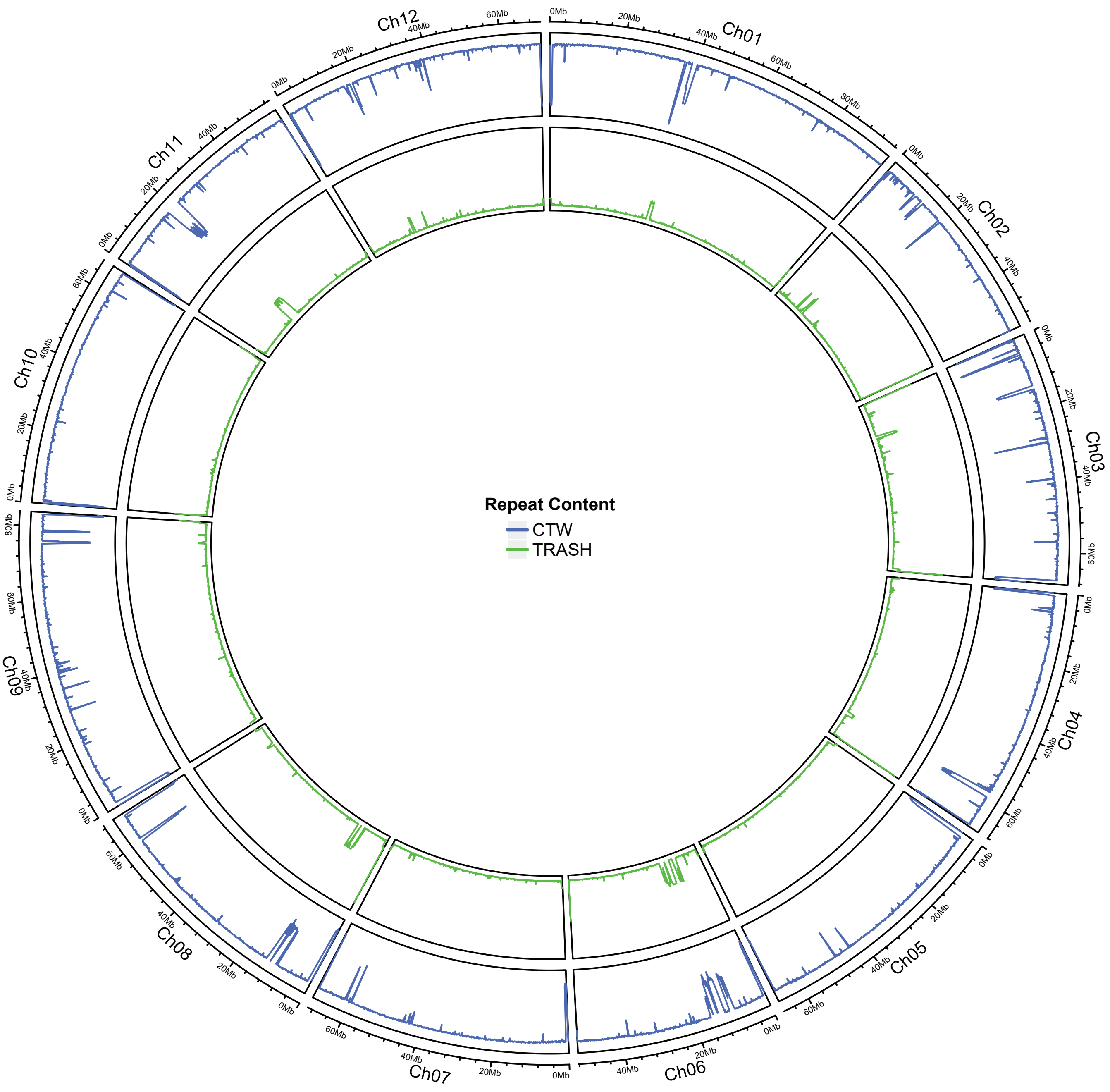


MbTMV repeat content. From outer to inner layer (in 100 Kbp windows): The log-probability of the CTW algorithm (Kontoyiannis et al., 2022) (blue line) and the abundance of monomeric repeats using TRASH (Wlodzimierz et al., 2023) (green line) for S.lycopersicum

### Supplementary Fig. 7

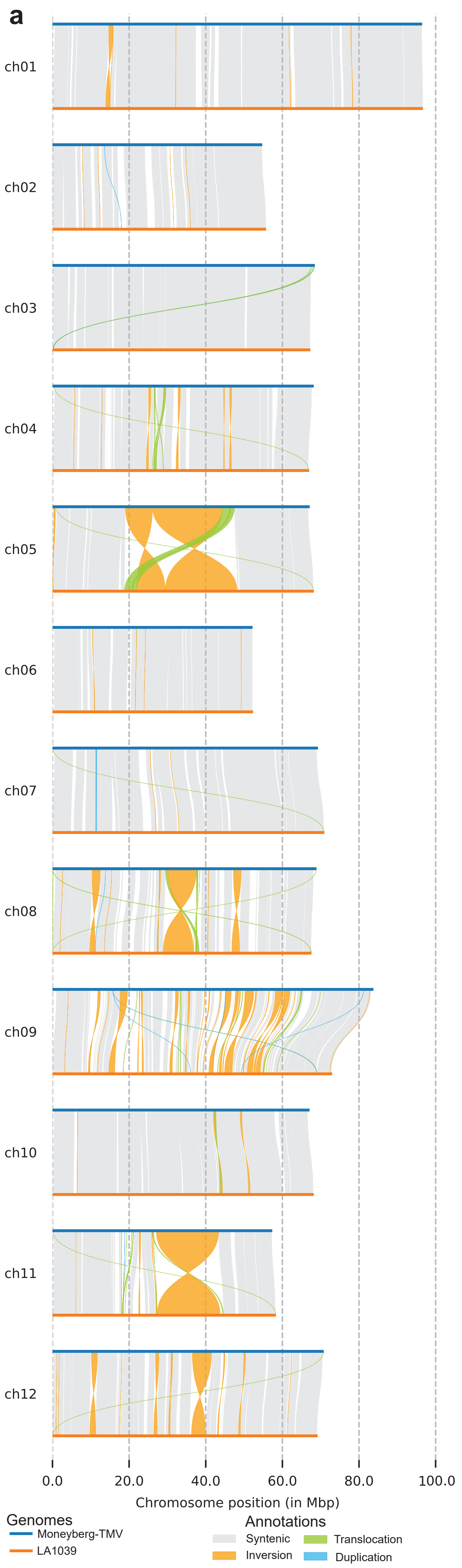


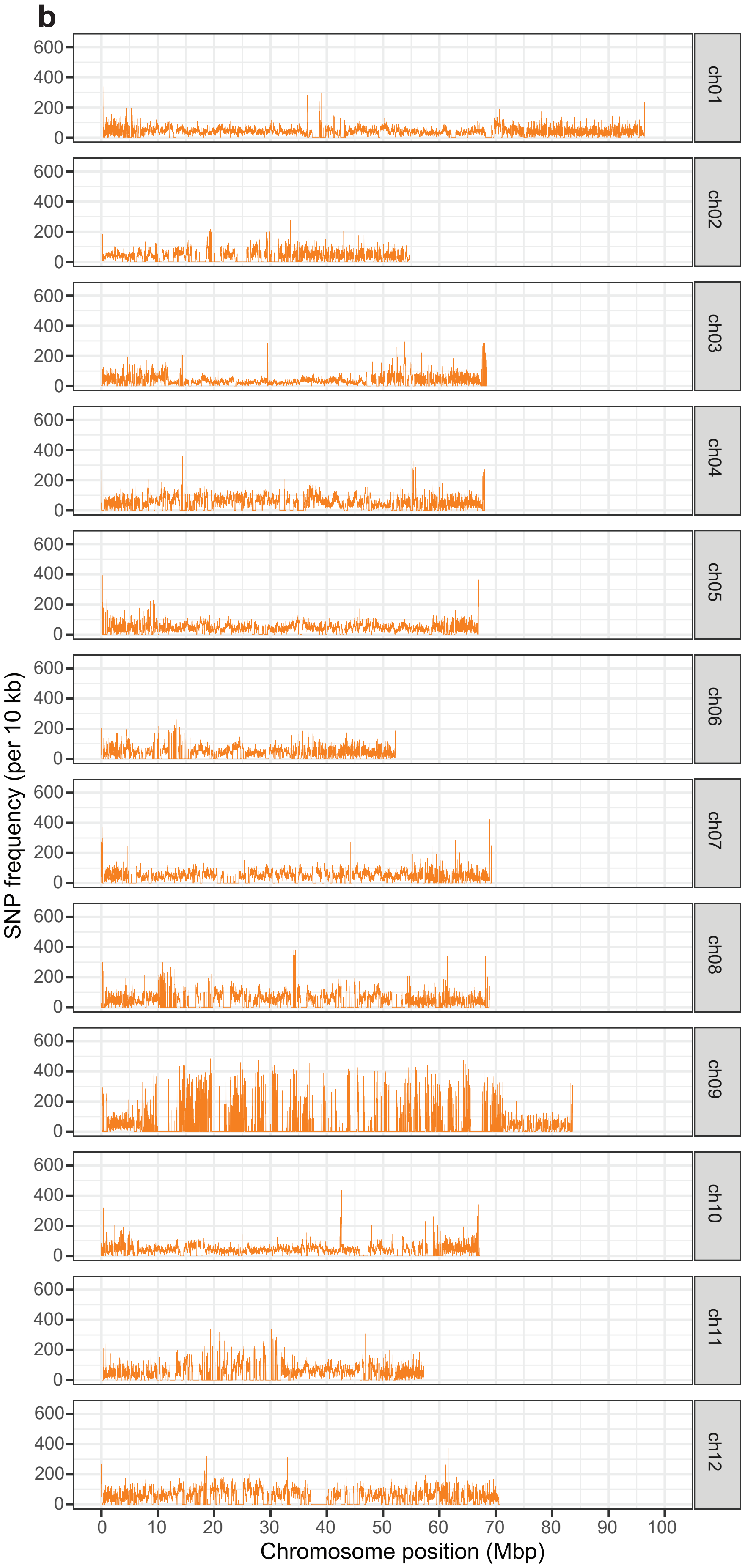


Genomic variation between Moneyberg-TMV (MbTMV) and LA1039 as identified by Synteny and Rearrangement identifier (SyRI)
**a** Synteny and Rearrangement plot (SyRI) between *S. lycopersicum* cv. Moneyberg-TMV and *S. cheesmaniae* LA1039. on-aligned or non-syntenic regions are visualized as white gaps and can include insertions and deletions.
**b** LA1039 specific SNPs identified by SyRI comparison to Moneyberg-TMV, plotted in 10kb windows over the genomic position on the Moneyberg-TMV genome.

#### Supplementary Fig. 8

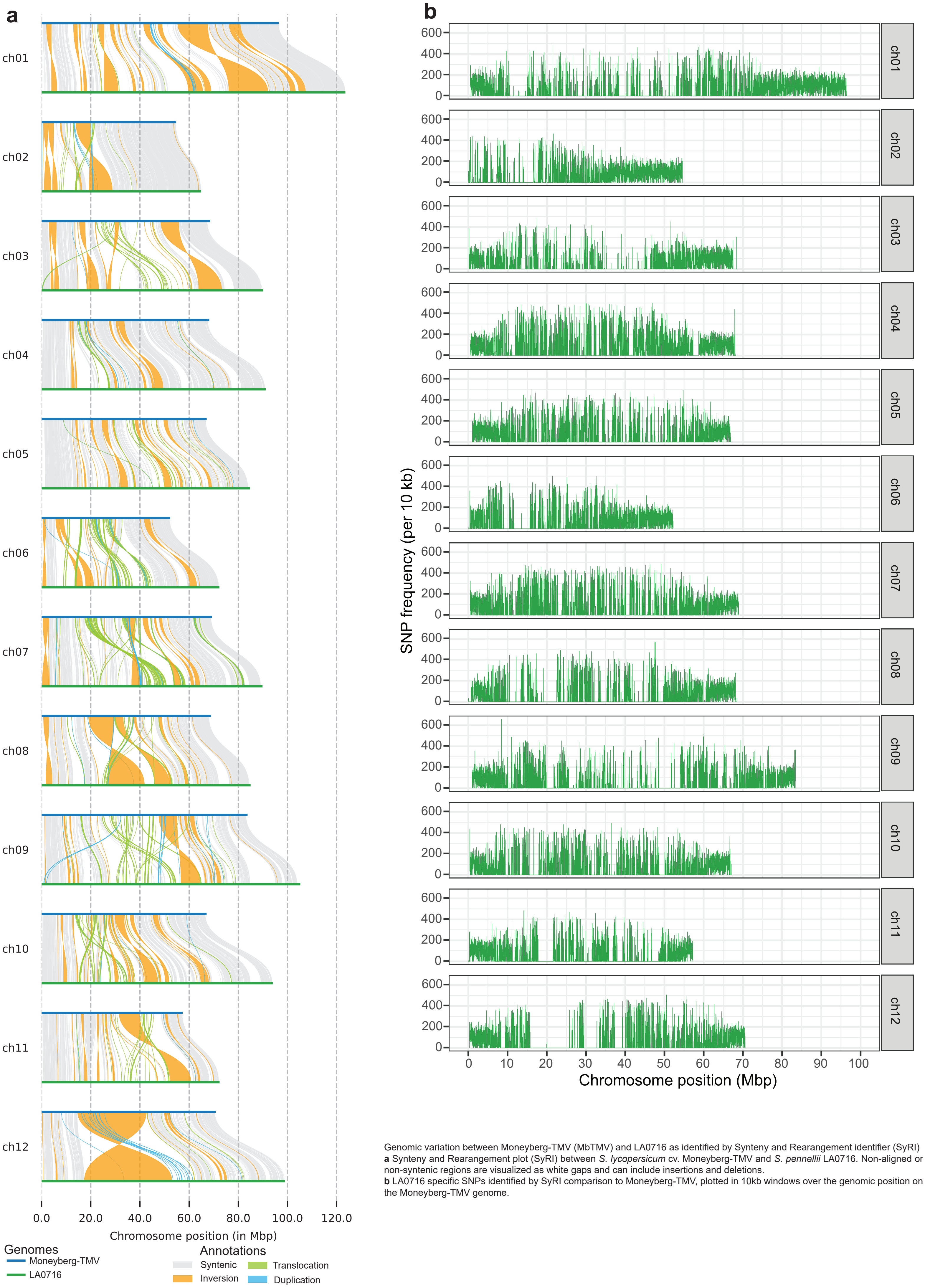


Genomic variation between Moneyberg-TMV (MbTMV) and LA0716 as identified by Synteny and Rearangement identifier (SyRI)
**a** Synteny and Rearangement plot (SyRI) between *S. lycopersicum* cv. Moneyberg-TMV and *S. pennellii* LA0716. Non-aligned or non-syntenic regions are visualized as white gaps and can include insertions and deletions.
**b** LA0716 specific SNPs identified by SyRI comparison to Moneyberg-TMV, plotted in 10kb windows over the genomic position on the Moneyberg-TMV genome.

### Supplementary Fig. 9

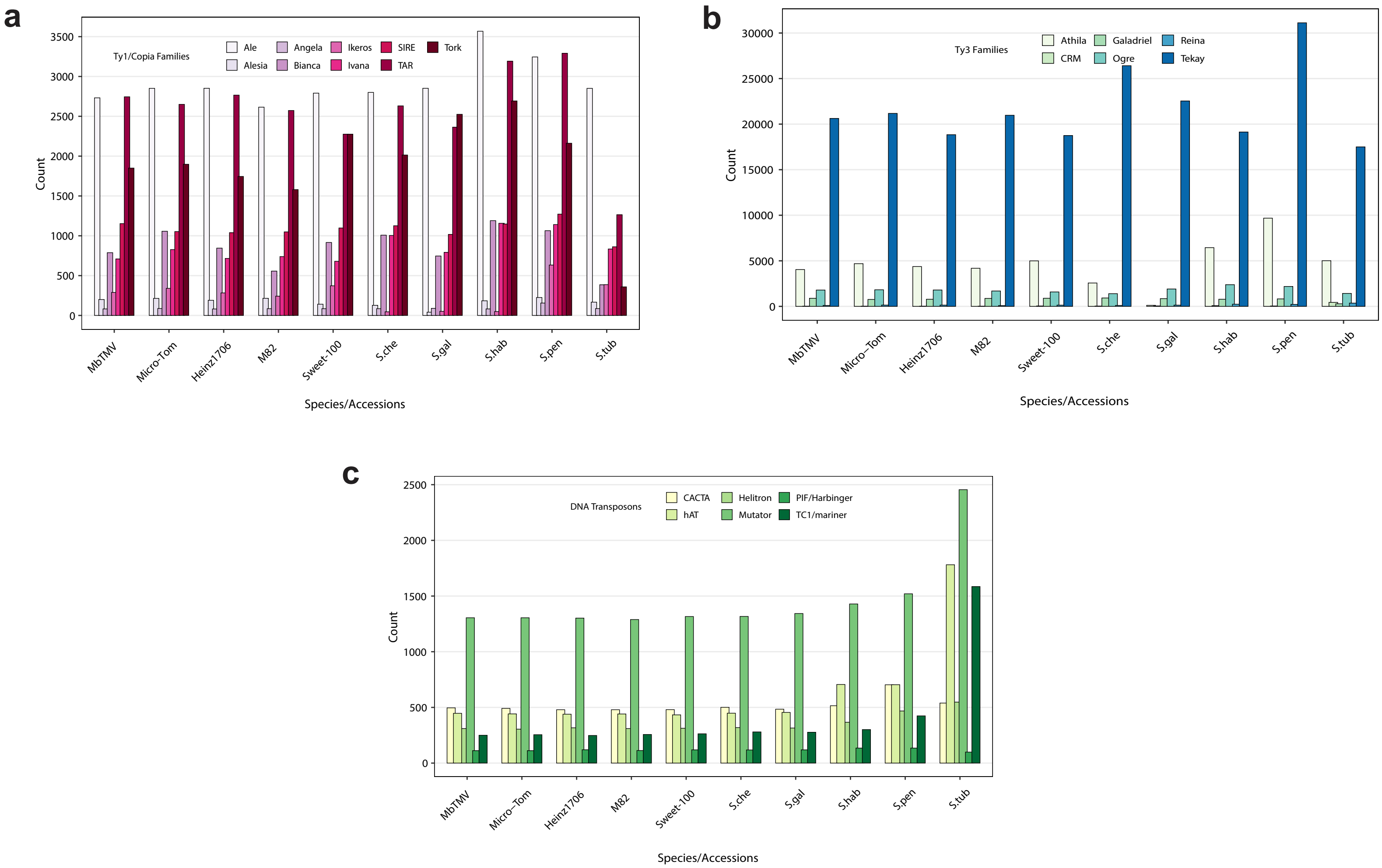


Abundance of LTR retrotransposon fragments and intact DNA transposons of the tomato pangenome. **a** and **b** Quantification of intact elements and high-quality LTR retrotransposon fragments of the main *Ty1/Copia* and *Ty3* lineages. **c** Abundance of the intact DNA transposon elements of each lineage.

### Supplementary Fig. 10

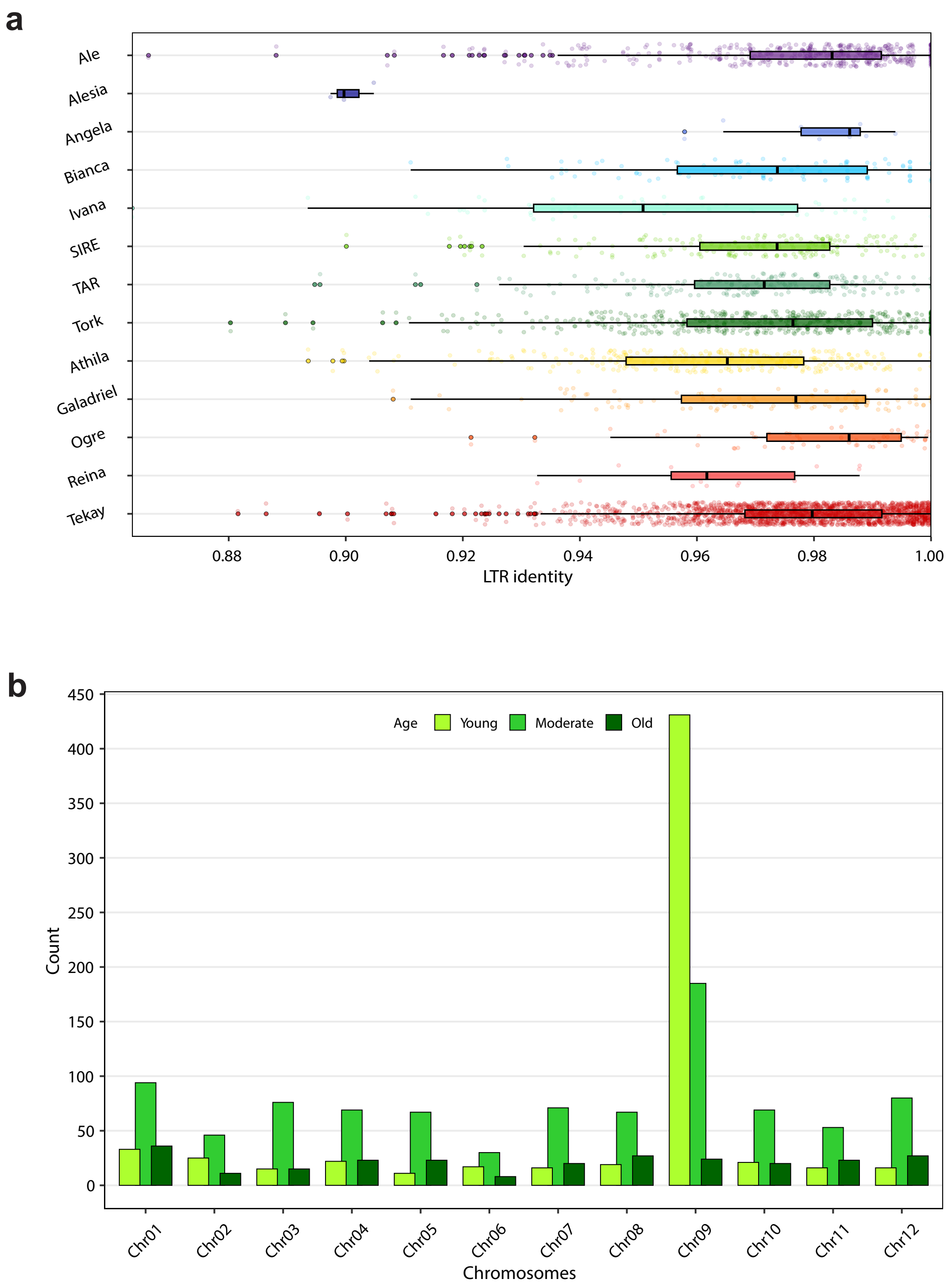


LTR identity of LTR retrotransposon lineages in MbTMV and position of Tekay elements on MbTMV chromosomes. **a** LTR identity of the main LTR retrotransposon lineages (Ty1/Copia and Ty3) in MbTMV. **b** Age-related abundance of Tekay elements on MbTMV chromosomes.

#### Supplementary Fig. 11

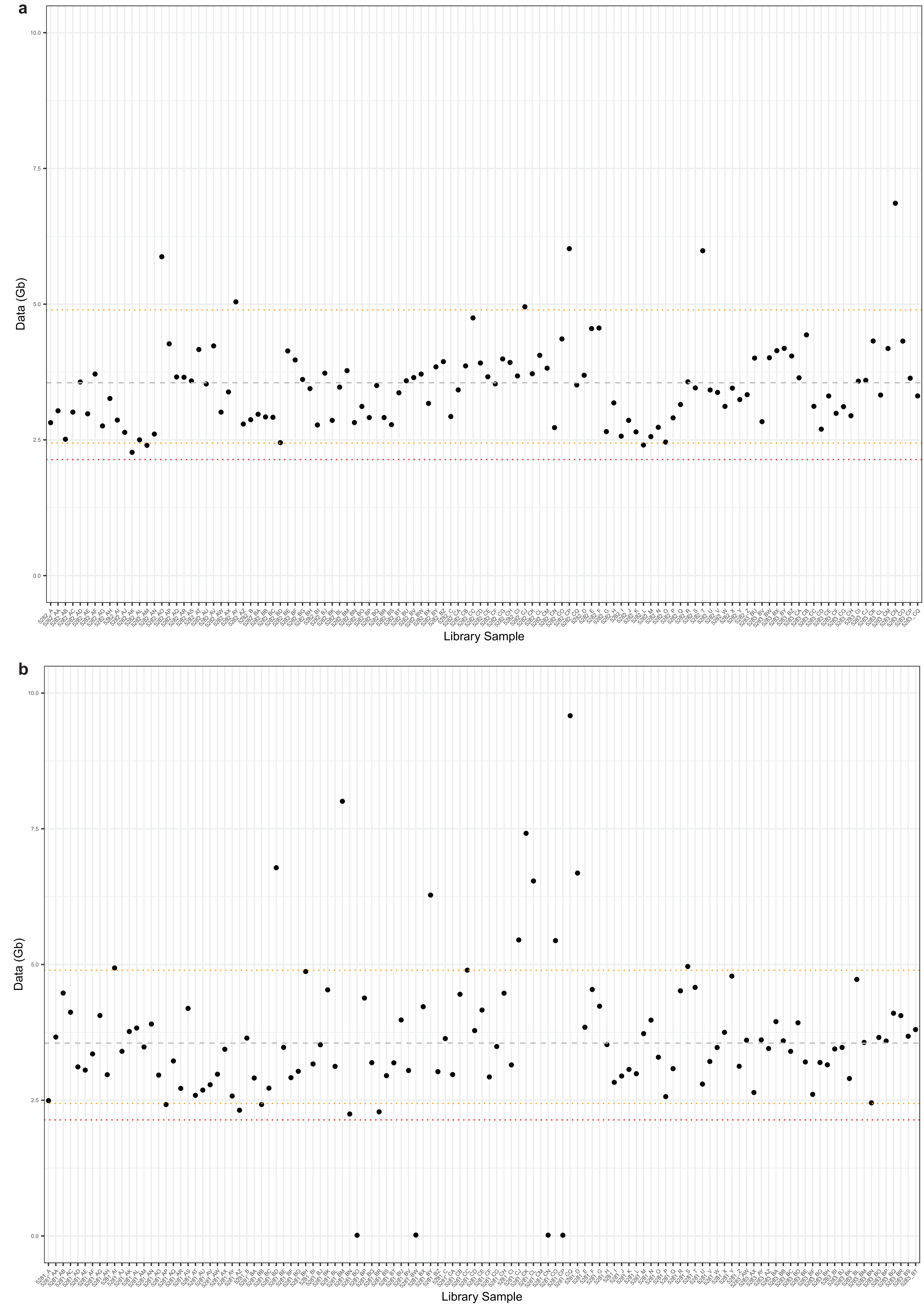


Generated sequencing data per Moneyberg-TMV x Micro-Tom F1 (MbTMV-MT) derived backcrossed individuals. **a** Female backcross samples, showing total data (Gb) generated per individual. **b** Male backcross samples, showing total data (Gb) generated per individual. Grey dashed line represent mean data. Yellow dashed lines represent 5% and 95% quantiles. Red dashed lines represent 2.5% and 97.5% percentiles. 4 Male backcross samples failed during sequencing.

### Supplementary Fig. 12

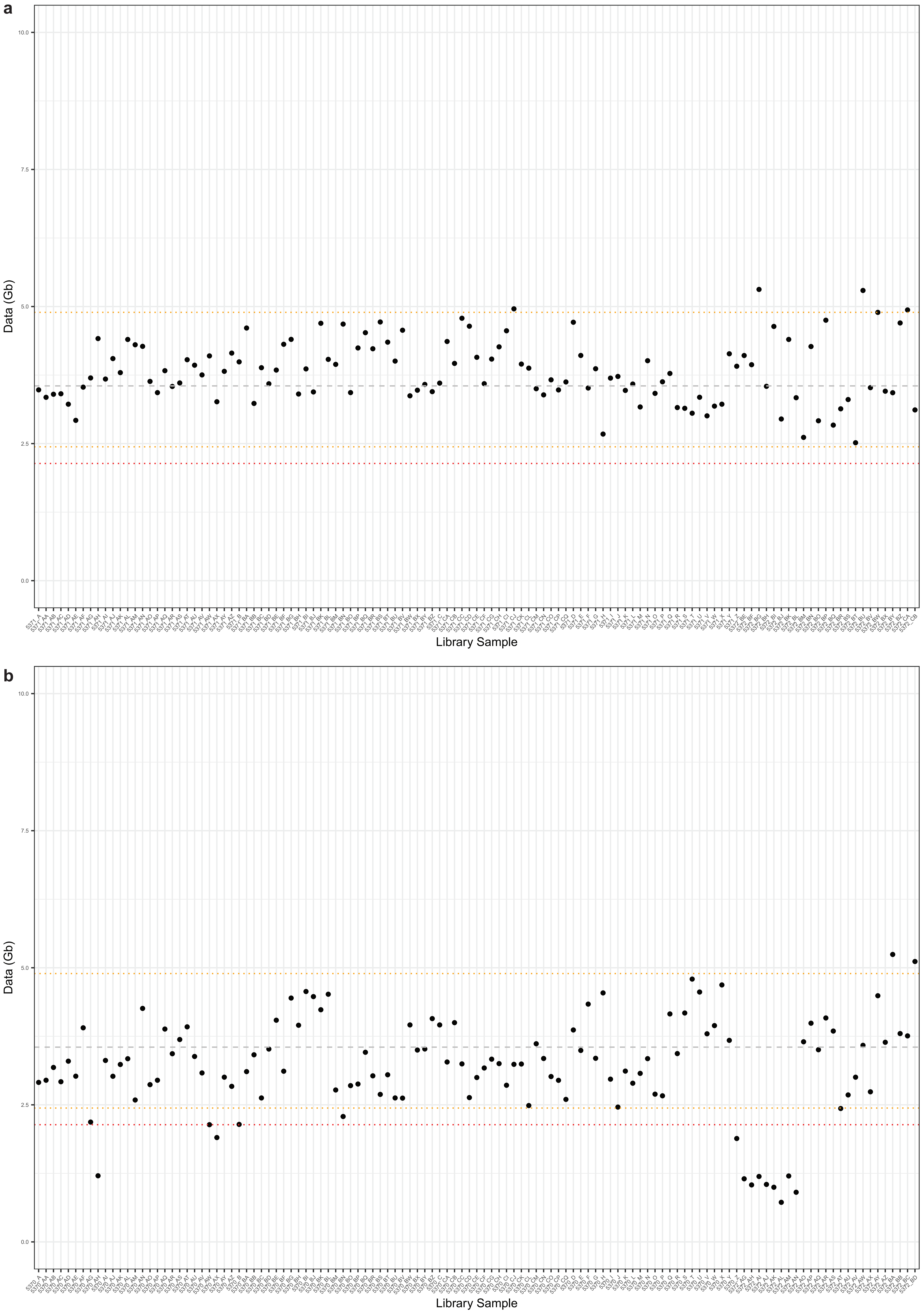


Generated sequencing data per Moneyberg-TMV x LA1039 F1 (MbTMV-S.che) derived backcrossed individuals. **a** Female backcross samples, showing total data (Gb) generated per individual. **b** Male backcross samples, showing total data (Gb) generated per individual.Grey dashed line represent mean data. Yellow dashed lines represent 5% and 95% quantiles. Red dashed lines represent 2.5% and 97.5% percentiles.

#### Supplementary Fig. 13

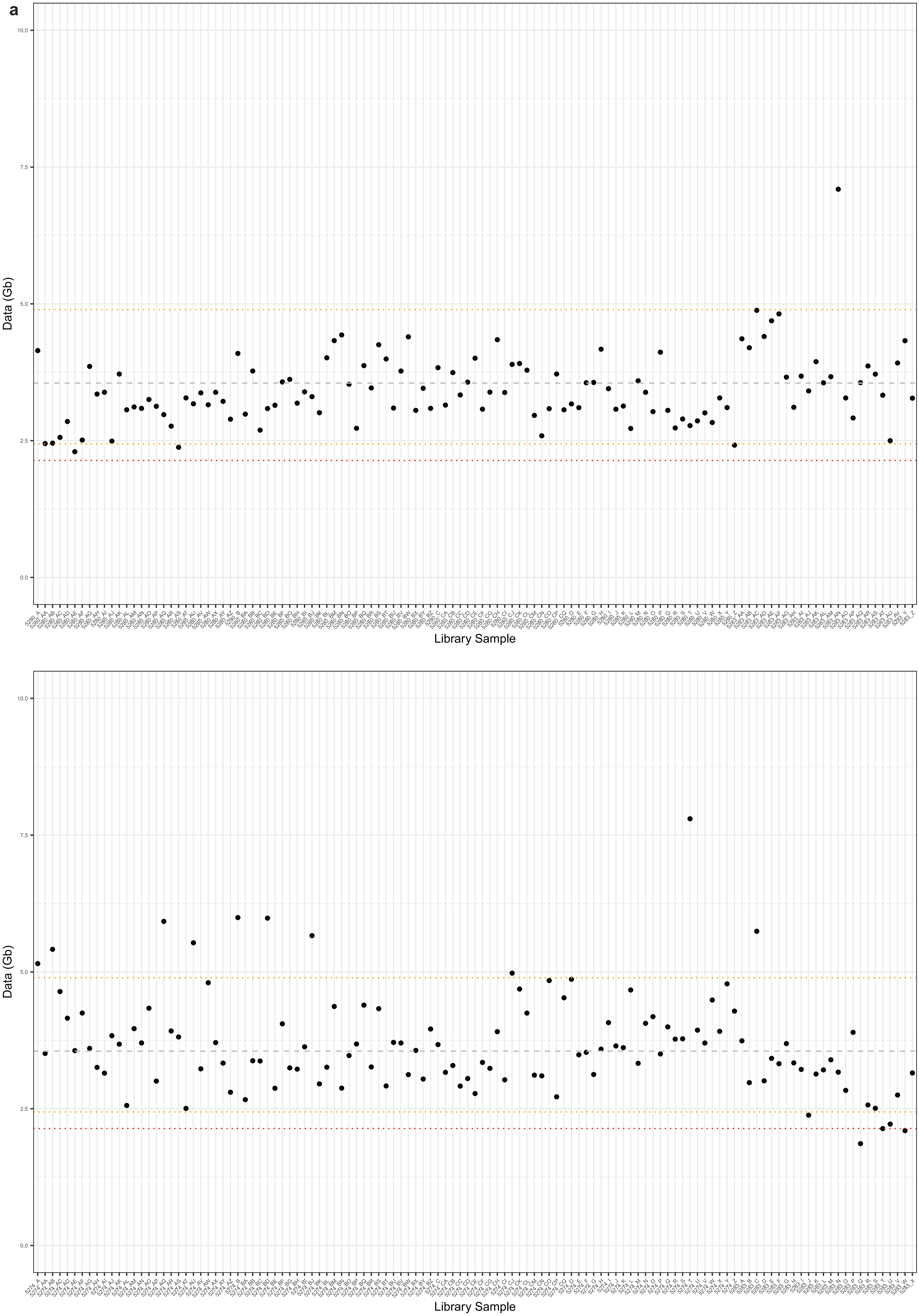


Generated sequencing data per Moneyberg-TMV x LA0716 F1 (MbTMV-S.pen) derived backcrossed individuals. **a** Female backcross samples, showing total data (Gb) generated per individual. **b** Male backcross samples, showing total data (Gb) generated per individual.Grey dashed line represent mean data. Yellow dashed lines represent 5% and 95% quantiles. Red dashed lines represent 2.5% and 97.5% percentiles.

### Supplementary Fig. 14

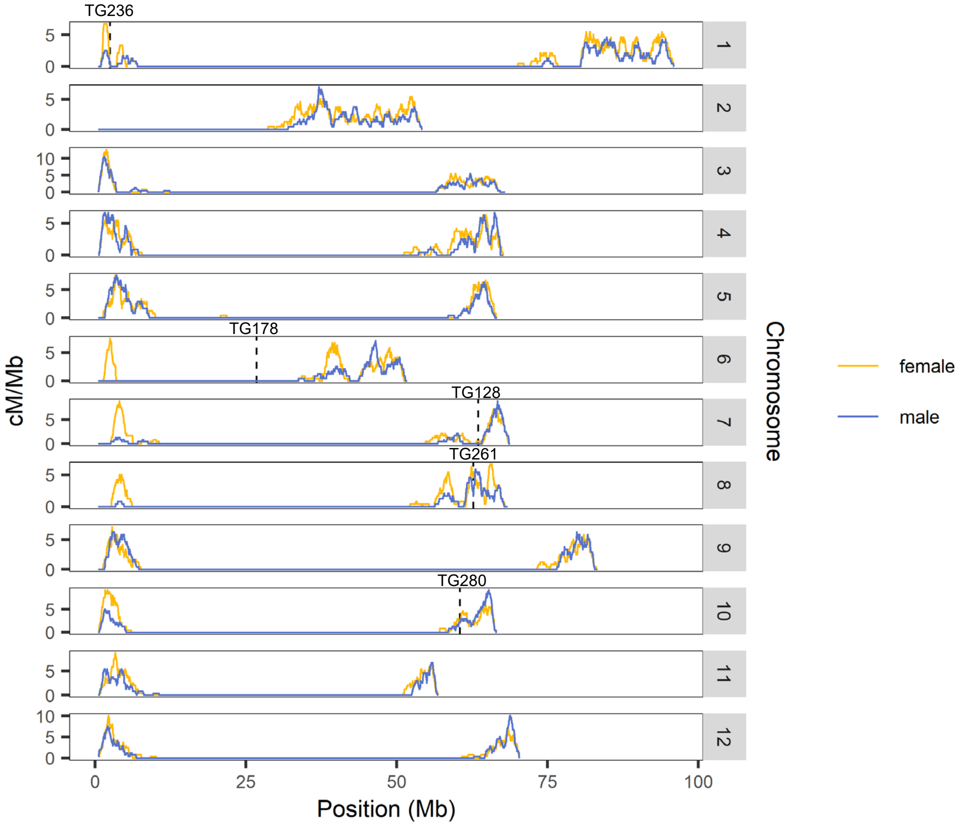


Female-specific recombination regions. Recombination landscape in male and female Mb-Pen backcross populations. The markers are based on the genetic maps constructed by de Vicente and Tanksley (1991). These markers are located upstream of the regions showing difference in genetic distance between male and female gametes, confirming the female-specific recombination regions in our backcross populations.

### Supplementary Fig. 15

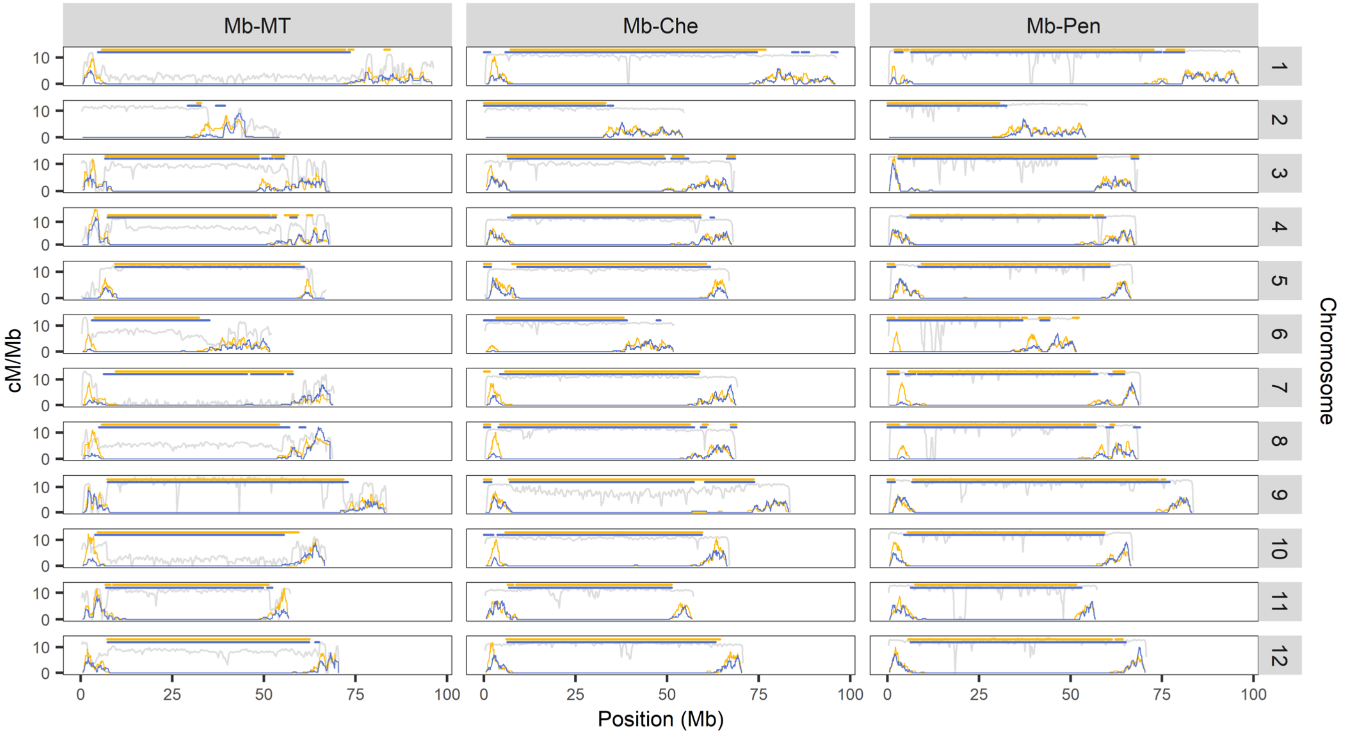


Recombination coldspots. Female (yellow) and male (blue) recombination landscape were plotted together with SNP distribution (gray). The top horizontal lines indicate the female and male recombination coldspots, which are mostly located in the pericentromeres.

### Supplementary Fig. 16

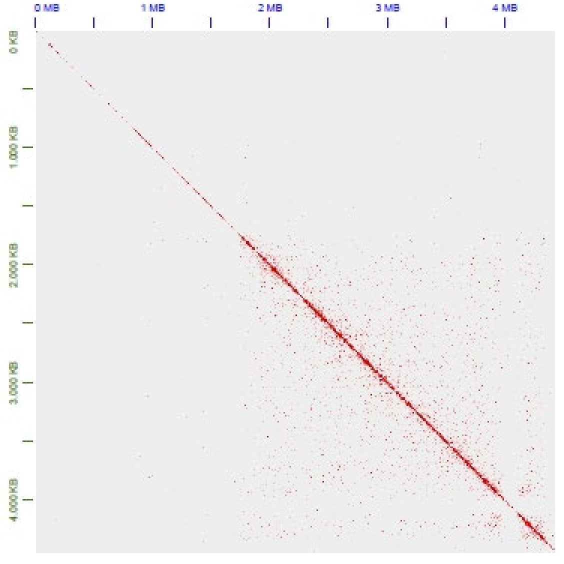


Inversion in chromosome 8. Hi-C map for an inversion causing crossover suppression between *MbTMV* and *S. pennellii*. The clean map confirms that the inversion is correct and not an assembly error or scaffolding artefact.

### Supplementary Fig. 17

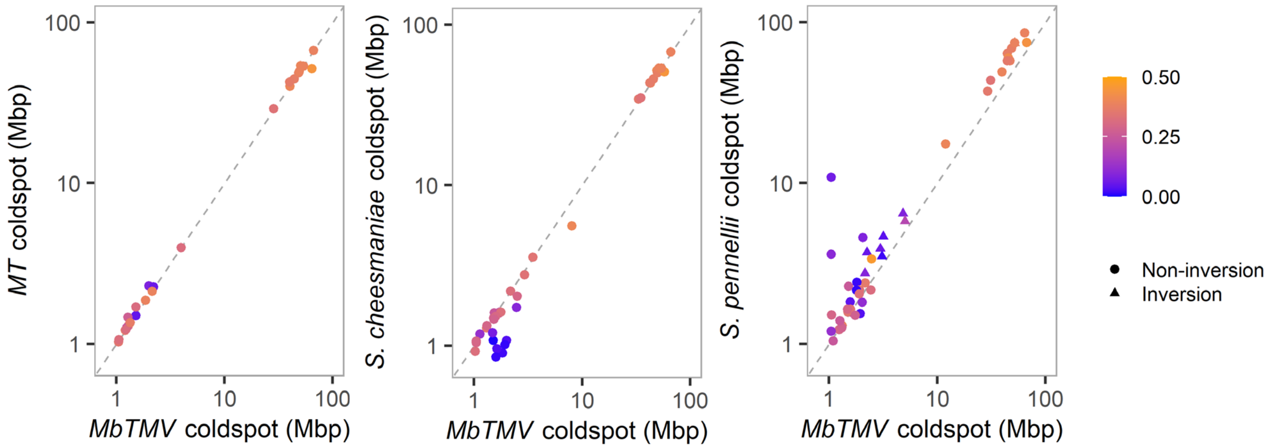


Coldspot comparison between parental genomes. Lengths of female coldspot regions relative to each parent. Deviation from the diagonal line indicates expansion or contraction of genomic segments, while the color indicates the total proportion of *Gyspy* and *Copia* elements relative to the MbTMV genome. We define inversion-associated coldspot as having more than 50% inversion coverage. Orange coldspots in the upper right of each plot are located in the pericentromeres.

### Supplementary Fig. 18

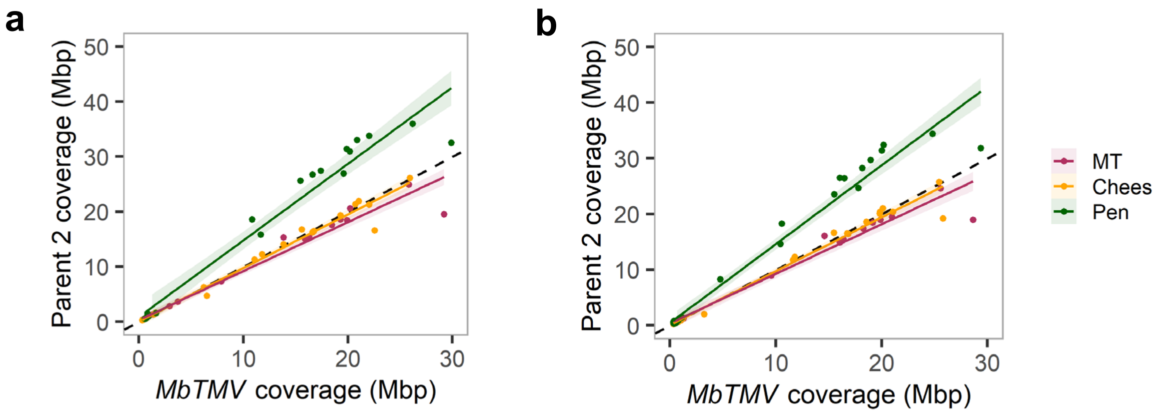


Retrotransposon copy changes in coldspots. Differential retrotransposon (i.e. Gypsy and Copia) content in **a** male and **b** female coldspot regions. Only coldspots with at least 25% retrotransposon coverage in MbTMV genome are included. Parent 2 is the parent crossed with MbTMV, differentiated by color. The broken diagonal line represents the equal parental copies of retrotransposons in the coldspot regions. Points above the diagonal indicate higher retrotransposon content in parent 2.

#### Supplementary Fig. 19

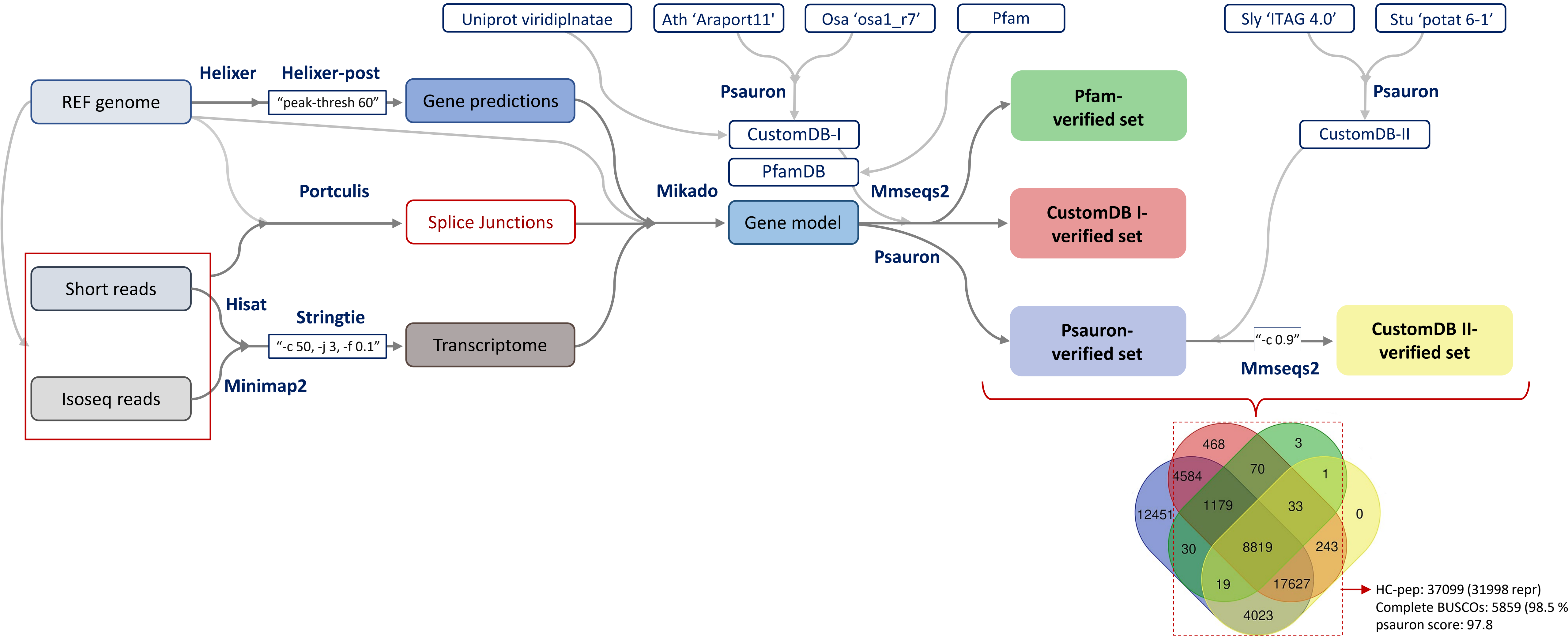


Pipeline used to develop the high-confidence gene model for LA0716. Structural gene annotation for LA0716 was conducted using an integrated pipeline combining ab-initio predictions (using Helixer v0.3.3, land_plant_v0.3_a_0080.h5) and evidence-based approaches using both short- and long-read RNA datasets (validated for splice junctions using Portcullis v1.2.4, aligned respectively using Hisat2 v2.2.1 and minimap2 v2.28, and assembled to the transcriptome using StringTie v2.2.3 ) via Mikado v2.3.3 to generate an initial consensus gene model, followed by a multi-layered refinement to minimize the potential false positives. Tool-specific settings are indicated in blue-bordered boxes within the schematic. Validation reference databases used for refinement are indicated in rounded blue-bordered boxes.
