## Supplementary Tables 1-9 for "Chromosome-scale *Solanum pennellii* and *Solanum cheesmaniae* genome assemblies reveal structural variants, repeat content and recombination barriers of the tomato clade"

### Supplementary Table 1: Overview of the publicly available genomes used for pairwise genomic comparison and TE annotation in this study

| Species | Accession | Genome Size (Mbp) | Accession number or Link | Citation |
|---|---|---|---|---|
| *S. lycopersicum* | Moneyberg-TMV | 824.45 | PRJEB44956 | van Rengs et al., 2022 |
| *S. lycopersicum* | Micro-Tom | 812.44 | https://datadryad.org/dataset/doi:10.5061/dryad.h9w0vt4qd | Wang et al., 2024 |
| *S. lycopersicum* | Heinz1706 | 800.12 | http://solomics.agis.org.cn/tomato/ftp/ | Zhou et al., 2022 |
| *S. lycopersicum* | M82 | 801.96 | https://github.com/pan-sol/pan-sol-data | Alonge et al., 2022 |
| *S. lycopersicum* | Sweet-100 | 805.19 | https://github.com/pan-sol/pan-sol-data | Alonge et al., 2022 |
| *S. galapagense* | LA0317 | 811.44 | PRJCA008297 | Yu et al., 2022 |
| *S. habrochaites* | LA0407 | 893.57 | PRJCA008297 | Yu et al., 2022 |
| *S. tuberosum* | DM8.1 | 737.87 | PRJCA011810 | Yang et al., 2022 |

### Supplementary Table 2: Statistics of PacBio HiFi sequencing data of *S. pennellii* LA0716 and *S. cheesmaniae* LA1039

Supplementary Table 2 shows the summary statistics of the PacBio HiFi sequencing datasets as calculated by Seqkit stats.

| Accession | Library | Number of sequences | Total length | Minimum length | Mean length | Maximum length | Q1 | Q2 | Q3 | sum_gap | N50 | Q20(%) | Q30(%) |
|---|---|---|---|---|---|---|---|---|---|---|---|---|---|
| **LA0716** | 5320.A | 1,556,810 | 29,916,414,922 | 86 | 19,216.50 | 44,530 | 17,981 | 19,063 | 20,335 | 0 | 19,229 | 98.46 | 96.52 |
| **LA0716** | 5405.A | 747,762 | 19,074,207,596 | 86 | 25,508.40 | 59,929 | 22,606 | 24,971 | 27,901 | 0 | 25,605 | 97.83 | 94.83 |
| **LA1039** | 5169.A | 1,496,892 | 29,661,755,807 | 48 | 19,815.60 | 43,566 | 18,566 | 19,669 | 20,969 | 0 | 19,835 | 97.7 | 94.56 |
| **LA1039** | 5405.B | 804,996 | 21,753,299,665 | 60 | 27,022.90 | 65,845 | 23,756 | 26,282 | 29,605 | 0 | 27,021 | 97.84 | 94.86 |

**Supplementary Table 3: Statistics of *S. pennellii* LA0716 ONT sequencing data**

Supplementary Table 3 shows the summary statistics of LA0716 ONT sequencing data as calculated by Seqkit stats.

| Accession | Cell | Number of sequences | Total length | Minimum length | Mean length | Maximum length | Q1 | Q2 | Q3 | sum_gap | N50 | Q20(%) | Q30(%) |
|---|---|---|---|---|---|---|---|---|---|---|---|---|---|
| LA0716 | Cell-1 | 6,248,270 | 27,654,234,087 | 2 | 4,425.90 | 319,192 | 540 | 1,318 | 3,479 | 0 | 17,686 | 82.24 | 70.77 |
| LA0716 | Cell-2 | 2,313,772 | 32,768,771,121 | 12 | 14,162.50 | 330,845 | 2,646 | 7,897 | 19,680 | 0 | 29,048 | 83.59 | 72.78 |
| LA0716 | Cell-3 | 2,005,259 | 22,595,261,204 | 1 | 11,268 | 282,104 | 1,571 | 5,749 | 15,039 | 0 | 24,967 | 86.5 | 76.75 |
| LA0716 | Cell-4 | 2,024,934 | 38,641,029,062 | 2 | 19,082.60 | 469,512 | 2,892 | 10,219 | 28,169 | 0 | 40,869 | 86.24 | 76.8 |
| LA0716 | Cell-5 | 2,619,041 | 43,127,741,818 | 4 | 16,467 | 386,073 | 1,745 | 7,298 | 23,893 | 0 | 39,607 | 86.49 | 77.16 |
| LA0716 | Cell-6 | 4,612,039 | 45,443,539,791 | 6 | 9,853.20 | 192,827 | 1,881 | 5,746 | 13,686 | 0 | 19,524 | 88.66 | 80.14 |
| LA0716 | Cell-7 | 2,171,619 | 36,943,275,247 | 11 | 17,011.90 | 368,751 | 1,825 | 8,470 | 25,580 | 0 | 37,745 | 88.37 | 79.74 |
| LA0716 | Cell-8 | 4,928,969 | 47,906,912,861 | 2 | 9,719.50 | 307,607 | 611 | 2,465 | 10,934 | 0 | 32,115 | 87.14 | 77.68 |
| LA0716 | Cell-9 | 5,733,819 | 37,511,690,014 | 1 | 6,542.20 | 289,982 | 400 | 1,218 | 5,312 | 0 | 28,469 | 89.46 | 81.33 |
| LA0716 | Cell-10 | 4,776,240 | 48,591,088,496 | 5 | 10,173.50 | 506,806 | 537 | 1,915 | 11,402 | 0 | 36,184 | 86.89 | 77.29 |

**Supplementary Table 4: Statistics of Dovetail Omni-C sequencing data of *S. pennellii* LA0716 and *S. cheesmaniae* LA1039**

Supplementary Table 4 shows the summary statistics of LA1039 and LA0716 Omni-C (Dovetail) data as calculated by Seqkit stats.

| Accession | Library | Number of sequences | Total length | Minimum length | Mean length | Maximum length | Q1 | Q2 | Q3 | sum_gap | N50 | Q20(%) | Q30(%) |
|---|---|---|---|---|---|---|---|---|---|---|---|---|---|
| LA0716 | 5319.D | 186,721,134 | 28,013,833,976 | 15 | 150.00 | 151 | 150 | 150 | 150 | 0 | 150 | 92.96 | 84.54 |
| LA0716 | 5319.D | 186,721,134 | 28,015,711,809 | 15 | 150 | 151 | 150 | 150 | 150 | 0 | 150 | 86.43 | 74.67 |
| LA1039 | 5319.A | 136,409,871 | 20,469,642,382 | 15 | 150.10 | 151 | 150 | 150 | 150 | 0 | 150 | 93.64 | 85.31 |
| LA1039 | 5319.A | 136,409,871 | 20,470,612,090 | 15 | 150.1 | 151 | 150 | 150 | 150 | 0 | 150 | 86.66 | 74.75 |

**Supplementary Table 5: Statistics of Hifiasm assemblies of *S. cheesmaniae* LA1039 and *S. pennellii* LA0716**

Supplementary Table 5 shows the summary statistics of LA1039 HiFI and LA0716 HiFi + ONT (>q90 >90kb) assemblies assembled using Hifiasm.

| Accession | LA1039 | LA0716 |
|---|---|---|
| **Number of contigs** | 836 | 423 |
| **Number of contigs (>50kb)** | 385 | 358 |
| **Cumulative size (Mbp)** | 862.66 | 1109.07 |
| **N50 (Mbp)** | 27.9 | 26.2 |
| **N90 (Mbp)** | 7.2 | 11 |
| **L50** | 11 | 14 |
| **L90** | 32 | 38 |
| **Longest contig (Mbp)** | 54 | 56.3 |

**Supplementary Table 6: Comparison of *S. pennellii* LA0716 chromosome lengths**

Supplementary Table 6 shows chromosome lengths (in Mbp) of each *S. pennellii* LA0716 chromosome assembled in this study and by Bolger *et al.* (2014).

| Chromosome | LA0716, this study | LA0716 Bolger et al., 2014 |
|---|---|---|
| ch01 | 123.5 | 109.3 |
| ch02 | 64.9 | 59.8 |
| ch03 | 90.2 | 75.4 |
| ch04 | 91.2 | 77.2 |
| ch05 | 84.7 | 78.0 |
| ch06 | 72.3 | 60.7 |
| ch07 | 89.8 | 79.3 |
| ch08 | 85.0 | 70.5 |
| ch09 | 105.2 | 84.1 |
| ch10 | 94.1 | 82.5 |
| ch11 | 72.4 | 66.2 |
| ch12 | 99.1 | 83.3 |
| Non-placed | 36.7 | 63.1 |

**Supplementary Table 7: SNP-density (per kb) in *S. lycopersicum* cv. Micro-Tom, *S. cheesmaniae* LA1039 and *S. pennellii* LA0716 genomes, compared to *S. lycopersicum* cv. Moneyberg-TMV**

Supplementary Table 7 shows the SNP density per kilobase derived from alignment of Micro-Tom, LA1039 and LA0716 PacBio HiFi data against the Moneyberg-TMV genome. SNP densities are shown per chromosome and whole genome.

| | Micro-Tom | LA1039 | LA0716 |
|---|---|---|---|
| **Chromosome** | **HiFi** | **HiFi** | **HiFi** |
| **ch01** | 0.28 | 4.21 | 22.83 |
| **ch02** | 4.10 | 5.00 | 22.39 |
| **ch03** | 1.74 | 3.82 | 22.44 |
| **ch04** | 1.79 | 5.73 | 23.67 |
| **ch05** | 8.37 | 4.76 | 25.34 |
| **ch06** | 0.96 | 5.08 | 21.79 |
| **ch07** | 0.52 | 5.11 | 24.53 |
| **ch08** | 0.22 | 6.13 | 23.20 |
| **ch09** | 15.93 | 16.96 | 23.84 |
| **ch10** | 0.18 | 4.40 | 24.15 |
| **ch11** | 4.26 | 6.93 | 21.07 |
| **ch12** | 1.12 | 6.79 | 24.49 |
| **Genome** | 3.29 | 6.24 | 23.31 |

**Supplementary Table 8: List of the public short-read RNA-seq data used in the current study for evidence-based gene predictions for *S. lycopersicum* cv. Moneyberg-TMV (MbTMV) and *S. pennellii* LA0716 genomes.**

| Sample ID | Species |
|---|---|
| SRR25660662 | Solanum lycopersicum |
| SRR25660663 | Solanum lycopersicum |
| SRR25660664 | Solanum lycopersicum |
| SRR25660665 | Solanum lycopersicum |
| SRR25660668 | Solanum lycopersicum |
| SRR25660679 | Solanum lycopersicum |
| SRR25660680 | Solanum lycopersicum |
| SRR25660666 | Solanum lycopersicum |
| SRR24864451 | Solanum lycopersicum |
| SRR24864452 | Solanum lycopersicum |
| SRR24864453 | Solanum lycopersicum |
| SRR24864457 | Solanum lycopersicum |
| SRR24864458 | Solanum lycopersicum |
| SRR24864459 | Solanum lycopersicum |
| SRR24864460 | Solanum lycopersicum |
| SRR24864461 | Solanum lycopersicum |
| SRR24864462 | Solanum lycopersicum |
| SRR24864463 | Solanum lycopersicum |
| SRR24864464 | Solanum lycopersicum |
| SRR24864465 | Solanum lycopersicum |
| SRR24864469 | Solanum lycopersicum |
| SRR24864470 | Solanum lycopersicum |
| SRR24864471 | Solanum lycopersicum |
| SRR24864474 | Solanum lycopersicum |
| SRR24864475 | Solanum lycopersicum |

| | |
|---|---|
| ERR3311252 | Solanum pennellii |
| ERR3311249 | Solanum pennellii |
| ERR3311250 | Solanum pennellii |
| ERR3311251 | Solanum pennellii |
| ERR3311253 | Solanum pennellii |
| ERR3311254 | Solanum pennellii |
| SRR27903361 | Solanum pennellii |
| SRR27903365 | Solanum pennellii |
| SRR27903360 | Solanum pennellii |
| SRR27903363 | Solanum pennellii |
| SRR27903362 | Solanum pennellii |
| SRR27903364 | Solanum pennellii |
| ERR2576914 | Solanum pennellii |
| ERR2576913 | Solanum pennellii |
| ERR2576918 | Solanum pennellii |
| SRR21505619 | Solanum pennellii |
| SRR21505620 | Solanum pennellii |
| SRR21505626 | Solanum pennellii |
| SRR21505623 | Solanum pennellii |
| SRR21505624 | Solanum pennellii |
| SRR21505625 | Solanum pennellii |
| SRR21505627 | Solanum pennellii |
| SRR21505622 | Solanum pennellii |
| SRR18392547 | Solanum pennellii |
| SRR18392543 | Solanum pennellii |
| SRR18392545 | Solanum pennellii |
| SRR18392551 | Solanum pennellii |
| SRR18392559 | Solanum pennellii |
| SRR18392548 | Solanum pennellii |
| SRR18392560 | Solanum pennellii |

| | |
|---|---|
| SRR18392561 | Solanum pennellii |
| SRR18392550 | Solanum pennellii |
| SRR18392556 | Solanum pennellii |
| SRR18392557 | Solanum pennellii |
| SRR18392562 | Solanum pennellii |
| SRR18392544 | Solanum pennellii |
| SRR18392546 | Solanum pennellii |
| SRR18392549 | Solanum pennellii |
| SRR18392552 | Solanum pennellii |
| SRR18392553 | Solanum pennellii |
| SRR18392554 | Solanum pennellii |
| SRR18392555 | Solanum pennellii |
| SRR18392558 | Solanum pennellii |
| ERR2576913 | Solanum pennellii |
| ERR2576914 | Solanum pennellii |
| ERR2576918 | Solanum pennellii |
| ERR3311249 | Solanum pennellii |
| ERR3311250 | Solanum pennellii |
| ERR3311251 | Solanum pennellii |
| ERR3311252 | Solanum pennellii |
| ERR3311253 | Solanum pennellii |
| ERR3311254 | Solanum pennellii |
| SRR18392543 | Solanum pennellii |
| SRR18392544 | Solanum pennellii |
| SRR18392545 | Solanum pennellii |
| SRR18392546 | Solanum pennellii |
| SRR18392547 | Solanum pennellii |
| SRR18392548 | Solanum pennellii |
| SRR18392549 | Solanum pennellii |
| SRR18392550 | Solanum pennellii |

| | |
|---|---|
| SRR18392551 | Solanum pennellii |
| SRR18392552 | Solanum pennellii |
| SRR18392553 | Solanum pennellii |
| SRR18392554 | Solanum pennellii |
| SRR18392555 | Solanum pennellii |
| SRR18392556 | Solanum pennellii |
| SRR18392557 | Solanum pennellii |
| SRR18392558 | Solanum pennellii |
| SRR18392559 | Solanum pennellii |
| SRR18392560 | Solanum pennellii |
| SRR18392561 | Solanum pennellii |
| SRR18392562 | Solanum pennellii |
| SRR21505619 | Solanum pennellii |
| SRR21505620 | Solanum pennellii |
| SRR21505622 | Solanum pennellii |
| SRR21505623 | Solanum pennellii |
| SRR21505624 | Solanum pennellii |
| SRR21505625 | Solanum pennellii |
| SRR21505626 | Solanum pennellii |
| SRR21505627 | Solanum pennellii |
| SRR27903360 | Solanum pennellii |
| SRR27903361 | Solanum pennellii |
| SRR27903362 | Solanum pennellii |
| SRR27903363 | Solanum pennellii |
| SRR27903364 | Solanum pennellii |
| SRR27903365 | Solanum pennellii |

**Supplementary Table 9: Statistics of PacBio Isoseq reads used in the current study for evidence-based gene predictions for *S. lycopersicum* cv. Moneyberg-TMV (MbTMV) and *S. pennellii* LA0716 genomes.**

Supplementary Table 9 shows the summary statistics of LA0716 and MbTMV PacBio Isoseq data as calculated by Seqkit stats.

| Species | Library | Number of sequences | Total length | Minimum length | Mean length | Maximum length | Q1 | Q2 | Q3 | sum_gap | N50 | Q20(%) | Q30(%) |
|---|---|---|---|---|---|---|---|---|---|---|---|---|---|
| Moneyberg-TMV | roots-anthers-leaf | 4,705,644 | 5,123,548,961 | 51 | 1,088.80 | 7,059 | 690 | 937 | 1,375 | 0 | 1,230 | 99.6 | 99.12 |
| LA0716_ pool 1 | fruit-root-leaf-stem | 11,485,181 | 44,688,801,326 | 44 | 3,891 | 25,242 | 2,952 | 3,669 | 4,599 | 0 | 4,061 | 99.36 | 98.54 |
| LA0716_ pool 2 | flower/bud-leaf-root | 3,688,941 | 9,402,439,437 | 77 | 2,548.80 | 13,039 | 1,931 | 2,433 | 3,018 | 0 | 2,719 | 99.19 | 98.1 |
